# Elucidation of α-glucosidase inhibitory activity and UHPLC-ESI-QTOF-MS based metabolic profiling of endophytic fungi *Alternaria* sp. BRN05. isolated from seeds of *Swietenia macrophylla* King

**DOI:** 10.1101/2024.09.25.615012

**Authors:** Piyush Kumar, Sai Anand Kannakazhi Kantari, Ranendra Pratap Biswal, Prasanth Ghanta, Malleswara Dharanikota

## Abstract

There is a growing demand for new diabetes drugs with fewer side effects to replace current medications known for their adverse effects. Inhibition of α-glucosidase responsible for postprandial hyperglycaemia among diabetes patients is a promising strategy for managing the disease. This study aims to explore and identify novel bioactive molecules with anti-diabetes potential from *Alternaria* sp. BRN05, an endophytic fungus isolated from a well-known medicinal plant *Swietenia macrophylla* King. Ethyl acetate extracts of *Alternaria* sp. BRN05 grown in full-strength (EFS) and quarter-strength (EQS) media respectively were evaluated for their α-glucosidase inhibitory activities. Based on IC_50_ values, EQS exhibited significantly greater inhibitory activity (0.01482 ± 1.809 mg/mL) as compared to EFS (1.16 ± 0.173 mg/mL) as well as acarbose control (0.494 ± 0.009 mg/mL). EFS and EQS were subjected to metabolic profiling using Ultra-High-Performance Liquid Chromatography - Electrospray Ionization - Quadrupole Time-of-Flight Mass Spectrometry (UHPLC-ESI-QTOF-MS). A total of nineteen molecules from EFS and twenty from EQS were tentatively identified based on MS/MS fragmentation. Molecular docking analysis revealed that twelve among these exhibited greater binding energies than that of acarbose (-6.6 kcal/mol). Two molecules from EQS, 3’,4’,7-trihydroxyisoflavanone (-7.5 kcal/mol) and alternariol 9-methyl ether (-7 kcal/mol), were subjected to ensemble docking studies, which further strengthened their promise as good α-glucosidase inhibitors for treating diabetes.

## 1. Introduction

*Swietenia macrophylla* King, belonging to family Meliaceae, is a medicinal plant traditionally used for treating various ailments. This tree grows in 40 countries, including South America, Brazil, Indonesia, India, Sri Lanka, Malaysia, etc. Extracts prepared from its leaves and seeds are used in ayurveda, siddha and folk medicine. Species of genus *Swietenia* are rich in phytochemicals like limonoid and polyphenolics known to exhibit medicinal properties. Some of the bioactive metabolites such as swietemacrophia and swietenolide possess good anti-inflammatory and antibacterial activities(Da Silva et al., 2008).

In India, decoction prepared from leaves and bark is traditionally used in the treatment of nerve ailments and diarrhoea, while fruit extracts are used in the treatment of hypertension and skin-related problems. In Malaysia and Indonesia, raw or crushed seeds of *Swietenia* are used in the treatment of hypertension, malaria, and diabetes. They also possess good anti-inflammatory, anti-mutagenic, and anti-tumour activities. During the last decade, *Swietenia* has gained prominence in the treatment of diabetes(Sukardiman and Ervina, 2020).

Genus *Alternaria* belonging to family Pleosporaceae is a fungus distributed widely across the world. It is a free-living form that grows well in the soil as well as inside plant tissues as an endophyte. It produces a diverse variety of bioactive metabolites. In the last twenty year at least 268 metabolites exhibiting good bioactivity have been reported from different species of *Alternaria*(Sabreen et al., 2015).

Bioactive metabolites with diverse medicinal uses are reported from endophytic species of *Alternaria* such as, saponin, phenols, steroids and glycosides possessing antioxidant properties; taxol, coumarins, paclitaxel and camptothecin showing anti-tumour properties and alternariol, alterchromone A, tannins and alternariol-9-methyl ether exhibiting antimicrobial properties (Eram et al., 2018). Porritoxin is a well-known chemo-preventive molecule with a potential for use as a drug in the treatment of cancer (Kim et al., 2009).

Many of the genes encoding secondary metabolites in endophytic fungi are usually silent or cryptic when cultured under normal conditions. Genes responsible for producing secondary metabolites normally occur in groups referred to as Biosynthetic Gene Clusters (BGCs). A fungal genome usually contains about 40-50 BGCs. Activation of a BGCs can lead to production of a set of novel compounds (Keller, 2019). Several methods can be employed for activating BGCs including co-culturing with microbes like bacteria and fungi; addition of small-molecule elicitors such as DNA methyltransferase inhibitors (e.g., 5-azacytidine) or histone deacetylase inhibitors (e.g., SAHA); addition of physical scaffolds like cotton scaffold and microparticles to the culture media; changing physical or chemical conditions such as temperature, light, salt concentration; and altering media concentration(Tomm et al., 2019). Analysis of available databases using AntiSMASH software showed that *Alternaria* harbours 22 BGCs involved in the production of secondary metabolites. Out of these BGCs, only eight have been studied to some extent, while information for the remaining fourteen is not available (Tao et al., 2022).

According to the World Health Organization (WHO) report of 2016, an estimated 422 million individuals worldwide were affected by diabetes mellitus. International Diabetes Federation (IDF) in 2017 reported that a staggering 72.9 million people in India are grappling with diabetes mellitus (Cho et al., 2018). This covers both Type 1 and Type 2 diabetes, with Type 2 diabetes being the more prevalent form accounting for approximately 90% of diabetes cases. It is well-documented in the literature that this number is expected to surge to 592 million by 2035(insight review articles 782, 2001).

Diabetes is characterized by elevated blood sugar levels (hyperglycemia). This can lead to a host of complications, like cardiovascular diseases, renal failure, neuropathy, retinopathy and disorders of lipid metabolism(Jiao et al., 2018). The key enzyme responsible for breaking down complex carbohydrates into absorbable monosaccharides is α-glucosidase, found in the brush border of the small intestine. In diabetic patients it significantly contributes to elevated blood sugar levels(Jaiswal, 2013). Addressing hyperglycemia is a key requirement in the management of diabetes and the most promising approach probably is inhibition of α-glucosidase activity. While several commercial drugs like acarbose, voglibose, and miglitol are readily available and widely used for the treatment of postprandial hyperglycaemia, they are often associated with numerous side effects, including diarrhoea, allergic reactions, stomach pain and flatulence (Patil et al., 2015). This underscores the need for developing alternative drugs which can effectively inhibit α-glucosidase and at the same time produce minimal or no side effects. In recent times, use of α-glucosidase inhibitors from natural sources has emerged as a highly effective strategy for managing postprandial hyperglycemia and its associated physiological complications, particularly in cases of Type 2 diabetes. One promising, yet relatively underexplored avenue is investigating endophytic fungi as sources for novel bioactive molecules.

## 2. Materials and methods

### 2.1 Chemicals and reagents

We purchased α-glucosidase derived from *Saccharomyces cerevisiae* (CAS No: 9001-42-7) from Merck Saint Louis, USA, PNPG (p-nitrophenyl-alpha-D-glucopyranoside) LOT: 3533550 was obtained from Merck Darmstadt, Germany, and Acarbose (CAS No: 56180-94-0, purity >98%) was sourced from TCI, Tokyo, Japan. Potato dextrose agar (PDA, GMO96-500G), Potato dextrose broth (PDB, GM403-G00G), Folin & Ciocalteu’s Phenol reagent Hi-LR^TM^ (RM10822-100ML), Ethyl acetate (CAS No: 141-78-6, purity 99.9%), and Methanol (CAS No: 67-56-1, purity 99.8%) of HPLC grade used for extraction were purchased from Himedia, Mumbai, India. LC-MS grade Acetonitrile (CAS No: 75-05-8, >99.9% purity) was procured from Fisher Scientific, Fair Lawn, NJ, USA. Milli Q: Type 1 water, purified using the Merck Milli Q IQ7000 water purification system.

### 2.1 Collection of samples

Seeds were collected from *Swietenia macrophylla* King located at the geographical coordinates (Latitude – 13.001549° and Longitude - 77.758875°) at Sri Sathya Sai Institute of Higher Learning, Brindavan campus, Whitefield, Kadugodi, Bengaluru, Karnataka-560067. L. Rasingam from the Botanical Survey of India (Deccan Regional Centre), Ministry of Environment, Forests & Climatic Change, Hyderabad, Telangana, India, authenticated the plant sample. The voucher number for the sample is BSI/DRC/2022-23/Identification/94.

### 2.2 Isolation of endophytic fungi

The mature fruits of *Swietenia macrophylla* King yielded approximately 35 to 40 seeds. The seeds were washed with distilled water and then cut into small pieces measuring 0.5 cm². To sterilize the seed surfaces, around 300 seed pieces were placed in a beaker and treated with 70% ethanol (50 mL) for 10 seconds, followed by a 3-minute treatment with 4% sodium hypochlorite (50 mL). After one minute, the seeds were rinsed with sterile distilled water (50 mL). Next, five sterilized seed pieces were placed on 9 mm petri plates containing 20 mL of Potato Dextrose Agar (PDA) media. The media was poured inside a laminar airflow hood at room temperature. The plated seeds were then incubated for 3-6 days until colonies of endophytic fungi became visible (refer to supplementary file 1). Fungal growth from the seeds was transferred using a loop into fresh petri plates containing PDA media to obtain pure cultures. The resulting pure fungal cultures were preserved at -20°C for future use.

### 2.3 Extracellular metabolites extraction

The pure isolates of endophytic fungi were inoculated into 100 mL full strength (FS) and Quarter strength (QS) PDB media and was cultured for 21 days. Before extraction, the fungal culture was homogenized with 10% methanol. Ethyl acetate was used as the solvent for extraction in a 1:1 ratio, and the process was repeated twice to extract maximum metabolites from the culture. Ethyl acetate was evaporated using rotary evaporator, and the resulting extract was dried and stored at -20°C for further analysis.

### 2.4 Phylogenetic Analysis of Endophytic Fungi Using ITS-rDNA Fragment

The isolated endophytic fungi were investigated by first extracting genomic DNA from their mycelium using a phenol-chloroform extraction method. Next, PCR was carried out by using specific primers (ITS1 forward primer and ITS4 reverse primer) to amplify the ITS-rDNA fragment. The resulting sequences were then compared to other sequences in the NCBI GenBank using BLAST searches. To determine the relationships between different fungal species, phylogenetic analysis was conducted using MEGA X (Kumar et al., 2018). The evolutionary distances were calculated using the maximum composite likelihood technique. The resulting tree was created using the Neighbour-Joining method. To ensure the accuracy of the tree, a process called bootstrapping was used (Tamura et al., 2004). The tree was tested 1000 times to determine the frequency with which the species grouped together. This allows us to confidently infer the evolutionary history of the endophytic fungi.

### 2.5 Total phenolic content

The total phenolic content (TPC) was determined with slight modifications from Akhtar et al. (Akhtar et al., 2018) using Folin-Ciocalteu reagents (FC reagents). Gallic acid served as the reference for constructing the standard curve. It was dissolved in methanol at concentrations ranging from 1 µg/mL to 10 µg/mL to prepare the standards. Samples of EFS and EQS were prepared at a concentration of 1 mg/mL in methanol. Thirty microliters (30 µL) of each standard solution and samples were added to the 96-well plate, followed by the addition of 10X-diluted Folin reagent (150 µL). Subsequently, 120 µL of 5% sodium carbonate solutions was added and plate was incubated in the dark at room temperature for thirty minutes. After incubation, the absorbance was measured at 765 nm using a microplate reader. The results of the TPC analysis were expressed in gallic acid equivalents per gram of dry extract weight (mg GAE/g).

### 2.6 Antidiabetic assays

The α-glucosidase inhibition assay was optimized by incorporating modification from Bhatia *et al* (Bhatia et al., 2019). Ethyl acetate (EtOAc) extracts from full strength (FS) and quarter strength (QS) PDB cultures of the nine endophytic fungi isolated from *Swietenia macrophylla* King seeds were screened for inhibitory activity of α-glucosidase. First, a 0.1 M phosphate buffer at pH 6.9 was prepared. Subsequently, α-glucosidase was dissolved in the phosphate buffer to achieve a concentration of 0.2 U/mL. Subsequently, a solution of 1 mM PNPG (p-nitrophenyl-alpha-D-glucopyranoside) was prepared in the phosphate buffer. Samples of EFS and EQS were initially prepared at concentrations of 2 mg/mL and 1 mg/mL, respectively, by dissolving them in a mixture of 10% dimethyl sulfoxide (DMSO) and phosphate buffer. Following favorable outcomes, these samples were prepared at various concentrations for IC_50_ calculation. Additionally, a 0.1 M Na_2_CO_3_ solution was prepared using MilliQ water. Acarbose, serving as the positive control, was prepared in the phosphate buffer. Moreover, 10% DMSO was utilized as the vehicle control for the experiment. In the 96-well plate, 50 µL of 0.1 M phosphate buffer at pH 6.9 was dispensed. Subsequently, 10 µL of α-glucosidase was added. Following this, 20 µL of extracts were introduced at various concentrations, ranging from 4 mg/mL to 0.6 mg/mL for FS and from 25 µg/mL to 5 µg/mL for QS. Then, 20 µL of PNPG was added, and the incubation was continued for another 20 minutes at 37°C. The reaction was terminated by the addition of 100 µL of Na_2_CO_3_. Finally, the absorbance of the reaction mixture was measured at 405 nm using a microplate reader (SpectraMax® M2e). Percentage inhibition was calculated using the following equation:

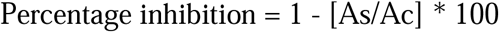

whereas As signifies the absorbance of the sample and Ac signifies the absorbance of the control. The IC_50_ value was calculated for the endophytic fungal strain in both EFS and EQS.

### 2.7 Metabolite profiling of UHPLC-ESI-QTOF-MS

The EFS and EQS was subjected to metabolite profiling using UHPLC-ESI-QTOF-MS mass spectrometry purchased from Agilent model 1290 that is coupled to Agilent 6550 Q-TOF LC-MS with dual jet stream ionization source. Agilent MassHunter version B.05.00 software (Agilent Technologies, USA) was employed for data acquisition.

Samples EFS and EQS were prepared by dissolving in methanol to achieve a final concentration of 0.1 mg/mL. Filtration of EFS and EQS was performed using an Agilent Econofilter column composed of polytetrafluoroethylene (PTFE) with a 13 mm diameter and a 0.22 µM pore size. Metabolite separation was carried out on an Agilent ZORBAX RRHD Eclipse Plus C18 column (3.0×100 mm, 1.8 µm) utilizing a gradient elution profile with mobile phase A (0.1% HCOOH in water) and mobile phase B (0.1% HCOOH in acetonitrile). The elution gradient consisted of the following time intervals: 0-2 min at 5% B, 2-21 min with a gradual transition from 5% to 20% B, 21-40 min with a progressive transition from 20% to 50% B, 40-45 min with a stepwise transition from 50% to 95% B, 45-49 min at a constant 95% B, and 49-51 min returning to 5% B. The injection volume was 10 µL, and the flow rate was maintained at 0.300 mL/min (Mohammed et al., 2021).

For the quadrupole time of flight analysis, specific parameters were set as follows: nozzle voltage of 1000 V, capillary voltage of 3.5 kV, nebulizer pressure of 35 psi, and drying gas nitrogen flow rates of 11 l/min with respective temperatures of 250 °C and 350 °C. Mass calibration was performed using the G1969-85,000 reference standard mix (Supelco, Inc.) in both positive and negative ionization modes, resulting in a minimal residual error value of 0.2 ppm (Biswal et al., 2022).

### 2.8 Library creation for metabolites database

A collection of metabolites derived from *Alternaria* sp., including known α-glucosidase inhibitors and well-known polyphenols, was gathered from existing literature. A comprehensive database comprising 850 small molecules was created using the Agilent Personal Compound Database and Library (PCDL) software (Agilent technologies, USA). The software is used to manage the content of personal compound databases and libraries and is part of the Agilent MassHunter Workstation software suite. The structures of these metabolites were either drawn using ChemDraw software (ChemBioOffice Suite, PerkinElmer Informatics, USA) or retrieved from the PubChem Database.

To analyze the data, Agilent MassHunter Qualitative Analysis software (B.7.00) was utilized. The uploaded data was screened using the "Find by formula" option, which enabled analysis within specified mass windows. This process provided match scores based on a combination of accurate mass, isotopic abundance, and isotopic spacing to identify the metabolites.

### 2.9 Molecular docking

Human Maltase-glucoamylase (MGAM) plays a crucial role in the final breakdown of starch into glucose within the small intestine. This enzyme is categorized under Glycosyl hydrolase family 31 (GH31) due to its α-glucosidase activity. MGAM comprises two subunits, NtMGAM located in the N-terminal region, and CtMGAM located in the C-terminal region. The primary function of human MGAM is to cleave α (1-4) linkages between glucosyl units, ultimately producing glucose as the final product. The goal is to assess the ligand’s inhibition capacity against NtMGAM through *in-silico* studies. For our docking studies, we focused on NtMGAM since it prefers cleaving α-1-4-bonds of shorter oligosaccharides and is involved in the final step of glucose formation. CtMGAM on the other hand prefers cleaving longer oligosaccharides (Brás et al., 2018; Elferink et al., 2020). Since standard drug acarbose weakly inhibits NtMGAM compared to CtMGAM, there is need for new drugs which target the latter (Laoud et al., 2018). NtMGAM is therefore selected as the representative of GH31 α-glucosidase family to which yeast α-glucosidase belongs to (Kashtoh and Baek, 2022). In the C-terminal region of NtMGAM, there exists a (β/α)8-barrel structure, which primarily constitutes the active site of the protein. Notable among these residues are catalytic nucleophiles like Asp443 and Asp542, which are shared across GH31 family members and play a vital role in acid-base catalysis. Additionally, residues such as His600, Asp327, and Arg526 contribute to hydrogen bonding and are also highly conserved in the GH31 family of α-glucosidase (Sim et al., 2008).

To analyze the docking studies of α-glucosidase in the GH31 family, we obtained the crystal structure of the NtMGAM protein with the PDB ID 2QMJ from the Protein Data Bank. The protein was prepared by removing heteroatoms and glycosylated residues. Any missing residues were modeled using a combination of Chimera and Modeller software. A comprehensive list of all metabolites identified in the LC-MS profiling was downloaded in 3D format as .sdf files from PUBCHEM. These metabolites were then subjected to energy minimization using HyperChem-7 and converted into PDF format. Subsequently, docking experiments were conducted for the modeled protein and ligands using AutoDock Vina, which incorporates PyMOL as a plugin (Seeliger and De Groot, 2010; Trott and Olson, 2010). To facilitate the docking procedure, we identified the active site of the protein based on literature findings. The grid dimensions were set to (x=29.184, y=29.795, z=25.403) for rigid docking. A default population size of 100 was chosen in the AutoDock program (Ghanta et al., 2022). The results of ligand binding to the protein were assessed using binding energy calculations, and their interactions were further analyzed using LigPlot.

### 2.10 Ensemble docking

The limitation posed by protein flexibility stands out as a prominent challenge in the realm of structure-based drug development. When a protein is confined to a singular configuration, essential dynamic aspects of protein-ligand binding may go unnoticed. Despite the existence of a variety of docking techniques designed to accommodate ligand flexibility, studies on protein flexibility have received limited attention until recently (Al-Nema et al., 2023). To address this issue, methodologies such as Monte Carlo and molecular dynamics simulations have been employed. The optimal approach for introducing protein flexibility involves simultaneously exploring and optimizing the complete degree of freedom for both the protein and the ligand (Huang and Zou, 2007). In the context of predicting the binding mechanism of a selected molecule into the α-glucosidase active site, obtained through clustering simulation, an ensemble docking was carried out. This comprehensive approach considers the dynamic nature of both the protein and the ligand, providing a more realistic representation of the binding process (Ghorbani et al., 2023). The protein preparation process involved several key steps to ensure the reliability of subsequent molecular dynamics simulations. Heteroatoms, including glycosylated residues, were removed from the protein structure due to issues with parametrization, and their absence from the active site. To address missing residues, modeling was performed using Chimera and Modeller, with Model 6 selected based on the best Z-DOPE score (-2.14) from the minimization log file. The resulting structure underwent further minimization using Swiss PDB Viewer with the GROMOS force field. Hydrogens were added to the structure for pH 7.4 using PDBFixer, resulting in the file named "2QMJ_rec_pdbfixed.pdb." To assess the quality of the structure, the Swiss Model Expasy server’s structure assessment tool was employed. Model refinement was conducted through the MODREFINER web server, and the refined model was submitted for assessment. As the refined model closely resembled the modeled structure, a 50 ns molecular dynamics simulation was initiated for the protein in a 1 nm^3^ periodic box, with TIP3P water and 0.15 M NaCl. Upon completion of the 50 ns simulation, clustering analysis based on the RMSD of the backbone of the pocket residues was performed using GROMACS. The trajectory was sampled from 25 ns to 50 ns, and representative structures from the top clusters were extracted. Nine representative structures were used for docking studies for the selected ligands.

## 3. Results

### 3.1 Molecular Identification

*Alternaria* sp. BRN05 (Sequence ID: SUB11758282) was identified by employing Internal transcribed spacer (ITS) sequencing method. Phylogenetic analysis of *Alternaria* sp. strain BRN05 was carried out using Neighbour-joining tree method. The sum of branch lengths for the optimal trees was calculated as 0.04229034. To analyse the tree value accurately, the test was carried out a thousand times using bootstrapping technique to ensure that the results were robust and reliable. The phylogenetic tree is given in Figure 1.

**Figure 1.**
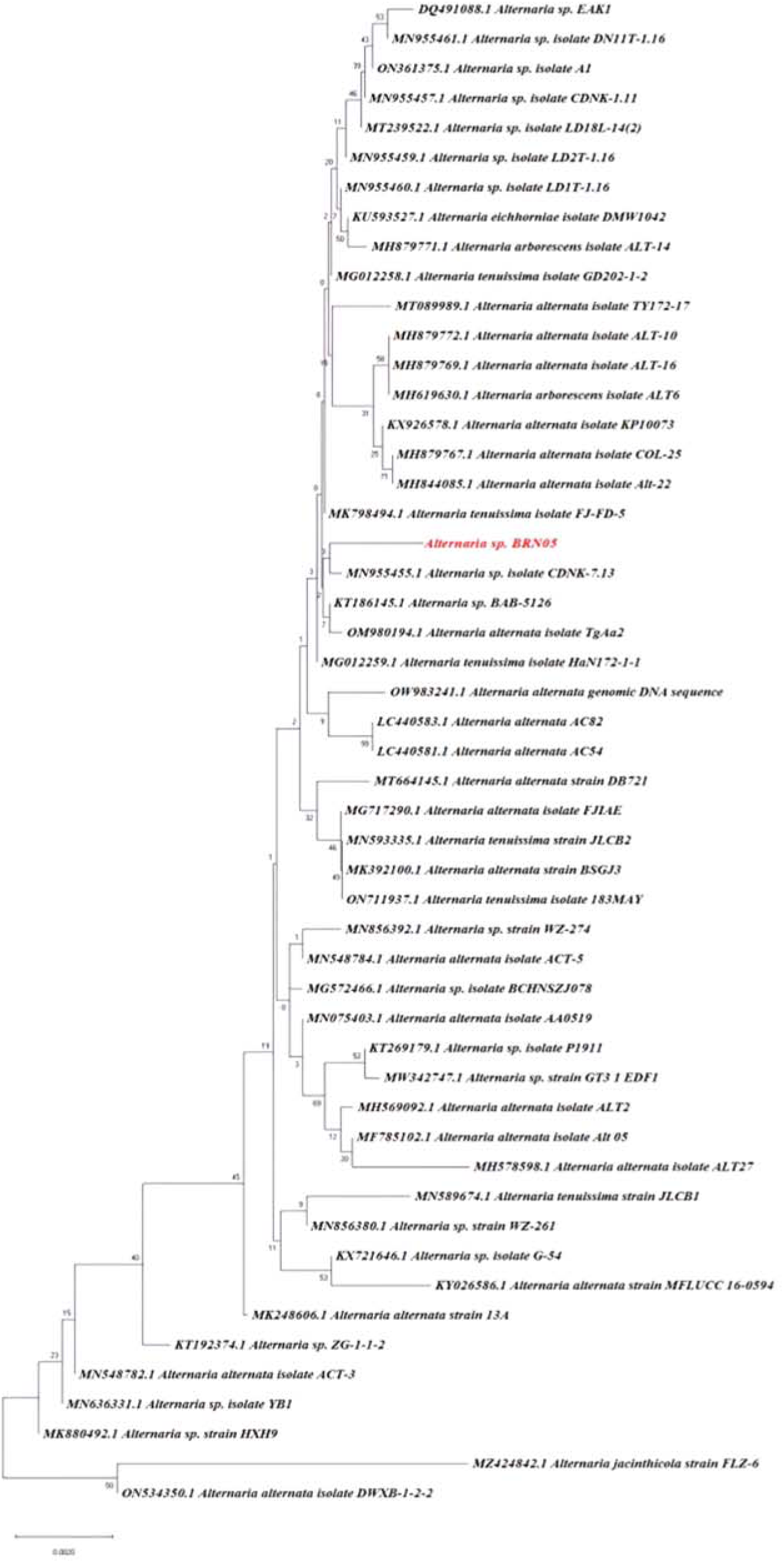
Phylogenetic tree identification of Alternaria sp. BRN05 using ITS sequencing.

### 3.1 Total phenolic content (TPC)

The total phenolic content in the samples were determined by preparing a calibration curve (Supplementary file 1 (B)) generated by plotting different concentrations of gallic acid standards ranging from 1 µg/mL to 10 µg/mL on x-axis and corresponding absorbance (OD) on y-axis. The results were expressed based on linear regression analysis (y = 0.0752x + 0.0068, R2 = 0.9983) and presented as gallic acid equivalents (GAE) per unit of dry extract weight. EQS sample exhibited a greater phenolic content (11.01 ± 0.06 mg GAE/g) as compared to EFS sample (9.22 ± 0.101 mg GAE/g) as shown in Table 1.

**Table 1.** IC_50_ values were determined for EFS and EQS against. α**-glucosidase.**

| S. No | Sample | IC<sub>50</sub> |
| --- | --- | --- |
| 1 | SMS6FS | 1.16 ± 0.173 mg/mL |
| 2 | SMS6QS | 0.01482 ± 1.809 mg/mL |

### 3.2 **α**-glucosidase Inhibition assay

The α-glucosidase inhibitory activity of EFS and EQS was carried out. EQS showed a significant IC_50_ value (0.01482 ± 1.809 mg/mL), as compared to that of EFS (1.16 ± 0.173 mg/mL) and the positive control, acarbose (0.494 ± 0.009 mg/mL) as shown in Figure 2. EQS exhibited stronger α-glucosidase inhibitory activity than that of EFS and acarbose.

**Figure 2.**
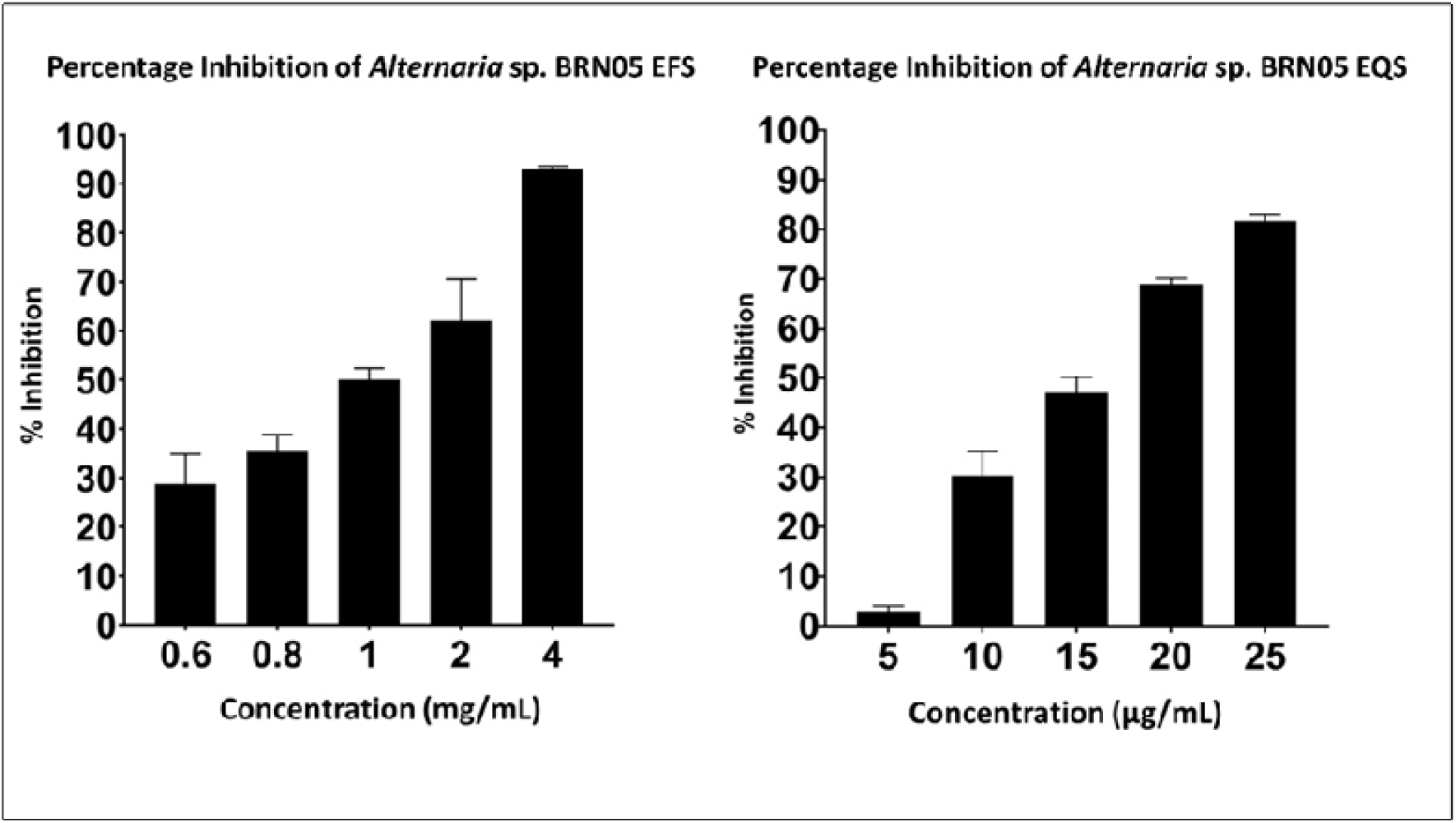
Percentage Inhibition of α-glucosidase by EFS and EQS from Alternaria sp. BRN05

### 3.3 Analysis of LC-MS/MS data in Negative mode

LC-MS/MS of EFS and EQS was run in negative mode. Total Ion Chromatography (TIC) were generated for both samples and the resulting chromatograms shown as Figure 3. Chromatograms of EFS and EQS extracts were analysed using the Agilent Personal Compound Database and Library (PCDL) software. Metabolite identification was conducted through precise mass measurement of precursor ions, comparing the experimental and theoretical isotopic patterns with those stored in the PCDL manager file, which contains a list of 850 metabolites. Based on the analysis, molecules from the chromatogram were categorized into "C" and "U" respectively. "C" represents metabolites common to both EFS and EQS, while "U" represents unique metabolites that are found only either in EFS or EQS. For example, compound 1CFS and 1CQS represent molecule 1, which is common in both EFS and EQS. 1 UQS represents molecule 1 which is unique to only EQS. A total of 19 metabolites belonging to EFS and 20 metabolites belonging to EQS were identified. In the EFS, out of 19 metabolites identified from EFS, 13 were commonly found from both EFS and EQS. Whereas 6 were found from EFS, and other 7 were found only from EQS. These molecules have codes assigned in Table 2. All metabolites were identified based on their experimental masses, achieving an overall identification score of over 80% for the isotopic pattern. This score was calculated considering exact masses, relative abundance, and spacing, with a mass error of less than 5 ppm. Total ion chromatogram (TIC) is shown below in Figure 3 (A) & 3 (B). Table 1 contains the list of compounds with assigned code, retention time, tentative or putative compound proposed, molecular formula, *m/z*, molecular ion form, identified score, MS/MS fragments, and the total number of fragments identified using *in-silico* fragmentation tool MetFrag(Ruttkies et al., 2016). Additional information about the MS/MS spectrum of each molecule, as well as the corresponding fragmentation pattern, is provided in supplementary file 2.

**Table 2. (A).** Molecules detected in the ethyl acetate fraction of *Alternaria* sp. BRN05, in both EFS and EQS and details of retention time, molecular formula, observed score, identified score, MS/MS fragments and the number of fragments identified in Metfrag.

| S. No | Retention time (Min) | Tentative/ putative compound proposed | Molecular Formula | *m/z* | Molecular ion form | Identified Score | MS/MS fragments | No of fragments identified in Metfrag |
| --- | --- | --- | --- | --- | --- | --- | --- | --- |
| 1CFS | 12.918 | 4-Hydroxymellein | C<sub>10</sub>H<sub>10</sub>O<sub>4</sub> | 221.0456 | (M+HCOO)-<br>[-H<sub>2</sub>O] | 98.25 | 177.0561, [C<sub>9</sub>H<sub>5</sub>O<sub>4</sub>]-<br>175.04, [C<sub>10</sub>H<sub>7</sub>O<sub>3</sub>]-<br>162.032, [C<sub>9</sub>H<sub>6</sub>O<sub>3</sub>]-<br>134.0863, [C<sub>8</sub>H<sub>7</sub>O<sub>2</sub>]- | 3 |
| 1CQS | 18.181 | 4-Hydroxymellein | C<sub>10</sub>H<sub>10</sub>O<sub>4</sub> | 221.0458 | (M+HC<br>O)- [-H<sub>2</sub>O] | 97.72 | 178.027, [C<sub>9</sub>H<sub>6</sub>O<sub>4</sub>]-<br>177.0553, [C<sub>9</sub>H<sub>5</sub>O<sub>4</sub>]-<br>162.034, [C<sub>9</sub>H<sub>6</sub>O<sub>3</sub>]- | 5 |
| 2CFS | 12.318 | 5'-Epialtenuen | C<sub>15</sub>H<sub>16</sub>O<sub>6</sub> | 291.0874 | (M-H)- | 98.93 | 273.0776, [C<sub>15</sub>H<sub>13</sub>O<sub>5</sub>]-<br>247.0984, [C<sub>14</sub>H<sub>15</sub>O<sub>4</sub>]-<br>231.0665, [C<sub>13</sub>H<sub>11</sub>O<sub>4</sub>]- | 12 |
|  |  |  |  |  |  |  | 186.0715, [C<sub>11</sub>H<sub>6</sub>O<sub>3</sub>]- |  |
| 2CQS | 16.664 | 5'-Epialtenuen | C<sub>15</sub>H<sub>16</sub>O<sub>6</sub> | 291.0878 | (M-H)- | 95.43 | 291.087, [C<sub>15</sub>H<sub>15</sub>O<sub>6</sub>]-<br>274.8918, [C<sub>15</sub>H<sub>14</sub>O<sub>5</sub>]-<br>230.0569, [C<sub>14</sub>H<sub>14</sub>O<sub>3</sub>]-<br>203.0343, [C<sub>12</sub>H<sub>11</sub>O<sub>3</sub>]- | 31 |
| 3CFS | 6.832 | Alternarienonic acid | C<sub>14</sub>H<sub>14</sub>O<sub>6</sub> | 278.0792 | (M-H)- | 98.67 | 277.0716, [C<sub>14</sub>H<sub>12</sub>O<sub>6</sub>]-<br>251.0563, [C<sub>12</sub>H<sub>11</sub>O<sub>6</sub>]-<br>246.9260, [C<sub>13</sub>H<sub>10</sub>O<sub>5</sub>]-<br>233.0813, [C<sub>13</sub>H<sub>13</sub>O<sub>4</sub>]-<br>214.09257, [C<sub>13</sub>H<sub>10</sub>O<sub>3</sub>]-<br>189.0542, [C<sub>11</sub>H<sub>9</sub>O<sub>3</sub>]- | 15 |
| 3CQS | 15.025 | Alternarienonic acid | C<sub>14</sub>H<sub>14</sub>O<sub>6</sub> | 278.0792 | (M-H)- | 98.43 | 277.0716, [C<sub>14</sub>H<sub>12</sub>O<sub>6</sub>]-<br>233.082, [C<sub>13</sub>H<sub>13</sub>O<sub>4</sub>]-<br>218.0584, [C<sub>12</sub>H<sub>10</sub>O<sub>4</sub>]-<br>205.086, [C<sub>12</sub>H<sub>13</sub>O<sub>3</sub>]-<br>190.0639, [C<sub>11</sub>H<sub>9</sub>O<sub>3</sub>]- | 18 |
| 4CFS | 15.753 | (+)-talaroflavone | C<sub>14</sub>H<sub>12</sub>O<sub>6</sub> | 257.0462 | (M-H)- [-<br>H<sub>2</sub>O] | 96.01 | 257.0461, [C<sub>14</sub>H<sub>10</sub>O<sub>5</sub>]-<br>246.9222, [C<sub>13</sub>H<sub>11</sub>O<sub>5</sub>]-<br>229.0516, [C<sub>13</sub>H<sub>11</sub>O<sub>4</sub>]-<br>213.0548, [C<sub>13</sub>H<sub>11</sub>O<sub>3</sub>]- | 6 |
| 4CQS | 21.168 | (+)-talaroflavone | C<sub>14</sub>H<sub>12</sub>O<sub>6</sub> | 257.0465 | (M-H)- [-<br>H<sub>2</sub>O] | 94.86 | 257.0457, [C<sub>14</sub>H<sub>10</sub>O<sub>5</sub>]-<br>228.9590, [C<sub>13</sub>H<sub>8</sub>O<sub>4</sub>]- | 8 |
|  |  |  |  |  |  |  | 215.0355, [C<sub>12</sub>H<sub>7</sub>O<sub>4</sub>]-<br>202.063, [C<sub>12</sub>H<sub>10</sub>O<sub>3</sub>]- |  |
| 5CFS | 10.022 | 2,5-dimethyl-7-hydroxychromone | C<sub>11</sub>H<sub>10</sub>O<sub>3</sub> | 189.0567 | (M-H)- | 92.64 | 189.0567, [C<sub>11</sub>H<sub>9</sub>O<sub>3</sub>]-<br>174.0324, [C<sub>10</sub>H<sub>6</sub>O<sub>3</sub>]-<br>160.08424, [C<sub>9</sub>H<sub>4</sub>O<sub>3</sub>]-<br>119.0491, [C<sub>7</sub>H<sub>3</sub>O<sub>2</sub>]- | 9 |
| 5CQS | 14.807 | 2,5-dimethyl-7-hydroxychromone | C<sub>11</sub>H<sub>10</sub>O<sub>3</sub> | 189.0563 | (M-H)- | 95.19 | 189.055, [C<sub>11</sub>H<sub>9</sub>O<sub>3</sub>]-<br>174.0355, [C<sub>10</sub>H<sub>6</sub>O<sub>3</sub>]-<br>162.8384, [C<sub>10</sub>H<sub>10</sub>O<sub>2</sub>]-<br>105.0328, [C<sub>11</sub>H<sub>7</sub>O]-<br>146.0358, [C<sub>9</sub>H<sub>7</sub>O<sub>2</sub>]-H- | 6 |
| 6CFS | 10.756 | Alternarian acid | C<sub>15</sub>H<sub>12</sub>O<sub>8</sub> | 319.0464 | (M-H)- | 96.33 | 231.0659, [C<sub>13</sub>H<sub>10</sub>O<sub>4</sub>+H]-<br>203.0699, [C<sub>12</sub>H<sub>10</sub>O<sub>3</sub>+H]-<br>188.0467, [C<sub>11</sub>H<sub>9</sub>O<sub>3</sub>]-H-<br>160.8409, [C<sub>10</sub>H<sub>9</sub>O<sub>2</sub>]-H- | 8 |
| 6CQS | 15.882 | Alternarian acid | C<sub>15</sub>H<sub>12</sub>O<sub>8</sub> | 319.0461 | (M-H)- | 98.03 | 231.0656, [C<sub>13</sub>H<sub>10</sub>O<sub>4</sub>+H]-<br>216.0434, [C<sub>12</sub>H<sub>9</sub>O<sub>4</sub>]-H-<br>203.0712, [C<sub>12</sub>H<sub>10</sub>O<sub>3</sub>+H]-<br>188.0476, [C<sub>11</sub>H<sub>9</sub>O<sub>3</sub>]-H- | 15 |
| 7CFS | 6.618 | 1,8-dihydroxynaphthalene | C<sub>10</sub>H<sub>8</sub>O<sub>2</sub> | 205.0204 | (M+HCOO)- | 97.77 | 149.0601, [C<sub>9</sub>H<sub>8</sub>O<sub>2</sub>]-<br>118.042, [C<sub>8</sub>H<sub>6</sub>O]- | 5 |
|  |  |  |  |  |  |  | 75.0032, [C<sub>6</sub>H<sub>3</sub>O]-<br>105.0338, [C<sub>7</sub>H<sub>5</sub>O]- |  |
| 7CQS | 10.951 | 1,8-dihydroxynaphthalene | C<sub>10</sub>H<sub>8</sub>O<sub>2</sub> | 205.0509 | (M+HCOO)- | 96.46 | 143.0502, [C<sub>10</sub>H<sub>7</sub>O]-<br>118.9939, [C<sub>8</sub>H<sub>6</sub>O]-<br>105.0351, [C<sub>7</sub>H<sub>5</sub>O]- | 13 |
| 8CFS | 11.999 | Aspergone Q | C<sub>11</sub>H<sub>14</sub>O<sub>6</sub> | 223.0609 | (M-H)- [-H<sub>2</sub>O] | 85.48 | 223.0606, [C<sub>11</sub>H<sub>13</sub>O<sub>5</sub>]-<br>191.0347, [C<sub>10</sub>H<sub>10</sub>O<sub>4</sub>-2H]-H-<br>162.0319, [C<sub>9</sub>H<sub>9</sub>O<sub>3</sub>-2H]-H-<br>122.0383, [C<sub>7</sub>H<sub>9</sub>O<sub>2</sub>-2H]-H- | 9 |
| 8CQS | 16.959 | Aspergone Q | C<sub>11</sub>H<sub>14</sub>O<sub>6</sub> | 223.0611 | (M-H)- [-H<sub>2</sub>O] | 84.46 | 191.0352, [C<sub>10</sub>H<sub>10</sub>O<sub>4</sub>-2H]-H-<br>149.0247, [C<sub>8</sub>H<sub>7</sub>O<sub>3</sub>-H]-H-<br>123.0462, [C<sub>7</sub>H<sub>9</sub>O<sub>2</sub>-H]-H- | 9 |
| 9CFS | 12.967 | 6-Epi-stemphytriol | C<sub>20</sub>H<sub>16</sub>O<sub>7</sub> | 349.0709 | (M-H)- [-H<sub>2</sub>O] | 96.12 | 349.0701, [C<sub>20</sub>H<sub>15</sub>O<sub>6</sub>-H]-H-<br>331.06, [C<sub>20</sub>H<sub>14</sub>O<sub>5</sub>-2H]-H-<br>145.029, [C<sub>9</sub>H<sub>5</sub>O<sub>2</sub>]-<br>117.0352, [C<sub>8</sub>H<sub>4</sub>O+H]- | 4 |
| 9CQS | 18.313 | 6-Epi-stemphytriol | C<sub>20</sub>H<sub>16</sub>O<sub>7</sub> | 349.0716 | (M-H)- [-H<sub>2</sub>O] | 99.01 | 331.0592, [C<sub>20</sub>H<sub>14</sub>O<sub>5</sub>-2H]-H-<br>292.061, [C<sub>14</sub>H<sub>12</sub>O<sub>7</sub>]-<br>243.0665, [C<sub>14</sub>H<sub>11</sub>O<sub>4</sub>]- | 4 |
| 10CFS | 13.254 | 12-Methoxycitromycin | C<sub>14</sub>H<sub>12</sub>O<sub>5</sub> | 287.0575 | (M+HCOO)- [-H<sub>2</sub>O] | 97.80 | 229.05, [C<sub>13</sub>H<sub>9</sub>O<sub>4</sub>]- | 17 |
|  |  |  |  |  |  |  | 228.0428, [C<sub>13</sub>H<sub>9</sub>O<sub>4</sub>]-<br>211.0402, [C<sub>13</sub>H<sub>8</sub>O<sub>3</sub>]-H-<br>200.0476, [C<sub>12</sub>H<sub>8</sub>O<sub>4</sub>]-<br>243.0665, [C<sub>14</sub>H<sub>11</sub>O<sub>4</sub>]- |  |
| 10CQS | 18.569 | 12-Methoxycitromycin | C<sub>14</sub>H<sub>12</sub>O<sub>5</sub> | 305.0665 | (M+HCOO)- | 93.62 | 229.05, [C<sub>13</sub>H<sub>9</sub>O<sub>4</sub>]-<br>228.0424, [C<sub>13</sub>H<sub>8</sub>O<sub>4</sub>]-<br>211.0368, [C<sub>13</sub>H<sub>8</sub>O<sub>3</sub>]-H-<br>200.0467, [C<sub>12</sub>H<sub>8</sub>O<sub>3</sub>]- | 15 |
| 11CFS | 14.251 | 4-Hydroxyalternariol<br>9-methyl ether | C<sub>15</sub>H<sub>12</sub>O<sub>6</sub> | 287.0565 | (M-H)- | 84.00 | 271.0609, [C<sub>15</sub>H<sub>11</sub>O<sub>5</sub>]-<br>245.0826, [C<sub>13</sub>H<sub>9</sub>O<sub>5</sub>]-<br>230.0586, [C<sub>13</sub>H<sub>10</sub>O<sub>4</sub>]-<br>202.0631, [C<sub>12</sub>H<sub>11</sub>O<sub>3</sub>]-<br>160.0527, [C<sub>10</sub>H<sub>9</sub>O<sub>2</sub>]- | 43 |
| 11CQS | 23.981 | 4-Hydroxyalternariol<br>9-methyl ether | C<sub>15</sub>H<sub>12</sub>O<sub>6</sub> | 287.0571 | (M-H)- | 92.02 | 287.0571, [C<sub>15</sub>H<sub>11</sub>O<sub>6</sub>]-<br>272.0331, [C<sub>14</sub>H<sub>8</sub>O<sub>6</sub>]-<br>257.0181, [C<sub>13</sub>H<sub>5</sub>O<sub>6</sub>]-<br>228.0422, [C<sub>14</sub>H<sub>12</sub>O<sub>3</sub>]-<br>188.047, [C<sub>11</sub>H<sub>9</sub>O<sub>3</sub>]-H- | 21 |
| 12CFS | 15.056 | Orthosporin | C<sub>12</sub>H<sub>12</sub>O<sub>5</sub> | 235.0614 | (M-H)- | 86.45 | 191.071, [C<sub>11</sub>H<sub>12</sub>O<sub>3</sub>]-H-<br>189.0549, [C<sub>11</sub>H<sub>11</sub>O<sub>3</sub>-H]-H-<br>176.0475, [C<sub>10</sub>H<sub>6</sub>O<sub>3</sub>+2H]- | 7 |
|  |  |  |  |  |  |  | 174.0316, [C<sub>10</sub>H<sub>6</sub>O<sub>3</sub>]- |  |
| 12CQS | 20.357 | Orthosporin | C<sub>12</sub>H<sub>12</sub>O<sub>5</sub> | 235.061 | (M-H)- | 84.64 | 191.0711, [C<sub>11</sub>H<sub>12</sub>O<sub>3</sub>]-H-<br>189.0556, [C<sub>11</sub>H<sub>11</sub>O<sub>3</sub>-H]-H-<br>176.0472, [C<sub>10</sub>H<sub>6</sub>O<sub>3</sub>+2H]-<br>161.0248, [C<sub>9</sub>H<sub>4</sub>O<sub>3</sub>+H]-<br>148.0522, [C<sub>9</sub>H<sub>7</sub>O<sub>2</sub>+H]- | 10 |
| 13CFS | 19.838 | Altenusin | C<sub>15</sub>H<sub>14</sub>O<sub>6</sub> | 317.0657 | (M+HCOO)-<br>[-H<sub>2</sub>O] | 8.96 | 272.068, [C<sub>15</sub>H<sub>12</sub>O<sub>5</sub>]-<br>271.016, [C<sub>15</sub>H<sub>11</sub>O<sub>5</sub>]-<br>270.0149, [C<sub>15</sub>H<sub>10</sub>O<sub>5</sub>]-<br>242.0223, [C<sub>14</sub>H<sub>10</sub>O<sub>4</sub>]-<br>198.0313, [C<sub>13</sub>H<sub>10</sub>O<sub>2</sub>]- | 4 |
| 13CQS | 22.856 | Altenusin | C<sub>15</sub>H<sub>14</sub>O<sub>6</sub> | 317.0666 | (M+HCOO)-<br>[-H<sub>2</sub>O] | 98.27 | 271.024, [C<sub>15</sub>H<sub>11</sub>O<sub>5</sub>]-<br>270.169, [C<sub>15</sub>H<sub>10</sub>O<sub>5</sub>]-<br>242.0206, [C<sub>14</sub>H<sub>10</sub>O<sub>4</sub>]-<br>214.0267, [C<sub>13</sub>H<sub>10</sub>O<sub>3</sub>]-<br>198.0313, [C<sub>13</sub>H<sub>10</sub>O<sub>2</sub>]- | 17 |

**Table 2. (B):** Molecules detected in the ethyl acetate fraction of *Alternaria* sp. BRN05, that are unique to EFS and details of retention time, molecular formula, observed score, identified score, MS/MS fragments and the number of fragments identified in Metfrag.

| S. No | Retention time (Min) | Tentative/putative compound proposed | Molecular Formula | m/z | Molecular ion form | Identified Score | MS/MS fragments | No of fragments identified in Metfrag |
| --- | --- | --- | --- | --- | --- | --- | --- | --- |
| 1UFS | 5.487 | 4-Ethylcatechol | C<sub>8</sub>H<sub>10</sub>O<sub>2</sub> | 119.0501 | (M-H)-[-H<sub>2</sub>O] | 87.35 | 119.05, [C<sub>8</sub>H<sub>9</sub>O-H]-H-<br>93.0340, [C<sub>6</sub>H<sub>4</sub>+H]-<br>61.9881, [C<sub>2</sub>H<sub>4</sub>O<sub>2</sub>]- | 4 |
| 2UFS | 6.93 | *p*-Coumaric acid | C<sub>9</sub>H<sub>8</sub>O<sub>3</sub> | 209.0458 | (M+HCOO)- | 98.57 | 123.0453, [C<sub>7</sub>H<sub>7</sub>O<sub>2</sub>]-<br>81.0343, [C<sub>5</sub>H<sub>6</sub>O]- | 5 |
| 3UFS | 9.327 | Diaportinol | C<sub>13</sub>H<sub>14</sub>O<sub>6</sub> | 247.0613 | (M-H)-[-H<sub>2</sub>O] | 85.17 | 219.8431, [C<sub>12</sub>H<sub>11</sub>O<sub>4</sub>]-<br>210.9479, [C<sub>10</sub>H<sub>10</sub>O<sub>4</sub>]-<br>203.703, [C<sub>12</sub>H<sub>11</sub>O<sub>3</sub>]-<br>188.0469, [C<sub>11</sub>H<sub>8</sub>O<sub>3</sub>]-<br>166.9587, [C<sub>9</sub>H<sub>10</sub>O<sub>3</sub>]-<br>161.0600, [C<sub>10</sub>H<sub>9</sub>O<sub>2</sub>]- | 9 |
| 4UFS | 5.677 | Phenylacetic acid | C<sub>8</sub>H<sub>8</sub>O<sub>2</sub> | 163.04 | (M+HCOO)-<br>[-H<sub>2</sub>O] | 87.29 | 119.0504, [C<sub>8</sub>H<sub>7</sub>O]-<br>91.0549, [C<sub>7</sub>H<sub>7</sub>]-<br>63.9965, [C<sub>4</sub>H<sub>3</sub>O-2H]-H- | 5 |
| 5UFS | 16.764 | Procyanidin dimer B1 | C<sub>30</sub>H<sub>26</sub>O<sub>12</sub> | 605.1305 | (M+HCOO)-<br>[-H<sub>2</sub>O] | 98 | 486.168, [C<sub>29</sub>H<sub>26</sub>O<sub>2</sub>]-<br>485.125, [C<sub>29</sub>H<sub>25</sub>O<sub>2</sub>]-<br>275.0545, [C<sub>14</sub>H<sub>12</sub>O<sub>6</sub>]-H-241.0491, [C<sub>14</sub>H<sub>12</sub>O<sub>4</sub>-2H]-H-61.9884, [C<sub>2</sub>H<sub>3</sub>O<sub>2</sub>]-[+H] | 5 |
| 6UFS | 22.491 | Theaflavin | C<sub>29</sub>H<sub>24</sub>O<sub>12</sub> | 591.1139 | (M+HCOO)-<br>[-H<sub>2</sub>O] | 96.34 | 515.0986, [C<sub>28</sub>H<sub>21</sub>O<sub>10</sub>-H]-<br>H-<br>203.0709, [C<sub>12</sub>H<sub>10</sub>O<sub>3</sub>-H]-<br>188.0461[C<sub>11</sub>H<sub>9</sub>O<sub>3</sub>]-H-<br>92.9968, [C<sub>5</sub>H<sub>4</sub>O<sub>2</sub>-H]- | 11 |

**Table 2. (C):**
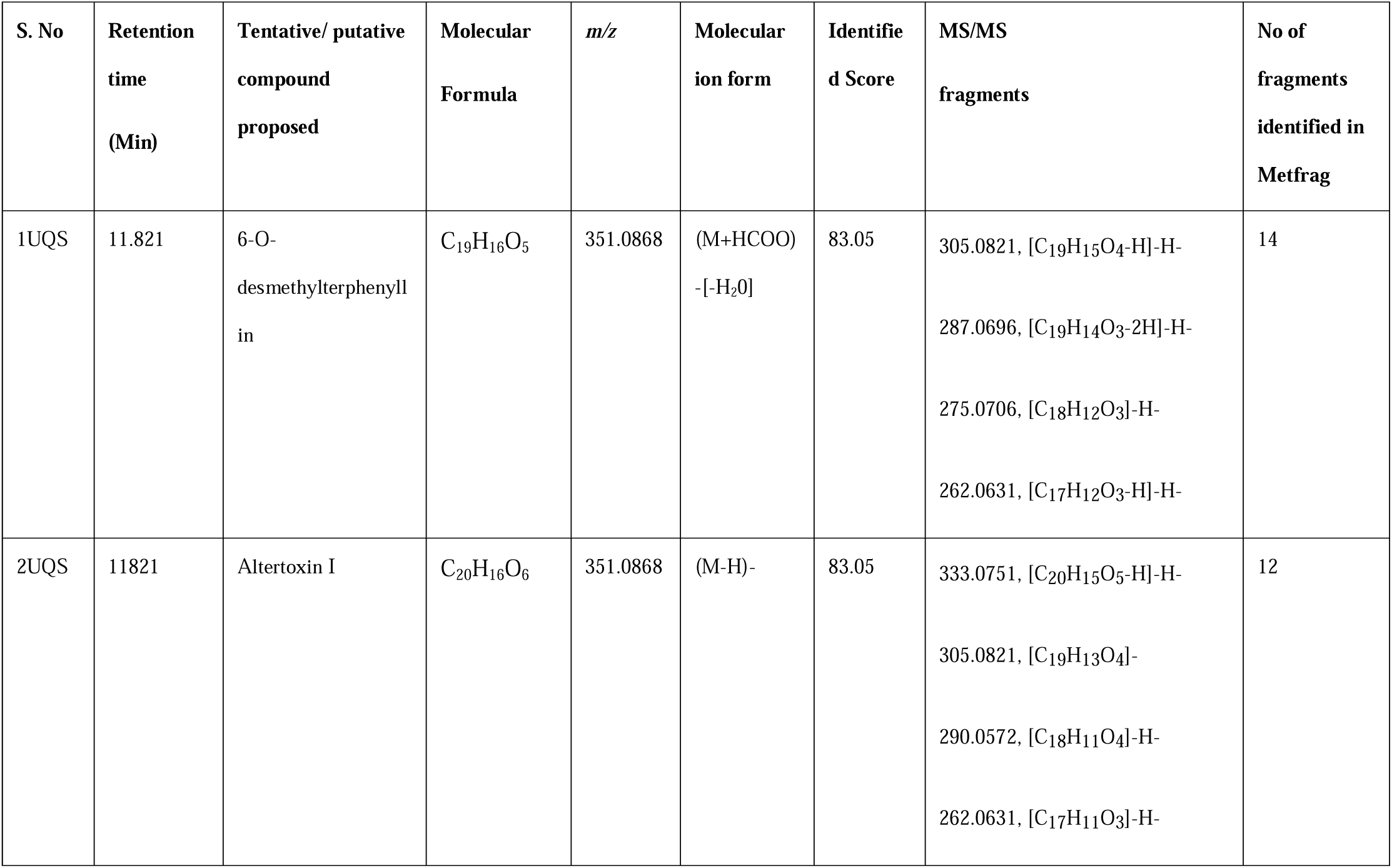
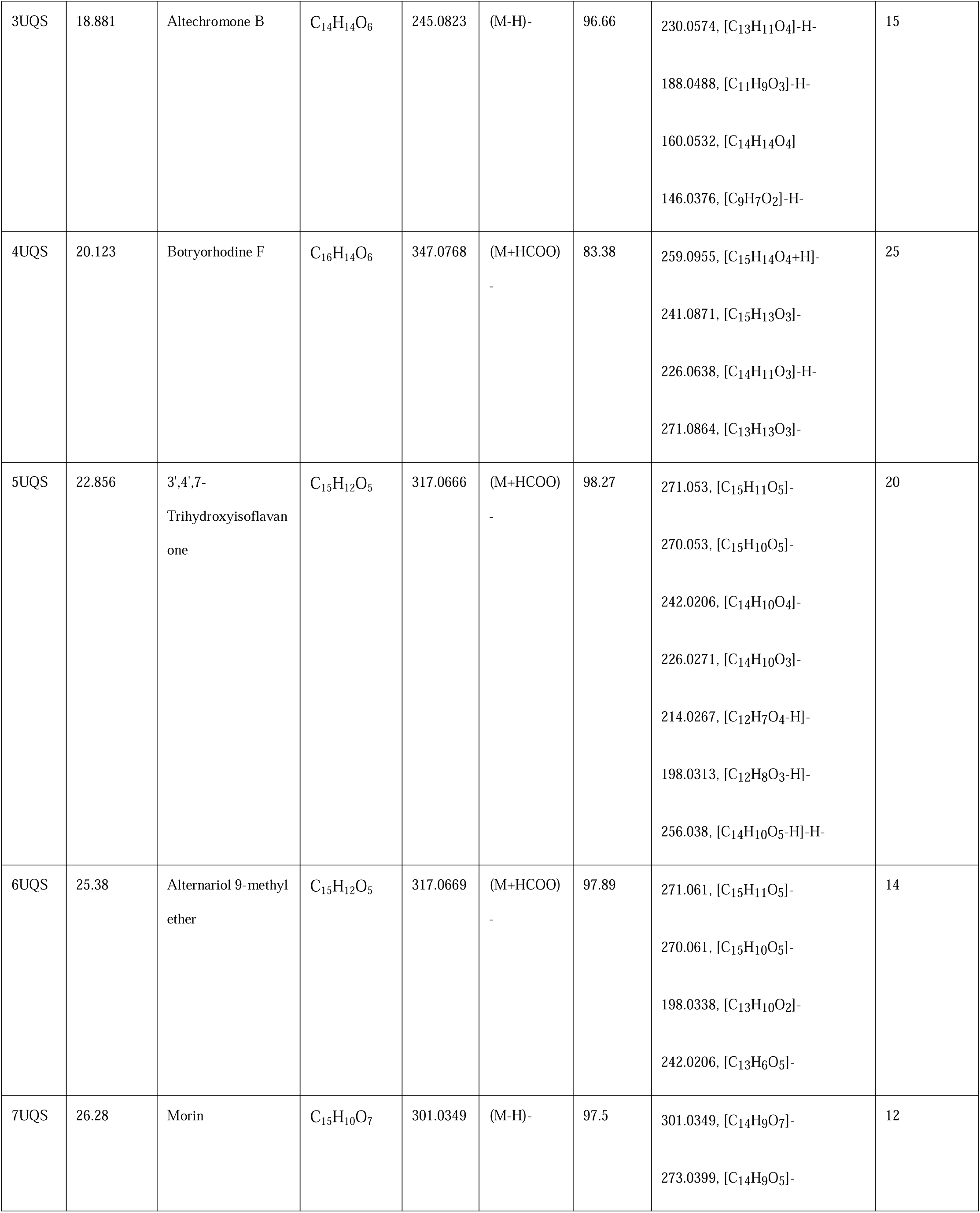
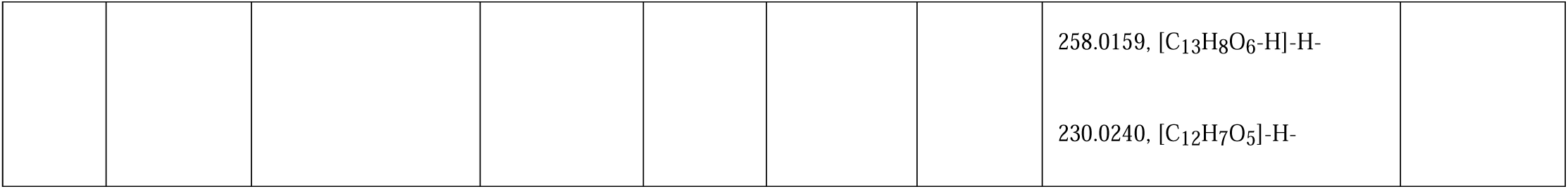
Molecules detected in the ethyl acetate fraction of *Alternaria* sp. BRN05, that are unique EQS and details of retention time, molecular formula, observed score, identified score, MS/MS fragments and the number of fragments identified in Metfrag.

**Figure 3 (A).**
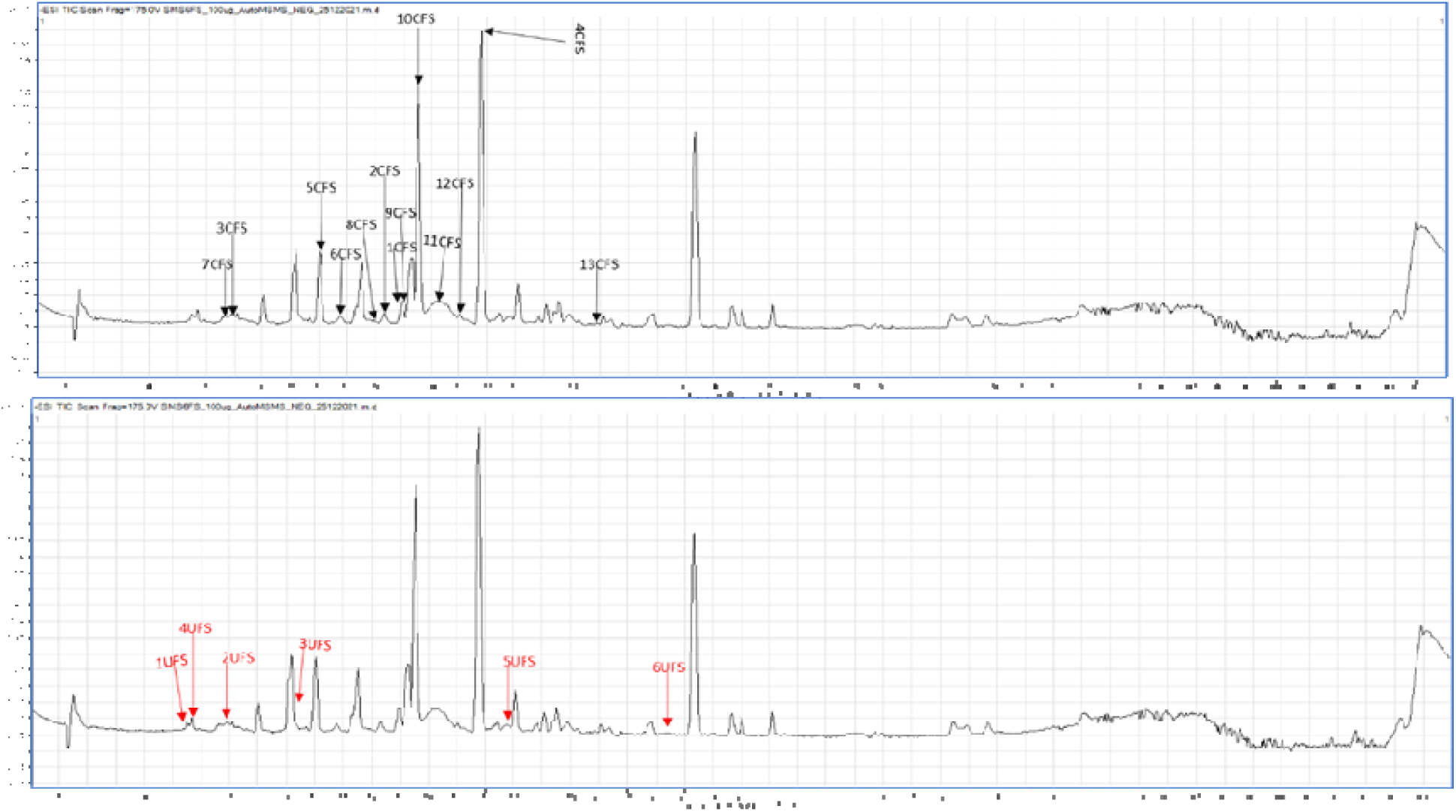
Metabolic profiling using UHPLC-ESI-QTOF-MS in negative mode for the Alternaria sp. BRN05. The top chromatogram highlights thirteen metabolites found in EFS, common to EQS (1CFS to 13 CFS). Bottom chromatograms show six metabolites found unique to EFS (1UFS to 6FS).

**Figure 3 (B).**
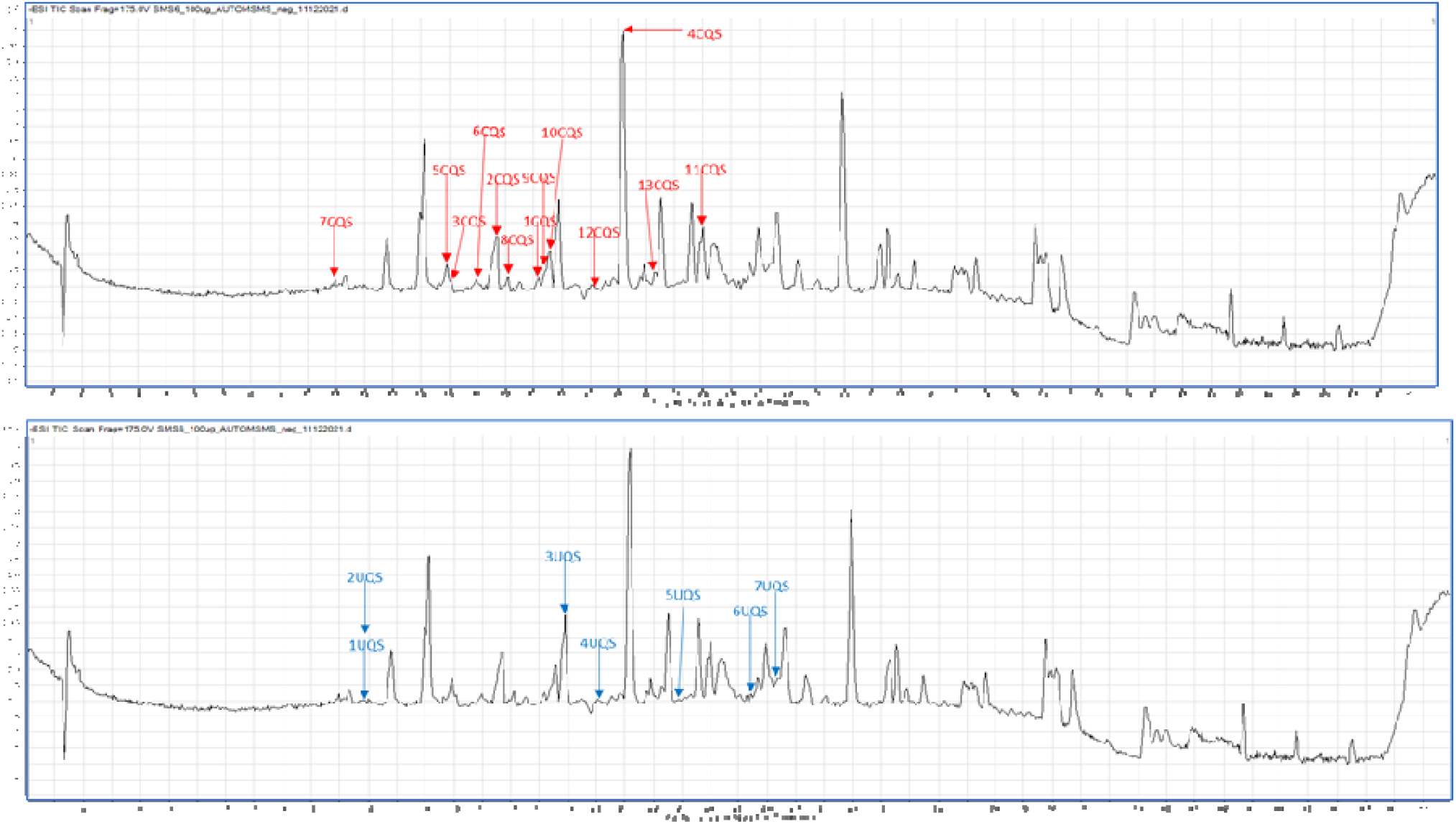
Metabolic profiling using UHPLC-ESI-QTOF-MS in negative mode for the Altern ria sp. BRN05. The top chromatogram highlights thirteen metabolites found in EQS, common to EFS (1 CQS to 13 CQS). Bottom chromatograms show seven metabolites found unique to EQS (1UQS to 7QS).

The MS/MS fragments obtained for each peak in the TIC were used for the tentative identification of the compounds. The MS/MS fragments were imported to MetFrag to help identify the fragment losses. Along with the MetFrag generated from *in-silico* fragmentation, the chemistry involved in the losses of various fragments was studied using ChemDraw Pro 8. The identification of a few representative compounds has been described below.

The theoretical molecular mass for Molecule 1 is 194.0594 Da (C□□H□□O□)□ It was found that three fragments of 1CFS and five fragments of 1CQS matched the predicted 4-Hydroxymellein fragments generated using MetFrag. Additionally, various fragment losses were studied with the help of ChemDraw. A common fragment at *m/z* 177 [C□□H□□O□]□, produced by both 1CFS and 1CQS, corresponds to the loss of a methyl radical (CH□) from the deprotonated molecule [(M-H)-CH□]. Another fragment at *m/z* 162 [C□□H□□O□]□, also observed in both, signifies the loss of methanol (CH□OH) from the deprotonated molecule [(M-H)-CH□OH]. Further analysis of 1CFS revealed fragments at *m/z* 175 [C□□H□□O□]□, resulting from the loss of water (H□O) from the deprotonated molecule [(M-H)-H□O], and at *m/z* 134, which corresponds to the combined loss of a methyl radical (CH□) and carbon dioxide (CO□) from the deprotonated molecule [(M-H)-CH□-CO□] Finally, 1CQS displayed a fragment at *m/z* 178, indicating the loss of a methyl radical (CH□) from the deprotonated molecule [(M-H)-CH□]□ Based on the consistent fragmentation patterns observed in both 1CFS and 1CQS, the identity of Molecule 1 as 4-Hydroxymellein can be tentatively confirmed.

Cleavage at various positions of the molecules using ChemDraw produces numerous fragments with different *m/z* values. For 2CFS, these fragments include those at *m/z* 273 [C□□H□□O□]□, 247 [C□□H□□O□]□, 231 [C□□H□□O□]□, and 186 [C□□H□□O□]□. The fragment at *m/z* 186 likely corresponds to losses of water [(M-H)-H□O], carbon dioxide [(M-H)-CO□], two formaldehyde molecules [(M-H)-2CH□O]□, and a combination of formaldehyde, methanol, and a methyl radical [(M-H)-CH□O-CH□OH-CH□]□. Similar analysis of 2CQS MS/MS fragments revealed fragments at various *m/z* values, including 291 [C□□H□□O□]□, 274 [C□□H□□O□]□, 230 [C□□H□□O□]□, and 203 [C□□H□□O□]□. These fragments are likely due to the loss of a proton [M-H]□, water [(M-H)-H□O], a combination of a hydroxyl and carbon dioxide group [(M-□H)-OH-CO□]□, and a methyl radical, carbonyl group, and formaldehyde [(M-H)-CH□-CO-CH□O]□, respectively. Based on the consistent fragmentation patterns observed in both 2CFS and 2CQS, which matched those predicted for 5’-Epialtenuen, the identity of the molecule can be tentatively confirmed as 5’-Epialtenuen. These fragmentations are explained in Figure 3 (C).

**Figure 3 (C).**
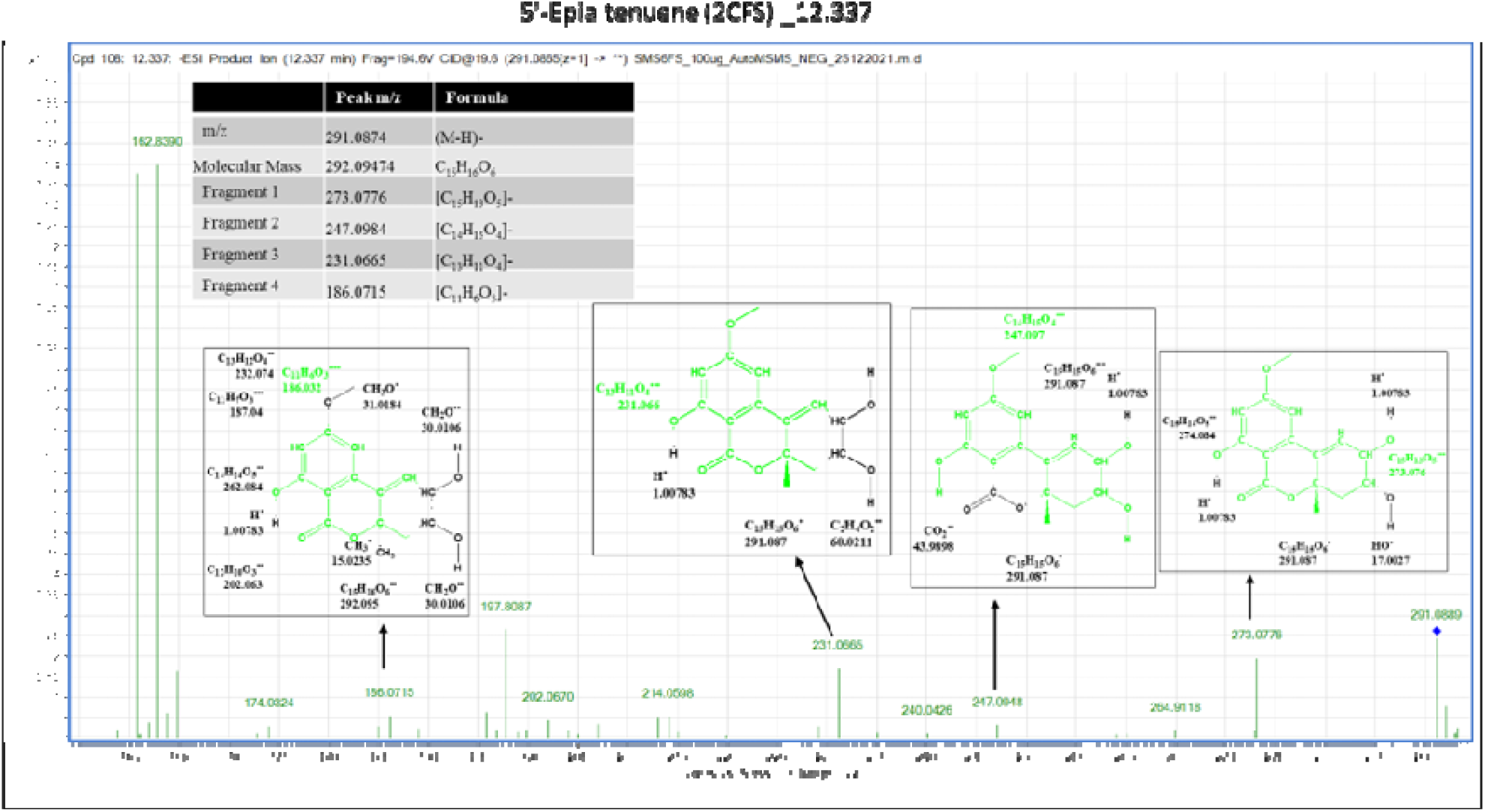
Fragmentation analysis was performed for 5’-Epialtenuene (2CFS) using two software programs: Metfrag and ChemDraw Pro 8.0. Metfrag predicted fragments at m/z 273.0776 231.0665. ChemDraw Pro 8.0 obtained fragments at m/z 247.098 and 186.0715.

Compounds 3CFS and 3CQS have a theoretical molecular mass of 278.0792, corresponding to the formula C□□H□□O□. MetFrag analysis identified fifteen fragments for 3CFS and eighteen fragments for 3CQS that matched the predicted fragmentation pattern. ChemDraw software was further employed to analyze some specific fragments. For 3CFS, fragments were observed at *m/z* 277 [C□□H□□O□]□, 246 [C□□H□□O□]□, 233 [C□□H□□O□]□, and 214 [C□□H□□O□]□.

These fragments likely correspond to the loss of various moieties from the parent molecule: [(M-H)]□ (loss of a proton), [(M-H)-OH-CH□]□(loss of a hydroxyl and a methyl group), [(M-H)-CO□] (loss of carbon dioxide), and [(M-H)-OH-CH O-OH] (loss of a hydroxyl, a methoxy group, and a hydroxyl group), respectively. Similarly, fragments in 3CQS at *m/z* 277 [C□□H□□O□]□, 233 [C□□H□□O□]□, 218 [C□□H□□O□]□, and 205 [C□H□O□] corresponded to the loss of [(M-H)] (proton), [(M-H)-CO□]□ (carbon dioxide), [(M-H)-CO□-CH□]□ (carbon dioxide and a methyl group), and [(M-H)-CO-CHO□]□ (carbon monoxide and a formyl group), respectively. Notably, fragments at *m/z* 233, 215, and 161 were also reported in literature for this molecule(Kjer, 2009). Based on the consistent fragmentation patterns observed in both 3CFS and 3CQS, along with the presence of key fragments identified in the literature, the molecule can be tentatively confirmed as Alternarienonic acid.

4CFS and 4CQS exhibit a theoretical molecular mass of 276.0634. MetFrag analysis identified six and eight fragments for 4CFS and 4CQS, respectively. Notably, a significant fragment in our spectra at *m/z* 257 [C□□H□□O□]□ corresponding to loss of a water [(M-H)-H□O], matches a fragment reported in the literature (Zhao et al., 2021). Further analysis using ChemDraw revealed that fragments in 4CFS at *m/z* 246 [C□□H□□O□]□ and 229 [C□□H□□O□]□ likely correspond to the loss of a methoxy group [(M-H)-CH□O]□ and a combination of water and carbon monoxide [(M-H)-H□O-CO]□, respectively. Similarly, fragments in 4CQS at *m/z* 228 [C□□H□□O□]□, 215 [C□□H□□O□]□, and 202 [C□□H□□O□]□ likely correspond to the loss of a hydroxyl and methoxy group [(M-H)-OH-CH□O]□, a combination of hydroxyl, methoxy, and methane [(M-H)-OH-CH□O-CH□]□, and a combination of hydroxyl, methoxy, and formaldehyde [(M-H)-OH-CH□O-□CHO]□, respectively. Based on these consistent fragmentation patterns, the compounds can be tentatively confirmed as (+)-talaroflavones.

Compounds 5CFS and 5CQS exhibit a theoretical molecular mass of 190.063. MetFrag analysis identified nine and six matching fragments for 5CFS and 5CQS, respectively. A fragment at *m/z* 146, reported in the literature [15], was also observed in our MS/MS spectra. Both molecules shared common fragments at *m/z* 189 [C□□H□□O□]□ and 174 [C□□H□□O□]□, corresponding to the loss of a proton [(M-H)-H]^-^] and a methyl group [(M-H)-CH□]□, respectively. Further analysis of 5CFS revealed fragments at *m/z* 160 [C□□H□□O□]□ and 119 [C□□H□□O□]□, consistent with the loss of a methyl group and a combination of methyl, hydroxyl, and acetylene groups: [(M-H)-CH□-OH]□and [(M-H)-CH□-CH□-C□H□O]□, respectively. Similarly, fragments in 5CQS at *m/z* 162 [C□□H□□O□]□ and 105 [C□H□]^-^ corresponded to the loss of a formyl group [(M-H)-COH]^-^ and a combination of formyl, methoxy, and acetylene groups [(M-H)-COH-CH□O-C□H□]□, respectively. Notably, the fragment at *m/z* 189 was the most prominent peak observed in both 5CFS and 5CQS. Based on these consistent fragmentation patterns and the presence of a key fragment identified in the literature, the molecules can be tentatively confirmed as 2,5-dimethyl-7-hydroxychromone. The detailed fragmentation patterns for all other molecules are provided in supplementary data 2. The most prominent peaks for other molecules are explained below.

The theoretical molecular mass of 6CFS and 6CQS is 320.0532. MetFrag analysis identified eight and fifteen fragments matching 6CFS and 6CQS, respectively. Several prominent peaks in the MS/MS spectra matched those predicted by MetFrag, including fragments at *m/z* 231.0656, 216.0434, and 203.0712. According to the literature, fragment 188 is present in the MS/MS spectra of both 6CFS and 6CQS, while MetFrag fragmentation analysis of 6CQS identified fragments 188 and 160 (Kjer, 2009). Based on these consistent fragmentation patterns, the compounds can be tentatively identified as Alternarian acid.

The theoretical molecular mass of compounds 7CFS and 7CQS was 160.0523. For 7CFS, five fragments, and for 7CQS, thirteen fragments matched the predicted fragments of 1,8-dihydroxynaphthalene. The matched MS/MS fragments in the EQS were at *m/z* 143.0502, 149.0601, 118.993, and 105.0351, while those in the EFS were at *m/z* 149.0601, 118.993, and 105.0338. Thus, these molecules can be putatively confirmed as 1,8-dihydroxynaphthalene. Compound 8CFS and 8CQS have a theoretical molecular mass of 242.079. Nine *in-silico* fragments matched for both. For 8CFS, significant fragments matched the MetFrag results, including *m/z* 223.0606, 191.0347, 162.0319, and 122.0383. The matched fragments for 8CQS were 191.0352, 149.0247, and 123.0462. Based on these results, the molecules can be tentatively confirmed as Aspergone Q. Compounds 9CFS and 9CQS have a theoretical molecular mass of 242.079. Four *in silico* fragments matched for both. For 9CFS, significant fragments matched the MetFrag results, including *m/z* 349.0701, 311.06, 145.029, and 117.035. The matched fragments for 9CQS were 331.0592, 292.061, and 243.0665. Based on these fragment patterns, the molecules can be tentatively confirmed as 6-Epi-stemphytriol.

Both 10CFS and 10CQS exhibit a theoretical molecular mass of 260.0698. MetFrag analysis identified twelve matching fragments for 10CFS and fifteen for 10CQS. ChemDraw software was then employed to explore various fragment losses from both molecules. Notably, common fragments were observed at *m/z* 229 [C□□H□□O□]□, 228 [C□□H□□O□]□, and 200 [C□□H□□O□]□, corresponding to the loss of formaldehyde [(M-H)-CH O], methoxy [(M-H)-CH O], and another methoxy group along with carbonyl loss [(M-H)-CH O-CO], respectively. Additionally, MetFrag analysis identified specific fragments for 10CFS at *m/z* 243.0661 and 160.053, while 10CQS displayed fragments at *m/z* 243.0654 and 211.0386. Based on these consistent fragmentation patterns and the matches identified by MetFrag, the identity of the molecules can be tentatively confirmed as 12-Methoxycitromycin.

Compounds 11CFS and 11CQS have a theoretical molecular mass of 288.0634. MetFrag analysis identified 43 fragments matching the predicted fragmentation pattern for 11CFS, while only 21 fragments matched for 11CQS. Significant fragments observed in the MS/MS spectra of 11CFS were at *m/z* 271.0609, 245.0826, 230.0568, 202.0631, and 160.0527. These fragments likely correspond to the loss of various moieties from the parent molecule. In comparison, the MS/MS spectra of 11CQS displayed significant fragments at *m/z* 287.0571, 272.0331, 257.0181, 228.0422, and 188.0047. Based on the consistent fragmentation patterns observed in both molecules and their matches to predicted fragments, 11CFS and 11CQS can be tentatively identified as 4-Hydroxyalternariol 9-methyl ether.

12CFS and 12CQS have a theoretical molecular mass of 236.0687. MetFrag analysis identified ten fragments matching the predicted fragmentation pattern for 12CQS, whereas only seven matched for 12CFS. A fragment at *m/z* 191 (Tseng et al., 2014) was confirmed by the literature for both 12CFS and 12CQS, and another most abundant fragment at *m/z* 189.0549 likely corresponds to the loss of [M-H-COH-H O]. The significant fragments identified for 12CFS were 176.0457 and 174.0316, while those matched for 12CQS were 176.0472, 161.0248, and 148.0522. Based on the consistent fragmentation patterns observed in both molecules and their matches to literature and predicted fragmentation, 12CFS and 12CQS can be tentatively confirmed as Orthosporin.

13CFS and 13CQS have a theoretical molecular mass of 290.0781. MetFrag analysis identified four fragments matching the predicted fragmentation pattern for 13CFS, while 17 fragments matched for 13CQS. A literature search confirmed that a fragment at *m/z* 202.02 was present in the 13CFS spectra, and another fragment at *m/z* 256 was identified in the 13CQS spectra (Kjer, 2009) The prominent peaks in the MS/MS spectra of 13CFS were at *m/z* 271.0149, 270.0149, and 242.0223. For 13CQS, the most prominent peaks were at *m/z* 271.019, 270.169, 214.0267, and 198.0313, which were further analyzed using ChemDraw. Based on the consistent fragmentation patterns observed in both molecules and their matches to literature, 13CFS and 13CQS can be tentatively confirmed as Altenusin.

The molecule 1UFS has theoretical molecular mass of 138.0679. Five fragments, 61.9881, 65.0391, 93.034 and 119.05, of the MS/MS spectra matched to the predicted fragments of the molecule 4-Ethylcatechol. These fragments also matched with the fragments reported in the literature(Allen et al., 2015). Thus, molecule can be tentatively confirmed as 4-Ethylcatechol. For 2UFS (theoretical molecular mass 164.0476), fragments of the MS/MS spectra matched the predicted fragments of the molecule *p*-Coumaric acid. *p*-Coumaric acid fragments at *m/z* 123.0453, 95.0129, 65.0396, 81.0343 (LC-MS/MS Spectrum - LC-ESI-QTOF (UPLC Q-Tof Premier, Waters), Negative (DB04066) | DrugBank Online, n.d.) and are the same as the fragments reported in the literature. The prominent peak was observed at 160.8422 and 118.9943. Thus, the molecule can be tentatively confirmed as *p*-Coumaric acid. Compound 3UFS has a molecular mass of 266.0791. Nine peaks of the MS/MS spectra matched the predicted fragments of the molecule Diaportinol. A few of 3UFS fragments are at *m/z* 219.8431, 210.9479, 203.703, 188.0469, and 166.9587. Thus, this molecule is tentatively confirmed as Diaportinol. 4UFS has a molecular mass of 136.0523. Five peaks of the MS/MS spectra matched the predicted fragments of the molecule. Phenylacetic acid fragments at *m/z* 119.0504 and 91.0549 (Phenylacetic acid Mass Spectrum, n.d.) are the same as the fragments reported in the literature. Thus, the molecule is tentatively confirmed as Phenylacetic acid. 5UFS has a molecular mass of 578.1427. Five peaks of the MS/MS spectra matched the predicted fragments of the molecule Procyanidin dimer B1. Procyanidin dimer B1 fragments at *m/z* 485.1245, 275.0545, 241.049, and 269.04 (Procyanidin B1 Mass Spectrum, n.d.) are the same as the fragments reported in the literature. Thus, the molecule is tentatively confirmed as Procyanidin dimer B1. 6UFS has a molecular mass of 564.11263. Eleven peaks of the MS/MS spectra matched the predicted fragments of the molecule Theaflavin. Theaflavin fragments at *m/z* 288.0622 (Verloop, n.d.) are consistent with the fragments reported in the literature. Additionally, some fragments for this molecule are observed at *m/z* 515.02986, 203.0709, 188.0461, and 92.9968, which show prominent spectra in MS/MS fragmentation. Thus, the molecule is tentatively confirmed as Theaflavin.

The theoretical molecular mass of 1UQS is 324.0992. Fourteen MS/MS spectra matched the predicted fragments of the molecule 6-O-desmethylterphenyllin. Fragments of 6-O-desmethylterphenyllin are observed at *m/z* 305.0821, 287.0696, 275.0706, and 262.0631. Thus, this molecule is tentatively confirmed as 6-O-desmethylterphenyllin. The theoretical molecular mass of 2UQS is 352.0941. Twelve fragments from the MS/MS spectra matched the predicted fragments of the molecule Altertoxin I. The fragments of Altertoxin I at *m/z* 333.0751 (Puntscher et al., 2019), 290.0572 (Stack et al., n.d.), matched those described in the literature, and some prominent fragments are observed at *m/z* 305.0821 and 262.0631 during the fragmentation. As a result, molecule I is tentatively confirmed as Altertoxin I. The theoretical molecular mass of 3UQS is 246.0895. The MS/MS spectra have fifteen peaks that correspond to predicted Altechromene B. A few of the Altechromene B fragments described in MetFrag are at *m/z* 230.0574, 188.0488, 160.0532, and 146.0376. Thus, the molecule is tentatively confirmed as Altechromene B. The theoretical molecular mass of 4UQS is 302.0786. Twenty-five MS/MS spectra match the predicted fragments of Botryorhodine F. Out of 25 fragments, a few of the Botryorhodine F fragments are observed at *m/z* 259.0955, 241.0871, 226.0638, and 217.0864. Thus, the molecule can be tentatively confirmed as Botryorhodine F. 5UQS has a theoretical molecular mass of 272.0684. Analysis of its MS/MS spectra revealed twenty fragments that matched those predicted for 3’,4’,7-Trihydroxyisoflavanone using MetFrag software. These matches align well with known fragmentation patterns for 3’,4’,7-Trihydroxyisoflavanone, including fragments at *m/z* 270.053, 257.047, 256.0358, and 242.0260 reported in the literature (Human Metabolome Database: LC-MS/MS Spectrum - ESI-TOF 30V, Negative (HMDB0041655), n.d.) Using ChemDraw, the structures of these fragments were analysed at *m/z* 271 [C_15_H_11_O_5_]-, 242 [C_14_H_10_O_4_]-, and 226 [C14H10O3]-. Their respective losses were calculated as [(M-H)]-, [(M-H)-CH_2_O]-, and [(M-H)-CH2O-OH]-. Based on the consistent fragmentation patterns and matches with MetFrag predictions, the molecule can be tentatively identified as 3’,4’,7-Trihydroxyisoflavanone.

The theoretical molecular mass of 6UQS matches that of alternariol 9-methyl ether is 272.0687. Fourteen MS/MS spectra matched the fragmentation pattern predicted for alternariol 9-methyl ether. Additionally, fragments identified in our spectra at *m/z* 257.0461 (Burkhardt et al., 2011), 242.0206 (23_Alternariol Monomethyl Ether and α, β-Dehydrocurvularin from Endophytic Fungi Alternaria spp. Inhibit Appressorium Formation of Magnaporthe griseajksabc.2010.007, n.d.), corresponded to those reported in the literature for alternariol 9-methyl ether. Furthermore, significant fragments were observed at *m/z* 271.061, 270.061, 242.0206 and 198.0338 during MS/MS fragmentation. Based on these consistent matches and fragmentation patterns, the identity of the molecule can be confirmed as alternariol 9-methyl ether. 7UQS exhibits a molecular mass of 302.0421. Notably, twelve of its MS/MS fragments matches those predicted for Morin. These matches align well with Morin fragments reported in literature, including those found at *m/z* 273.0399, 258.0159, 229.0145, and 189.0176 (McNab et al., 2009) Furthermore, analysis using ChemDraw software on a few additional fragments revealed structures at *m/z* 273 [C_14_H_9_O_7_]- and 258 [C_14_H_10_O_5_]-, corresponding to the loss of [(M-H)-CH_2_O]- and [(M-H)-CHO-OH]-, respectively. Based on these consistent fragmentation patterns, the molecule can be tentatively identified as Morin.

### 3.4 Molecular Docking

Docking was conducted for the twenty-six metabolites identified from EFS and EQS. Docking studies of these metabolites are based on their respective binding energies compared to that of standard drug acarbose. The details are provided in supplementary file 3 (A). Seven molecules unique to EQS exhibited significantly greater binding energy than acarbose. They are 6-O-desmethylterphenyllin (-6.9 kcal/mol), Altertoxin I (-7.8 kcal/mol), Altechromone B (-6.8 kcal/mol), Botryorhodine F (-6.7 kcal/mol), 3’,4’,7-Trihydroxyisoflavanone (-7.5 kcal/mol), alternariol 9-methyl ether (-7 kcal/mol), Morin (-7.5 kcal/mol), and acarbose (-6.6 kcal/mol). However, two molecules unique to EFS, *p*-Coumaric acid (-6.7 kcal/mol) and Diaportinol (-7 kcal/mol), showed greater binding energy than acarbose. Three molecules common to both EFS and EQS, 5’-Epialtenuen (-6.7 kcal/mol), 1,8-dihydroxynaphthalene (-6.7 kcal/mol), and 6-Epi-stemphytriol (-7.2 kcal/mol) displayed greater binding energy than acarbose. A 2D ligplot image of all 12 molecules is shown in supplementary file 3 (B).

### 3.5 Ensemble docking

3’,4’,7-Trihydroxyisoflavanone and alternariol 9-methyl ether were selected for Ensemble docking based on their higher binding energy compared to that of acarbose. Docking was carried out with nine different poses of 2QMJ obtained through simulation studies. In the case of 3’,4’,7-Trihydroxyisoflavanone, the first pose, labelled 2QMJ01_3’,4’,7-Trihydroxyisoflavanone, exhibited no hydrogen bonds. In the second pose, 2QMJ02_3’,4’,7-Trihydroxyisoflavanone, five hydrogen bonds were observed: Arg526, Asp571, Gly541, Trp539, and His600. The third pose, 2QMJ03_3’,4’,7-Trihydroxyisoflavanone, exhibited a single hydrogen bond with Tyr 299. The fourth pose, 2QMJ04_3’,4’,7-Trihydroxyisoflavanone, displayed three hydrogen bonds: Trp406 and Asp203 (2). In the fifth pose, 2QMJ05_3’,4’,7-Trihydroxyisoflavanone, three hydrogen bonds were observed with Arg334, Asp366, and Trp406. The sixth pose, 2QMJ06_3’,4’,7-Trihydroxyisoflavanone, demonstrated a total of three hydrogen bonds involving Asp571 (2) and His600. The seventh pose, 2QMJ07_3’,4’,7-Trihydroxyisoflavanone, exhibited one hydrogen bond with His600. The eighth pose, 2QMJ08_3’,4’,7-Trihydroxyisoflavanone, displayed one hydrogen bond: Asp542. The ninth pose, 2QMJ09_3’,4’,7-Trihydroxyisoflavanone, exhibited three hydrogen bonds involving Trp441 and Arg598 (2). All nine poses are displayed in the supplementary file 3 (C). A few amino acids like Asp443 and Met444, and Trp539 were observed across different poses.

Ensemble docking was performed for the nine different poses of 2QMJ with alternariol 9-methyl ether as the ligand. In the first pose, labelled 2QMJ01_alternariol 9-methyl ether, no hydrogen bonds were observed. In the second pose, 2QMJ02_alternariol 9-methyl ether, three hydrogen bonds were observed: Trp539, His600 (2). The third pose, 2QMJ03_alternariol 9-methyl ether, exhibited one hydrogen bond with Asp329. The fourth pose, 2QMJ04_alternariol 9-methyl ether, displayed four hydrogen bonds: Arg526, Asp571, Trp539, His600. In the fifth pose, 2QMJ05_alternariol 9-methyl ether, four hydrogen bonds were observed: Tyr299, Trp406, Arg335, Arg334. The sixth pose, 2QMJ06_ alternariol 9-methyl ether, demonstrated a total of two hydrogen bonds involving His600 (2). The seventh pose, 2QMJ07_ alternariol 9-methyl ether, exhibited two hydrogen bonds with His600 (2). The eighth pose, 2QMJ08_ alternariol 9-methyl ether, displayed two hydrogen bonds: Asp327, Trp406. The ninth pose, 2QMJ09_ alternariol 9-methyl ether, exhibited two hydrogen bonds involving Asp327, His600. Amnio acid like Asp 327 was observed across different poses. All the nine poses 2D images are shown in the supplementary file 3 (D).

## 4. Discussion

Enzyme α-glucosidase breaks down complex carbohydrates into simple sugars in the intestine, which are readily absorbed into the blood stream resulting in a sudden spike in blood glucose levels after a meal. Therefore, inhibiting α-glucosidase activity is a key strategy for managing diabetes. While α-glucosidase inhibitor drugs currently available in the market control sugar levels, they often produce some side effects as well. Bioactive molecules from endophytic fungi hold promise as effective α-glucosidase inhibitors with minimal or no side effects for lowering postprandial blood glucose levels in diabetic patients (Naveen and Baskaran, 2018). Phenolic compounds produced by endophytic fungi through metabolic pathways exhibit both antioxidant properties as well as α-glucosidase enzyme inhibitory effects. Because of their unique chemical and structural properties, these phenolic compounds present themselves as potential molecules with α-glucosidase inhibiting activity(Akhtar et al., 2018).

Nutrient availability in the culture medium in which endophytic fungi are grown has a profound impact on their growth and metabolism. Alteration in media composition brings about differential activation of BGCs leading to synthesis of certain set of secondary metabolites. Partial starvation or limited nutrient availability occasionally leads to activation of specific BGCs resulting in synthesis of a set of metabolites with potential therapeutic value (Pan et al., 2019). In our study, metabolites from EQS exhibited significantly higher IC_50_ values for α-glucosidase inhibition as compared to those from EFS. Further, EQS also exhibited higher phenolic content as compared to EFS.

Bioactivity-guided fractionation is a vital method for isolating and studying bioactive compounds from fungal extracts. While fractionation is lengthy, time-consuming and challenging due to low concentrations of bioactive compounds in the extracts, dereplication offers a quicker alternative for tentative identification of molecules based on those already reported, without having to actually isolate them. Our study employed dereplication approach involving UHPLC-ESI-QTOF-MS for tentative identification of molecules. LC-MS, being highly sensitive can tentatively identify metabolites during the first fractionation step itself and there by facilitate targeted fractionation of the desired molecules. (Ito and Masubuchi, 2014)

Docking studies were carried out for all tentatively identified molecules using AutoDock vina software. Based on the free energy and hydrogen bonding interactions, twelve molecules form EFS and EQS together showed greater binding affinity for the active site of α-glucosidase compared to that of acarbose (Table 3). On the basis of available reports, six molecules (Morin (4.48 ± 0.04 μM) (Zeng et al., 2016), Botryorhodine F (12.01 ± 1.2 μg/mL) (Chen et al., 2015), 6’-O-Desmethylterphenyllin (0.9 μM) (Huang et al., 2011), and Alternarienonic acid (7.95± 1.2µM) (Elbermawi et al., 2022), *p*-Coumaric acid (99.8±0.2 µM) (Takahashi and Miyazawa, 2012)) out of twenty-six tentatively identified molecules from EQS and EFS showed lower IC_50_ values, indicating their significant α-glucosidase inhibition activity.

**Table 3.** Twelve molecules identified from EFS and EQS exhibited greater α-glucosidase inhibitory activity compared to acarbose.

| Code | Compound Name | Best pose<br>Binding energy<br>(kcal/mol) | RMSD | H-bond interaction | No. of Hydrogen<br>Bonds |
| --- | --- | --- | --- | --- | --- |
| 2UQS | Altertoxin I | -7.8 | 4.518 | Asp203 | 1 |
| 5UQS | 3',4',7-Trihydroxyisoflavanone | -7.5 | 1.811 | Arg526, Asp571,<br>Gly541, Trp539,<br>His600 | 5 |
| 7UQS | Morin | -7.5 | 2.805 | Asp327, Asp542,<br>Arg526, Asp203 | 4 |
| 9CFS &<br>9CQS | 6-Epi-stemphytriol | -7.2 | 1.544 | Arg334(2), Glu404 | 2 |
| 6UQS | Alternariol 9-methyl ether | -7 | 3.741 | Arg526, Asp571,<br>Trp539, His600 | 4 |
| 3UFS | Diaportinol | -7 | 1.569 | Asp327, His600,<br>Asp542, Arg526 | 4 |
| 1UQS | 6'-O-Desmethylterphenyllin | -6.9 | 1.443 | Asp327, His600 | 2 |
| 3UQS | Altechromone B | -6.8 | 2.083 | Arg526 | 1 |
| 4UQS | Botryorhodine F | -6.7 | 1.521 | Tyr605, Gln603,<br>Asp203 | 3 |
| 2UFS | *p*-Coumaric acid | -6.7 | 0.563 | Asn543, Ser553,<br>Asp549 | 3 |
| 2CFS &<br>2CQS | 1,8-ihydroxynaphthalene | -6.7 | 0.093 | Asp443, Arg526 | 3 |
| 7CFS &<br>7CQS | 5'-Epialtenuen | -6.7 | 1.757 | Met444, Asp443,<br>Arg526 | 3 |
| Positive<br>Ctrl | Acarbose | -6.6 | 4.668 | Trp406, Asp443,<br>Asp327, His600,<br>Arg334 | 5 |

For the first time we report docking studies for twenty-six molecules (Supplementary file 3 (A)) with 2QMJ subunit of α-glucosidase. Among these twelve molecules, seven from EQS and five from EFS exhibited greater binding energies than acarbose. Since 3’,4’,7-Trihydroxyisoflavanone and alternariol 9-methyl ether have high binding energies of -7.5 kcal/mol and -7.0 kcal/mol respectively, they could be acting synergistically along with the other molecules from EQS having high binding energies in contributing to α-glucosidase inhibiting activity. However, their IC_50_ values for α-glucosidase inhibition are yet to be worked out.

Ensemble docking, a computational approach employed in drug discovery, utilizes a range of target conformations accommodating protein flexibility via molecular dynamics simulations. It accurately predicts ligand binding and helps in the process of drug discovery for diseases like diabetes (Amaro et al., 2018). In this study two molecules from EQS were taken for ensemble docking to gain deeper insight into their interactions with the active site. Molecules 3’,4’,7-Trihydroxyisoflavanone and alternariol 9-methyl ether were found to have stable hydrogen bonding interactions across all the poses, especially with His 600, a highly conserved residue of GH31 family (Sim et al., 2008) that includes α-glucosidase (Figure 4). 3’,4’,7-Trihydroxyisoflavanone and alternariol 9-methyl ether are being reported for the first time from *Alternaria* species. Based on docking studies, we propose that 3’,4’,7-Trihydroxyisoflavanone and alternariol 9-methyl ether are potential therapeutic molecules for treating diabetes.

**Figure 4.**
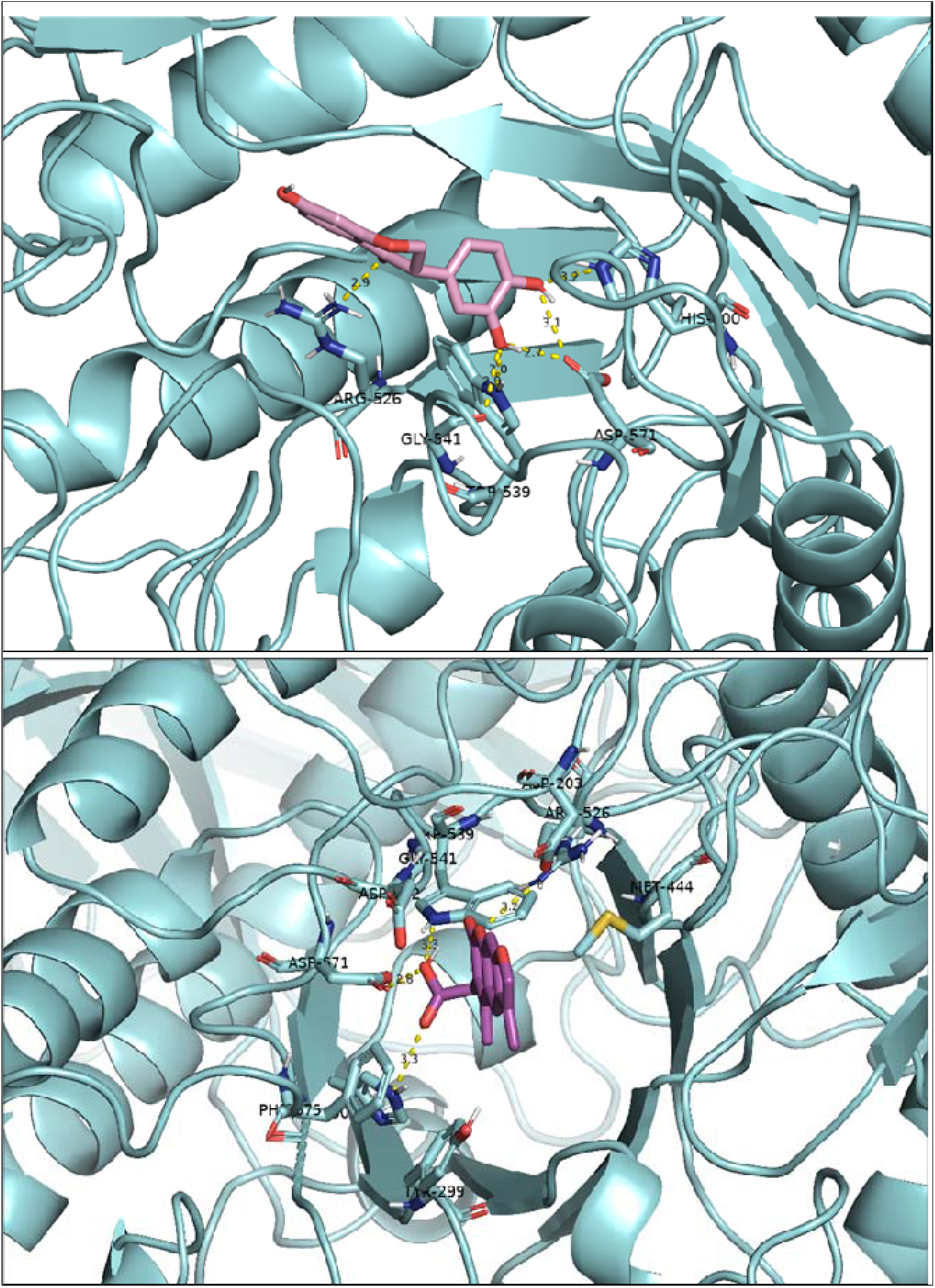
Docking results for two ligands: 3’,4’,7-Trihydroxyisoflavanone at pose 2 (top) and Alternariol 9-Methyl Ether at pose 4 (bottom). The protein is represented in cyan, and hydrogen bonds between the ligands and the protein’s amino acids are shown in yellow. The figure was generated using PyMOL software.

The key finding of this study is that *Alternaria sp.* grown in EQS exhibited greater potential for producing bioactive secondary metabolites compared to those grown in EFS, highlighting the importance of nutrient concentration in modulating fungal metabolite production. Secondly, *Alternaria* sp. cultured in the EQS are a good source of 3’,4’,7-Trihydroxyisoflavanone and alternariol 9-methyl ether. Thirdly this study unveils several previously unknown metabolites from *Alternaria* species, expanding our knowledge and understanding of fungal metabolism and opening more avenues for bioactive compound discovery. Moving forward, we aim to undertake large-scale cultivation of the fungi, purification of the desired metabolites, and NMR-based characterization.

## 5. Conclusion

In conclusion, this study highlights the potential of *Alternaria* sp. BRN05, isolated from *Swietenia macrophylla* King, is a source of novel bioactive molecules with α-glucosidase inhibitory activity. Cultivation in quarter-strength media significantly enhances α-glucosidase inhibitory activity compared to full-strength media. Metabolic profiling identified 19 molecules from EFS and 20 molecules from EQS. Molecular docking reveals that 12 molecules exhibit superior binding energies compared to acarbose. Notably, ensemble docking of 3’,4’,7-Trihydroxyisoflavanone and alternariol 9-methyl ether from EQS demonstrates stable interactions with the active site, confirming their potential as α-glucosidase inhibitors. Thus, these molecules can be potential therapeutic agents that are safer and more effective in the treatment of diabetes.

## 7. Author Contributions

Piyush Kumar - Method development, Performing the experiment, Writing - Original Draft, Data analysis and presentation. Sai Anand Kannakazhi Kantari **-** Conception or design of the work, analysis and presentation. Ranendra Pratap Biswal - Method development for LC-MS analysis. Prasanth Ghanta - Application of computational analysis; Malleswara Dharanikota - Writing - Review & Editing, Supervision.

## Acknowledgments

The authors are ever grateful to the Founder Chancellor, Sri Sathya Sai Institute of Higher Learning (SSSIHL), for his constant inspiration. The authors are thankful to SSSIHL and UGC-SAP for providing financial support to carry out this work. The authors are grateful to Central Research Instrumentation Facility (CRIF), SSSIHL, for providing all necessary facilities. The authors wish to thank V. N. Ravi Kishore Vutukuri for help provided in optimizing mass spectrometry protocols and A Ashok for guidance and help in carrying out this work.

## 9. Supplementary Material

Figure (Supplementary 1), Figure (Supplementary 2), Figure (Supplementary 3) and Table (Supplementary 3).

## Reference styles

Alternariol Monomethyl Ether and α, β-Dehydrocurvularin from Endophytic Fungi Alternaria spp. Inhibit Appressorium Formation of Magnaporthe griseajksabc.2010.007 (n.d.).

Akhtar, N., Ihsan-ul-Haq, and Mirza, B. (2018). Phytochemical analysis and comprehensive evaluation of antimicrobial and antioxidant properties of 61 medicinal plant species. Arabian Journal of Chemistry 11, 1223–1235. doi: 10.1016/j.arabjc.2015.01.013

Allen, F., Greiner, R., and Wishart, D. (2015). Competitive fragmentation modeling of ESI-MS/MS spectra for putative metabolite identification. Metabolomics 11, 98–110. doi: 10.1007/S11306-014-0676-4

Al-Nema, M., Gaurav, A., and Lee, V. S. (2023). Designing of 2,3-dihydrobenzofuran derivatives as inhibitors of PDE1B using pharmacophore screening, ensemble docking and molecular dynamics approach. Comput Biol Med 159. doi: 10.1016/j.compbiomed.2023.106869

Amaro, R. E., Baudry, J., Chodera, J., Demir, Ö., McCammon, J. A., Miao, Y., et al. (2018). Ensemble Docking in Drug Discovery. Biophys J 114, 2271–2278. doi: 10.1016/j.bpj.2018.02.038

Bhatia, A., Singh, B., Arora, R., and Arora, S. (2019). In vitro evaluation of the α-glucosidase inhibitory potential of methanolic extracts of traditionally used antidiabetic plants. BMC Complement Altern Med 19. doi: 10.1186/s12906-019-2482-z

Biswal, R. P., Dandamudi, R. B., Patnana, D. P., Pandey, M., and Vutukuri, V. N. R. K. (2022). Metabolic fingerprinting of Ganoderma spp. using UHPLC-ESI-QTOF-MS and its chemometric analysis. Phytochemistry 199, 113169. doi: 10.1016/J.PHYTOCHEM.2022.113169

Brás, N. F., Santos-Martins, D., Fernandes, P. A., and Ramos, M. J. (2018). Mechanistic Pathway on Human α-Glucosidase Maltase-Glucoamylase Unveiled by QM/MM Calculations. Journal of Physical Chemistry B 122, 3889–3899. doi: 10.1021/acs.jpcb.8b01321

Burkhardt, B., Wittenauer, J., Pfeiffer, E., Schauer, U. M. D., and Metzler, M. (2011). Oxidative metabolism of the mycotoxins alternariol and alternariol-9-methyl ether in precision-cut rat liver slices in vitro. Mol Nutr Food Res 55, 1079–1086. doi: 10.1002/mnfr.201000487

Chen, S., Liu, Z., Liu, Y., Lu, Y., He, L., and She, Z. (2015). New depsidones and isoindolinones from the mangrove endophytic fungus Meyerozyma guilliermondii (HZ-Y2) isolated from the South China Sea. Beilstein Journal of Organic Chemistry 11, 1187–1193. doi: 10.3762/bjoc.11.133

Cho, N. H., Shaw, J. E., Karuranga, S., Huang, Y., da Rocha Fernandes, J. D., Ohlrogge, A. W., et al. (2018). IDF Diabetes Atlas: Global estimates of diabetes prevalence for 2017 and projections for 2045. Diabetes Res Clin Pract 138, 271–281. doi: 10.1016/j.diabres.2018.02.023

Da Silva, M. N., Arruda, M. S. P., Castro, K. C. F., Da Silva, M. F. D. G. F., Fernandes, J. B., and Vieira, P. C. (2008). Limonoids of the phragmalin type from Swietenia macrophylla and their chemotaxonomic significance. J Nat Prod 71, 1983–1987. doi: 10.1021/np800312h

Elbermawi, A., Ali, A. R., Amen, Y., Ashour, A., Ahmad, K. F., Mansour, E. S. S., et al. (2022). Anti-diabetic activities of phenolic compounds of Alternaria sp., an endophyte isolated from the leaves of desert plants growing in Egypt. RSC Adv 12, 24935–24945. doi: 10.1039/d2ra02532a

Elferink, H., Bruekers, J. P. J., Veeneman, G. H., and Boltje, T. J. (2020). A comprehensive overview of substrate specificity of glycoside hydrolases and transporters in the small intestine: “A gut feeling.” Cellular and Molecular Life Sciences 77, 4799–4826. doi: 10.1007/s00018-020-03564-1

Eram, D., Arthikala, M., Melappa, G., Santoyo, G., Eram, D., Arthikala, M., et al. (2018). Alternaria species: endophytic fungi as alternative sources of bioactive compounds. Ital J Mycol 47. doi: 10.6092/issn.2531-7342/8468

Ghanta, P., Sinha, S., Doble, M., and Ramaiah, B. (2022). Potential of pyrroquinazoline alkaloids from Adhatoda vasica Nees. as inhibitors of 5-LOX–a computational and an in-vitro study. J Biomol Struct Dyn 40, 2785–2796. doi: 10.1080/07391102.2020.1848635

Ghorbani, H., Ebadi, A., Faramarzi, M. A., Mojtabavi, S., Mahdavi, M., and Najafi, Z. (2023). Synthesis, in vitro α-glucosidase inhibitory activity and molecular dynamics simulation of some new coumarin-fused 4H-pyran derivatives as potential anti-diabetic agents. J Mol Struct 1284. doi: 10.1016/j.molstruc.2023.135349

Huang, H., Feng, X., Xiao, Z., Liu, L., Li, H., Ma, L., et al. (2011). Azaphilones and p-terphenyls from the mangrove endophytic fungus Penicillium chermesinum (ZH4-E2) isolated from the south China sea. J Nat Prod 74, 997–1002. doi: 10.1021/np100889v

Huang, S. Y., and Zou, X. (2007). Ensemble docking of multiple protein structures: Considering protein structural variations in molecular docking. Proteins: Structure, Function and Genetics 66, 399–421. doi: 10.1002/prot.21214

Human Metabolome Database: LC-MS/MS Spectrum - ESI-TOF 30 V, Negative (HMDB0041655) (n.d.). Available at: https://hmdb.ca/spectra/ms_ms/373881 (Accessed November 29, 2023).

insight review articles 782 (2001). Available at: www.nature.com

Ito, T., and Masubuchi, M. (2014). Dereplication of microbial extracts and related analytical technologies. Journal of Antibiotics 67, 353–360. doi: 10.1038/ja.2014.12

Jaiswal, N. (2013). Inhibition of Alpha-Glucosidase by Acacia nilotica Prevents Hyperglycemia along with Improvement of Diabetic Complications via Aldose Reductase Inhibition. J Diabetes Metab 01. doi: 10.4172/2155-6156.s6-004

Jiao, Y., Hua, D., Huang, D., Zhang, Q., and Yan, C. (2018). Characterization of a new heteropolysaccharide from green guava and its application as an α-glucosidase inhibitor for the treatment of type II diabetes. Food Funct 9, 3997–4007. doi: 10.1039/c8fo00790j

Kashtoh, H., and Baek, K. H. (2022). Recent Updates on Phytoconstituent Alpha-Glucosidase Inhibitors: An Approach towards the Treatment of Type Two Diabetes. Plants 11. doi: 10.3390/plants11202722

Keller, N. P. (2019). Fungal secondary metabolism: regulation, function and drug discovery. Nat Rev Microbiol 17, 167–180. doi: 10.1038/s41579-018-0121-1

Kim, K.-H., Lawrence, C. B., Mcdowell, J. B., Li, L., and Tholl, D. (2009). Functional Analysis of Secondary Metabolite Biosynthesis-Related Genes in Alternaria brassicicola.

Kjer, J. (2009). New Natural Products from Endophytic Fungi from Mangrove Plants-Structure Elucidation and Biological Screening Neue Naturstoffe aus endophytischen Pilzen aus Mangroven-Strukturaufklärung und Evaluierung der biologischen Aktivität.

Kumar, S., Stecher, G., Li, M., Knyaz, C., and Tamura, K. (2018). MEGA X: Molecular evolutionary genetics analysis across computing platforms. Mol Biol Evol 35, 1547–1549. doi: 10.1093/molbev/msy096

Laoud, A., Ferkous, F., Maccari, L., Maccari, G., Saihi, Y., and Kraim, K. (2018). Identification of novel nt-MGAM inhibitors for potential treatment of type 2 diabetes: Virtual screening, atom based 3D-QSAR model, docking analysis and ADME study. Comput Biol Chem 72, 122–135. doi: 10.1016/j.compbiolchem.2017.12.003

LC-MS/MS Spectrum - LC-ESI-QTOF (UPLC Q-Tof Premier, Waters), Negative (DB04066) | DrugBank Online (n.d.). Available at: https://go.drugbank.com/spectra/ms_ms/6493 (Accessed November 28, 2023).

McNab, H., Ferreira, E. S. B., Hulme, A. N., and Quye, A. (2009). Negative ion ESI-MS analysis of natural yellow dye flavonoids-An isotopic labelling study. Int J Mass Spectrom 284, 57–65. doi: 10.1016/j.ijms.2008.05.039

Mohammed, A. E., Sonbol, H., Alwakeel, S. S., Alotaibi, M. O., Alotaibi, S., Alothman, N., et al. (2021). Investigation of biological activity of soil fungal extracts and LC/MS-QTOF based metabolite profiling. Sci Rep 11. doi: 10.1038/s41598-021-83556-8

Naveen, J., and Baskaran, V. (2018). Antidiabetic plant-derived nutraceuticals: a critical review. Eur J Nutr 57, 1275–1299. doi: 10.1007/s00394-017-1552-6

Pan, R., Bai, X., Chen, J., Zhang, H., and Wang, H. (2019). Exploring structural diversity of microbe secondary metabolites using OSMAC strategy: A literature review. Front Microbiol 10. doi: 10.3389/fmicb.2019.00294

Patil, P., Mandal, S., Tomar, S. K., and Anand, S. (2015). Food protein-derived bioactive peptides in management of type 2 diabetes. Eur J Nutr 54, 863–880. doi: 10.1007/s00394-015-0974-2

Phenylacetic acid Mass Spectrum (n.d.). Available at: https://massbank.eu/MassBank/RecordDisplay?id=MSBNK-Metabolon-MT000056 (Accessed November 28, 2023).

Procyanidin B1 Mass Spectrum (n.d.). Available at: https://massbank.eu/MassBank/RecordDisplay?id=MSBNK-BS-BS003948 (Accessed November 28, 2023).

Puntscher, H., Marko, D., and Warth, B. (2019). The fate of altertoxin ii during tomato processing steps at a laboratory scale. Front Nutr 6. doi: 10.3389/fnut.2019.00092

Ruttkies, C., Schymanski, E. L., Wolf, S., Hollender, J., and Neumann, S. (2016). MetFrag relaunched: Incorporating strategies beyond in silico fragmentation. J Cheminform 8. doi: 10.1186/s13321-016-0115-9

Sabreen, A. K., Lena, F. H., and Imad, H. H. (2015). Antibacterial activity of secondary metabolites isolated from Alternaria alternata. Afr J Biotechnol 14, 2972–2994. doi: 10.5897/ajb2015.14906

Seeliger, D., and De Groot, B. L. (2010). Conformational transitions upon ligand binding: Holo-structure prediction from apo conformations. PLoS Comput Biol 6. doi: 10.1371/journal.pcbi.1000634

Sim, L., Quezada-Calvillo, R., Sterchi, E. E., Nichols, B. L., and Rose, D. R. (2008). Human Intestinal Maltase-Glucoamylase: Crystal Structure of the N-Terminal Catalytic Subunit and Basis of Inhibition and Substrate Specificity. J Mol Biol 375, 782–792. doi: 10.1016/j.jmb.2007.10.069

Stack, M. E., Mazzola, E. P., Page, S. W., Pohland, A. E., Highet, R. J., Tempesta, M. S., et al. (n.d.). MUTAGENIC PERYLENEQUINONE METABOLITES OF ALTERNARIA ALTERNATA: ALTERTOXINS I, 11, AND 111’.

Sukardiman, and Ervina, M. (2020). The recent use of Swietenia mahagoni (L.) Jacq. as antidiabetes type 2 phytomedicine: A systematic review. Heliyon 6. doi: 10.1016/j.heliyon.2020.e03536

Takahashi, T., and Miyazawa, M. (2012). Potent α-glucosidase inhibitors from safflower (Carthamus tinctorius L.) seed. Phytotherapy Research 26, 722–726. doi: 10.1002/ptr.3622

Tamura, K., Nei, M., and Kumar, S. (2004). Prospects for inferring very large phylogenies by using the neighbor-joining method. Available at: www.pnas.orgcgidoi10.1073pnas.0404206101

Tao, J., Bai, X., Zeng, M., Li, M., Hu, Z., Hua, Y., et al. (2022). Whole-Genome Sequence Analysis of an Endophytic Fungus Alternaria sp. SPS-2 and Its Biosynthetic Potential of Bioactive Secondary Metabolites. Microorganisms 10. doi: 10.3390/microorganisms10091789

Tomm, H. A., Ucciferri, L., and Ross, A. C. (2019). Advances in microbial culturing conditions to activate silent biosynthetic gene clusters for novel metabolite production. J Ind Microbiol Biotechnol 46, 1381–1400. doi: 10.1007/s10295-019-02198-y

Trott, O., and Olson, A. J. (2010). AutoDock Vina: Improving the speed and accuracy of docking with a new scoring function, efficient optimization, and multithreading. J Comput Chem 31, 455–461. doi: 10.1002/jcc.21334

Tseng, M. N., Chung, C. L., and Tzean, S. S. (2014). Mechanisms relevant to the enhanced virulence of a dihydroxynaphthalene-melanin metabolically engineered entomopathogen. PLoS One 9. doi: 10.1371/journal.pone.0090473

Verloop, A. J. W. (n.d.). Oligomerization and hydroxylation of green tea catechins by oxidative enzymes.

Zeng, L., Zhang, G., Liao, Y., and Gong, D. (2016). Inhibitory mechanism of morin on α-glucosidase and its anti-glycation properties. Food Funct 7, 3953–3963. doi: 10.1039/c6fo00680a

Zhao, S., Wang, B., Tian, K., Ji, W., Zhang, T., Ping, C., et al. (2021). Novel metabolites from the Cercis chinensis derived endophytic fungus Alternaria alternata ZHJG5 and their antibacterial activities. Pest Manag Sci 77, 2264–2271. doi: 10.1002/ps.6251

