## Supplementry Material for "Elucidation of α-glucosidase inhibitory activity and UHPLC-ESI-QTOF-MS based metabolic profiling of endophytic fungi *Alternaria* sp. BRN05. isolated from seeds of *Swietenia macrophylla* King": Supplementry Material 1.pdf

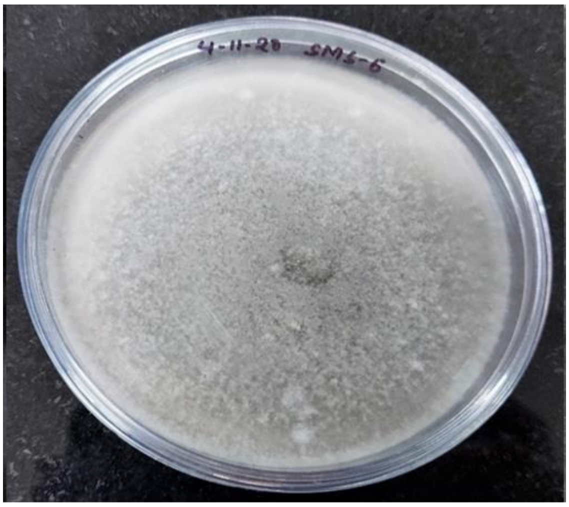


Supplementary file 1 (A). Endophytic fungi isolated from the seeds of *Swietenia macrophylla* cultured on PDA.

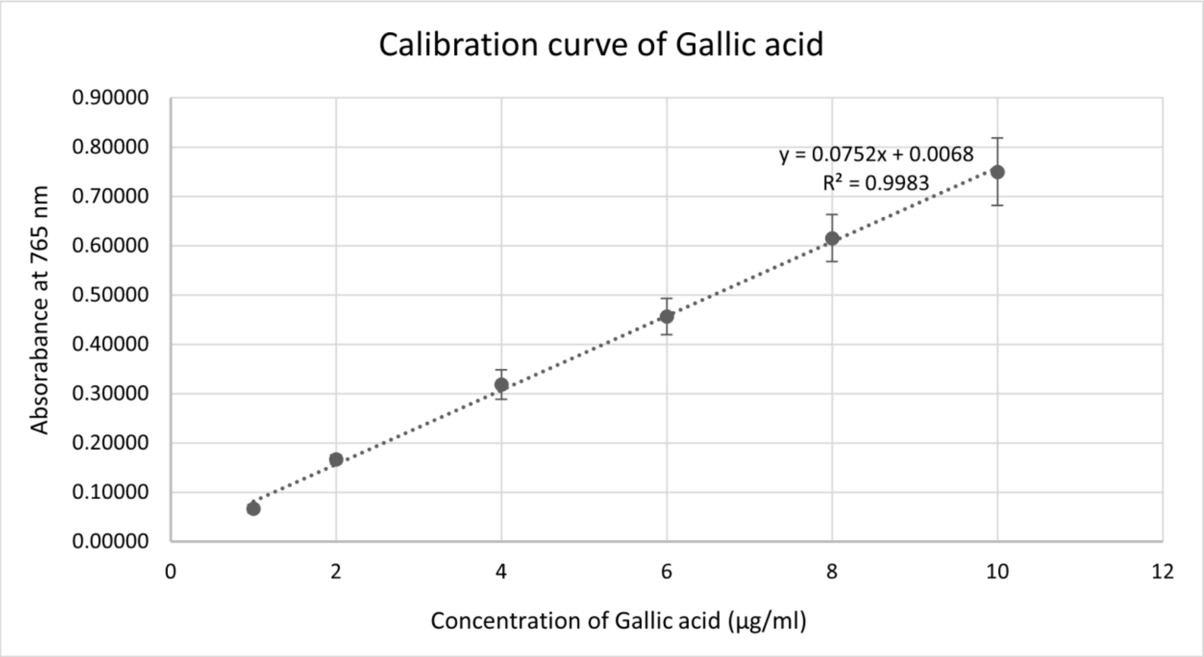


Supplementary file 1 (B). Calibration curve for Gallic acid expressed in gallic acid equivalents per gram of dry extract weight
