## Supplementry Material for "Elucidation of α-glucosidase inhibitory activity and UHPLC-ESI-QTOF-MS based metabolic profiling of endophytic fungi *Alternaria* sp. BRN05. isolated from seeds of *Swietenia macrophylla* King": Supplementry Material 2.pdf

### Supplementary File

- MS/MS spectra for the 39 compounds, along with fragments generated from Metfrag, as well as those analysed using ChemDraw Pro 8.0 software.

- In the Common Full-strength mode (CFS), MS/MS spectra of 13 molecules were generated

### 4-Hydroxymellein(1CFS) _12.953

| | Peak m/z | Formula |
|---|---|---|
| m/z | 221.0456 | (M+HCOO)- [-$H_2O$] |
| Molecular Mass | 194.05794 | $C_{10}H_{10}O_4$ |
| Fragment 1 | 177.0561 | $[C_9H_5O_4]^-$ |
| Fragment 2 | 175.04 | $[C_{10}H_7O_3]^-$ |
| Fragment 3 | 162.8385 | $[C_9H_6O_3]^-$ |
| Fragment 4 | 134.8639 | $[C_8H_7O_2]^-$ |

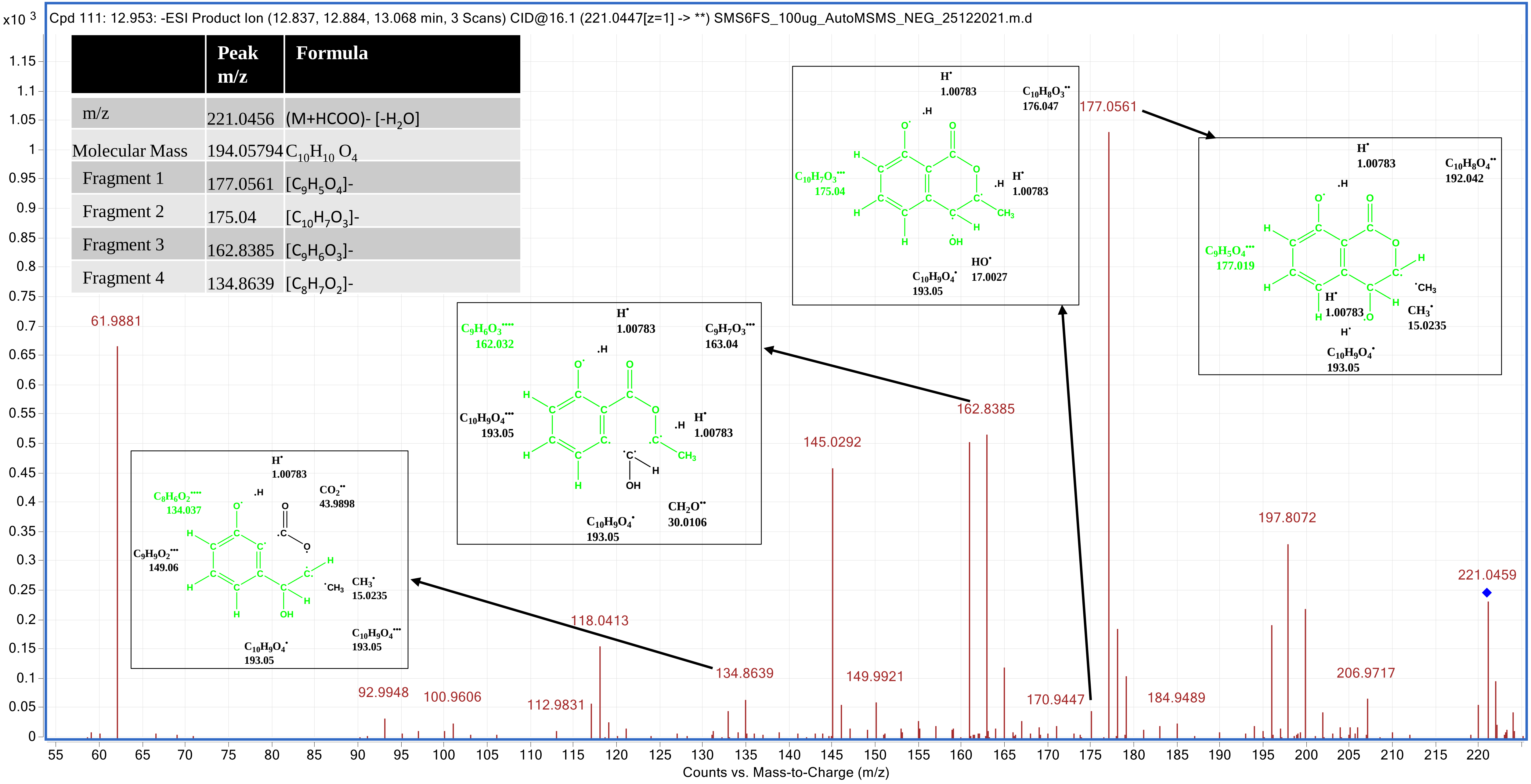

### 5'-Epialtenuene (2CFS) _12.337

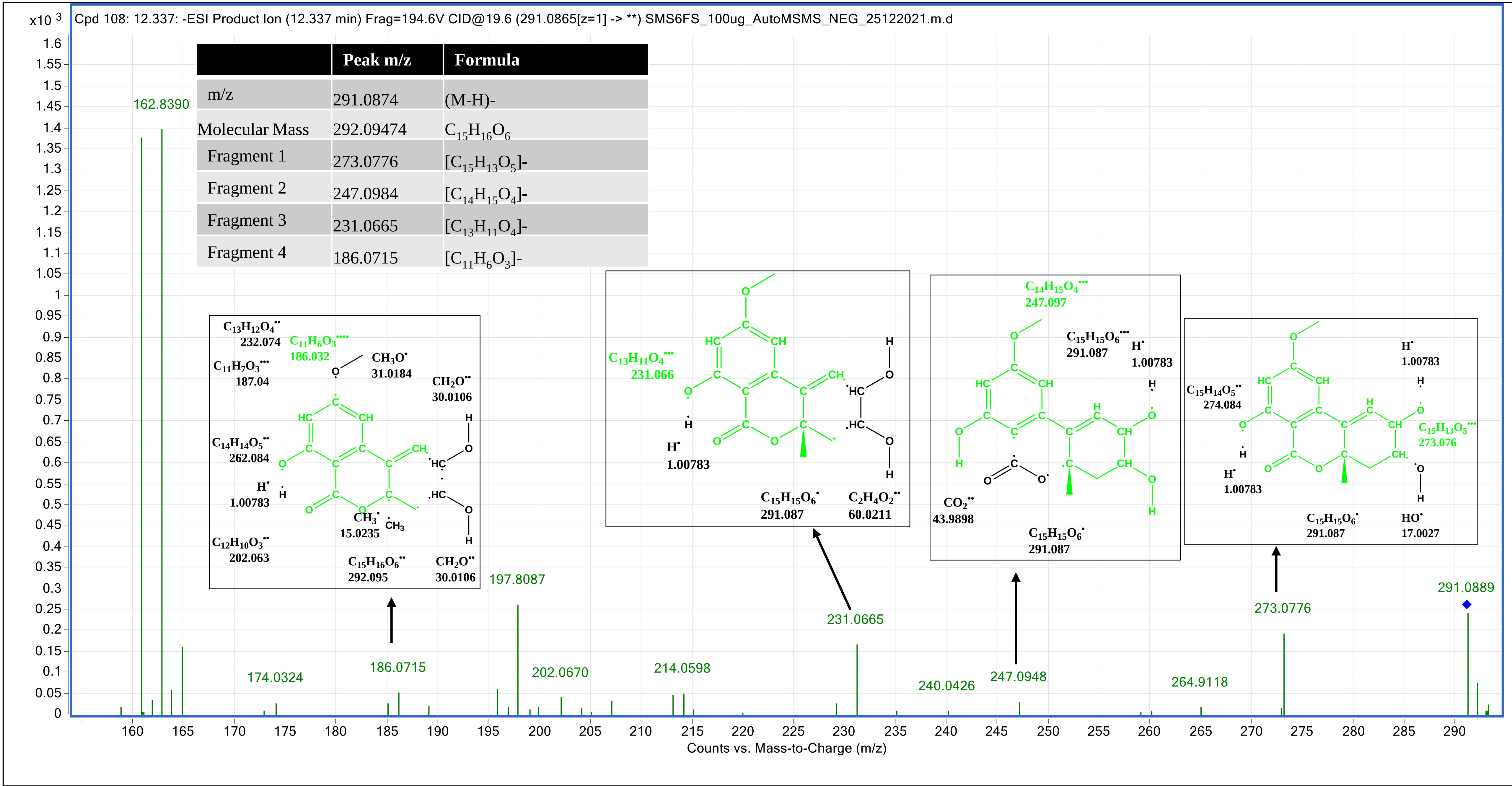


| | Peak m/z | Formula |
|---|---|---|
| m/z | 291.0874 | (M-H)- |
| Molecular Mass | 292.09474 | $C_{15}H_{16}O_6$ |
| Fragment 1 | 273.0776 | $[C_{15}H_{13}O_5]^-$ |
| Fragment 2 | 247.0984 | $[C_{14}H_{15}O_4]^-$ |
| Fragment 3 | 231.0665 | $[C_{13}H_{11}O_4]^-$ |
| Fragment 4 | 186.0715 | $[C_{11}H_6O_3]^-$ |

### Alternarienonic acid (3CFS) _7.41

| | Peak m/z | Formula |
|---|---|---|
| m/z | 277.07908 | (M-H)- |
| Molecular Mass | 278.0792 | $C_{14}H_{14}O_6$ |
| Fragment 1 | 277.0716 | $[C_{14}H_{12}O_6]$- |
| Fragment 2 | 246.9260 | $[C_{13}H_{10}O_5]$- |
| Fragment 3 | 233.0813 | $[C_{13}H_{13}O_4]$- |
| Fragment 4 | 214.9257 | $[C_{13}H_{10}O_3]$- |
| Fragment 5 | 189.0542 | $[C_{11}H_9O_3]$- |
| Fragment 6 | 251.0563 | $[C_{12}H_{11}O_6]$- |

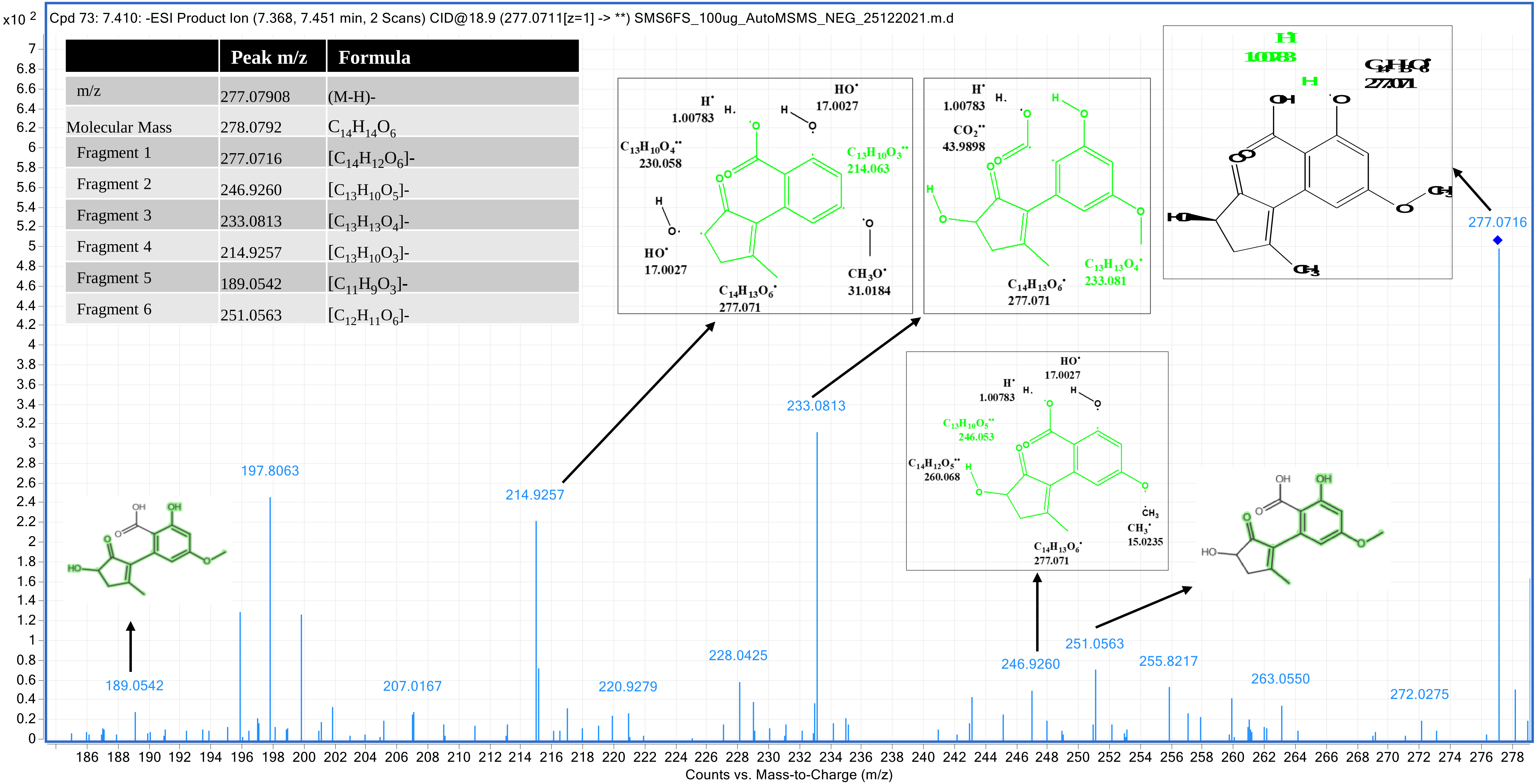

### (+)-talaroflavone (4CFS) _15.758

| | Peak m/z | Formula |
|---|---|---|
| m/z | 257.0462 | (M-H)- [-$H_2O$] |
| Molecular Mass | 276.06342 | $C_{14}H_{12}O_6$ |
| Fragment 1 | 257.0461 | [$C_{14}H_{10}O_5$]- |
| Fragment 2 | 246.9222 | [$C_{13}H_{11}O_5$]- |
| Fragment 3 | 229.0516 | [$C_{13}H_{11}O_4$]- |
| Fragment 4 | 213.0548 | [$C_{13}H_{11}O_3$]- |

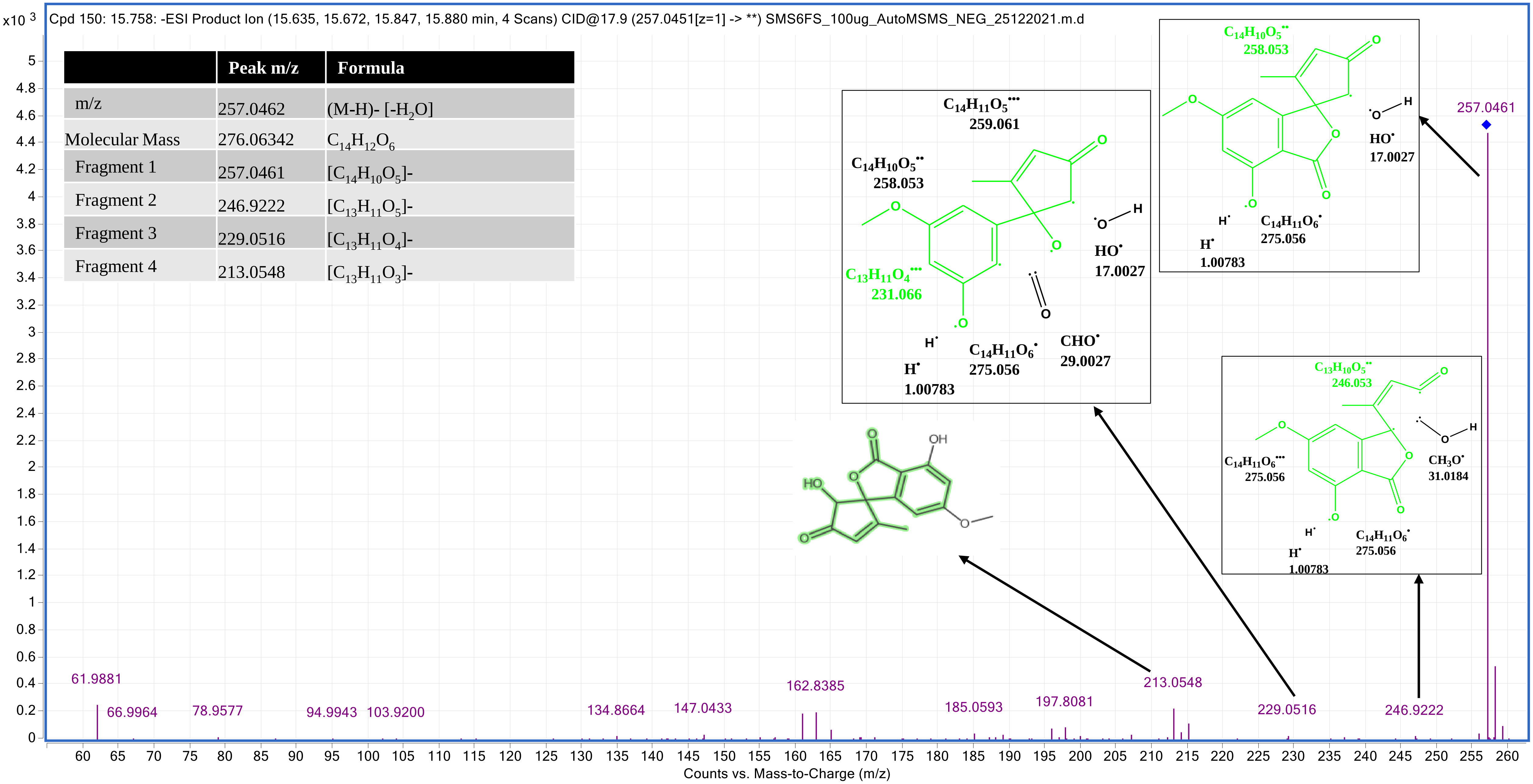

### 2,5-dimethyl-7-hydroxychromone (5CFS) _10.019

| | Peak m/z | Formula |
|---|---|---|
| m/z | 189.0567 | (M-H)- |
| Molecular Mass | 190.06303 | $C_{11}H_{10}O_3$ |
| Fragment 1 | 189.0567 | $[C_{11}H_9O_3]$- |
| Fragment 2 | 174.0324 | $[C_{10}H_6O_3]$- |
| Fragment 3 | 160.08424 | $[C_9H_4O_3]$- |
| Fragment 4 | 119.0491 | $[C_7H_3O_2]$- |
| Fragment 5 | 133.0646 | $[C_9H_8O_2 +H]$- |

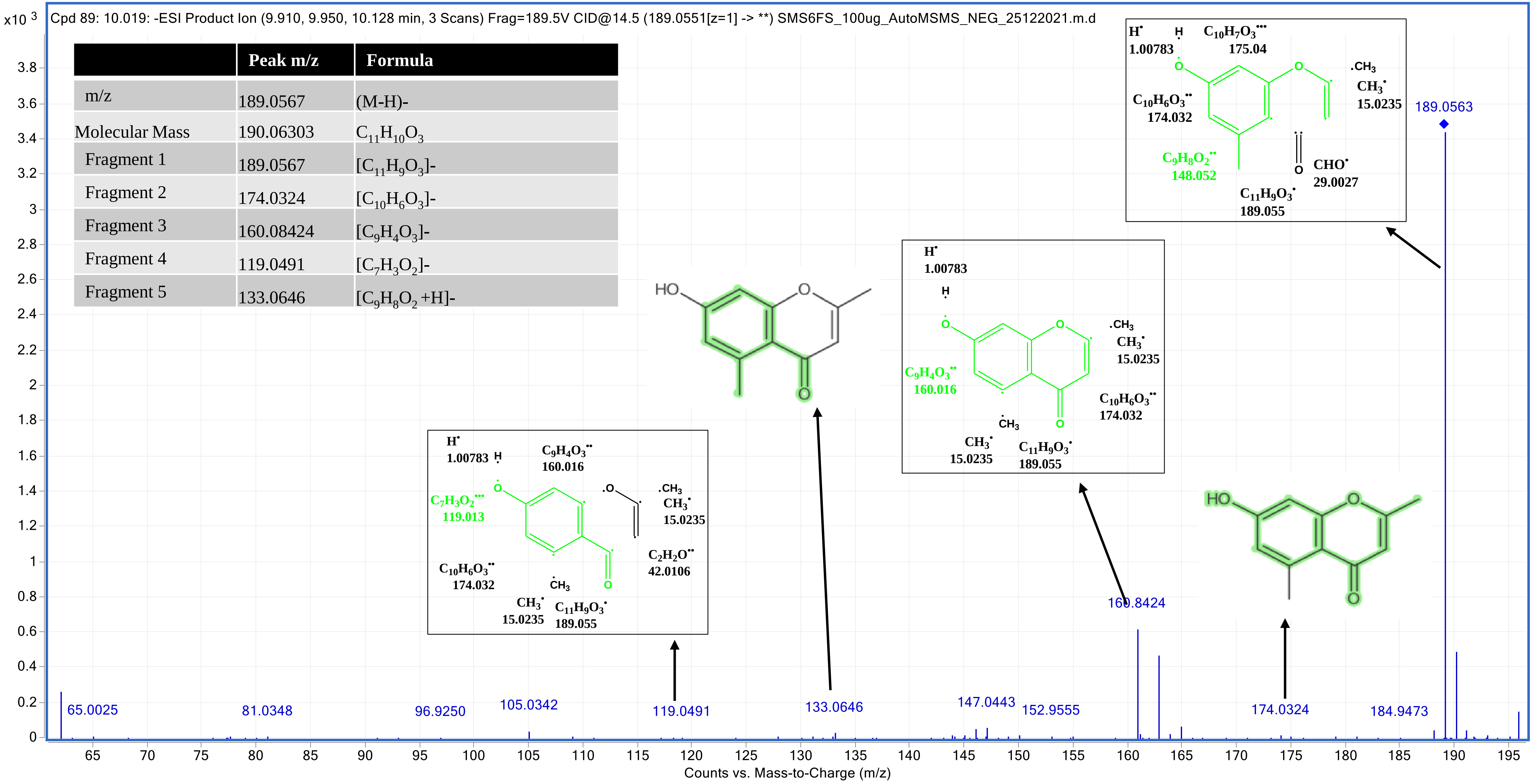

### Alternarian acid (6CFS) _10.752

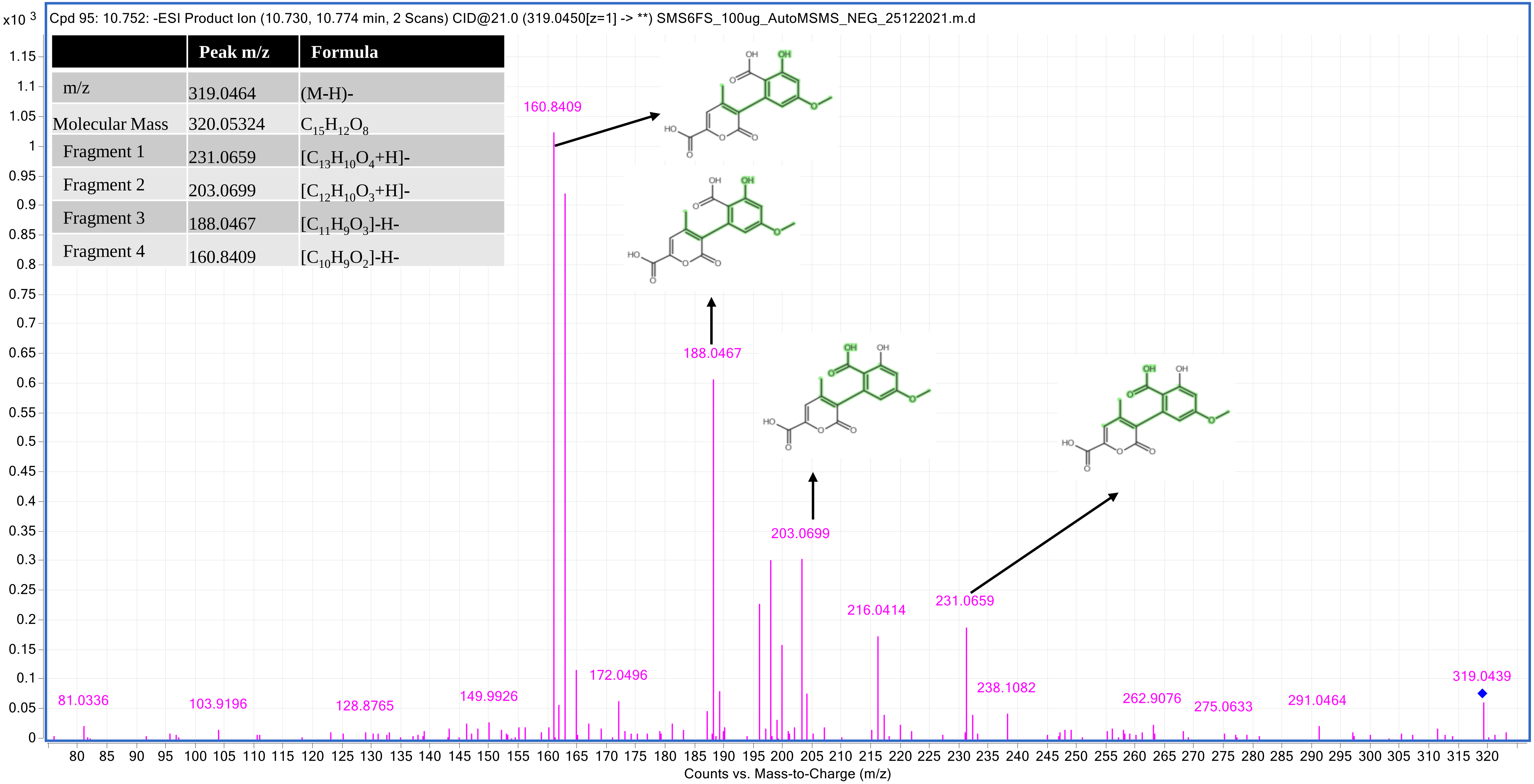


| | Peak m/z | Formula |
|---|---|---|
| m/z | 319.0464 | (M-H)- |
| Molecular Mass | 320.05324 | $C_{15}H_{12}O_8$ |
| Fragment 1 | 231.0659 | $[C_{13}H_{10}O_4+H]^-$ |
| Fragment 2 | 203.0699 | $[C_{12}H_{10}O_3+H]^-$ |
| Fragment 3 | 188.0467 | $[C_{11}H_9O_3]$-H- |
| Fragment 4 | 160.8409 | $[C_{10}H_9O_2]$-H- |

### 1,8-dihydroxynaphthalene (7CFS) _6.536

| | Peak m/z | Formula |
|---|---|---|
| m/z | 205.504 | (M+HCOO)- |
| Molecular Mass | 160.05246 | $C_{10}H_8O_2$ |
| Fragment 1 | 118.9931 | $[C_8H_6O]^-$ |
| Fragment 2 | 75.0032 | $[C_6H_3O]^-$ |
| Fragment 3 | 149.0256 | $[C_9H_8O_2]^-$ |
| Fragment 4 | 105.0338 | $[C_7H_5O]^-$ |

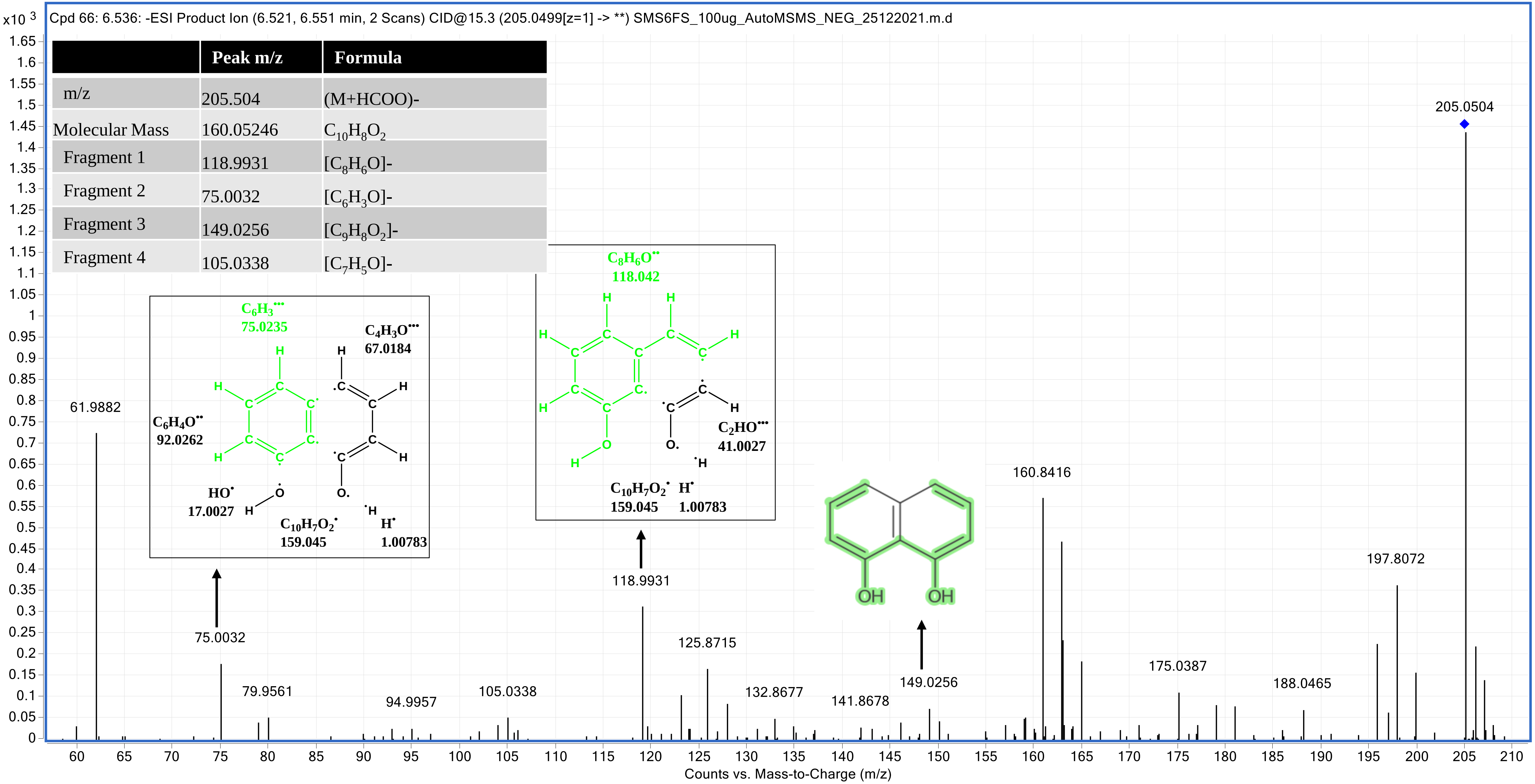

### Aspergone Q (8CFS) _11.941

| | Peak m/z | Formula |
|---|---|---|
| m/z | 223.0609 | (M-H)- [-$H_2O$] |
| Molecular Mass | 242.07908 | $C_{11}H_{14}O_6$ |
| Fragment 1 | 223.0606 | $[C_{11}H_{13}O_5]^-$ |
| Fragment 2 | 191.0347 | $[C_{10}H_{10}O_4$-2H]-H- |
| Fragment 3 | 162.0319 | $[C_9H_9O_3$-2H]-H- |
| Fragment 4 | 122.0383 | $[C_7H_9O_2$-2H]-H- |

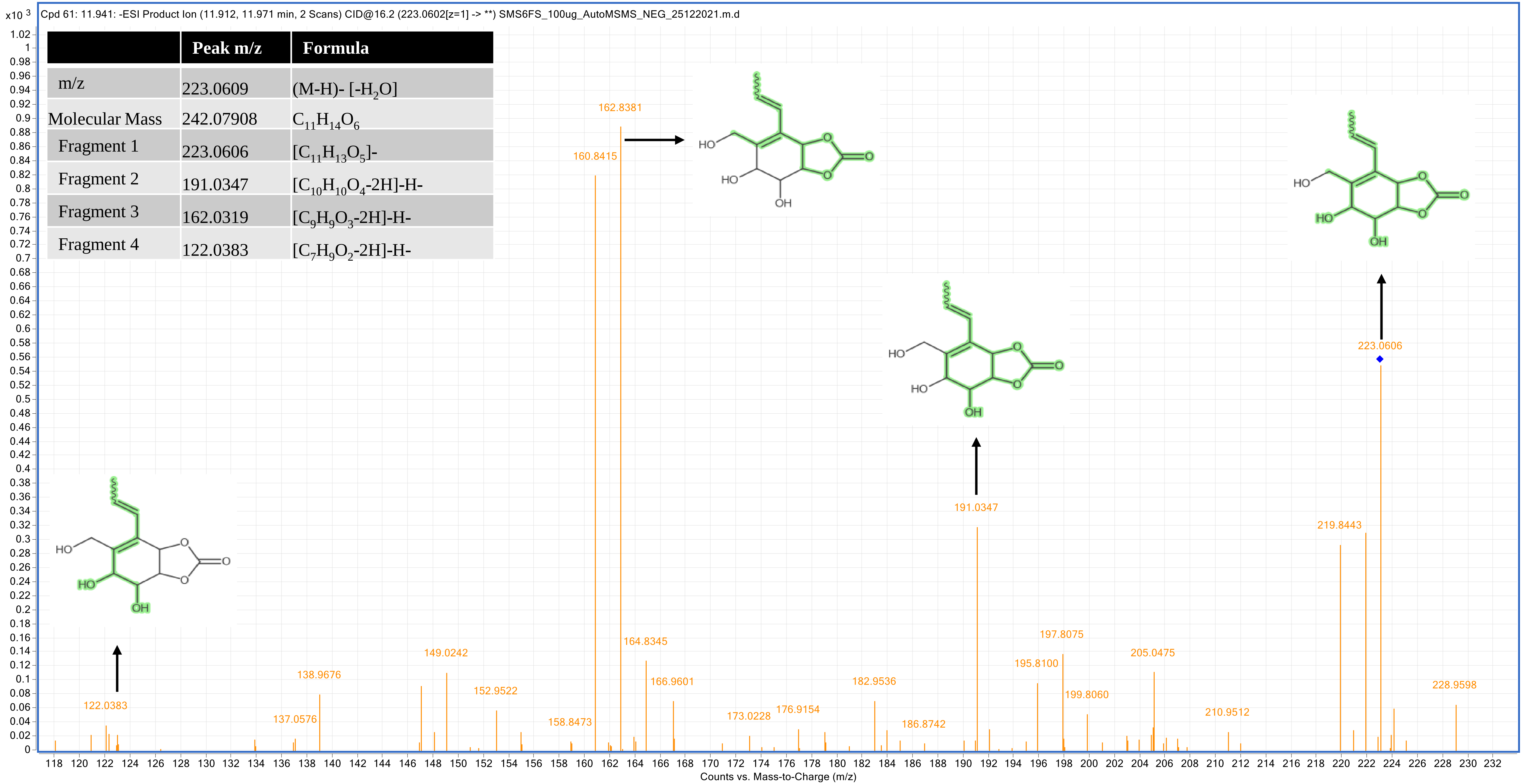

### 6-Epi-stemphytriol (9CFS) _12.968

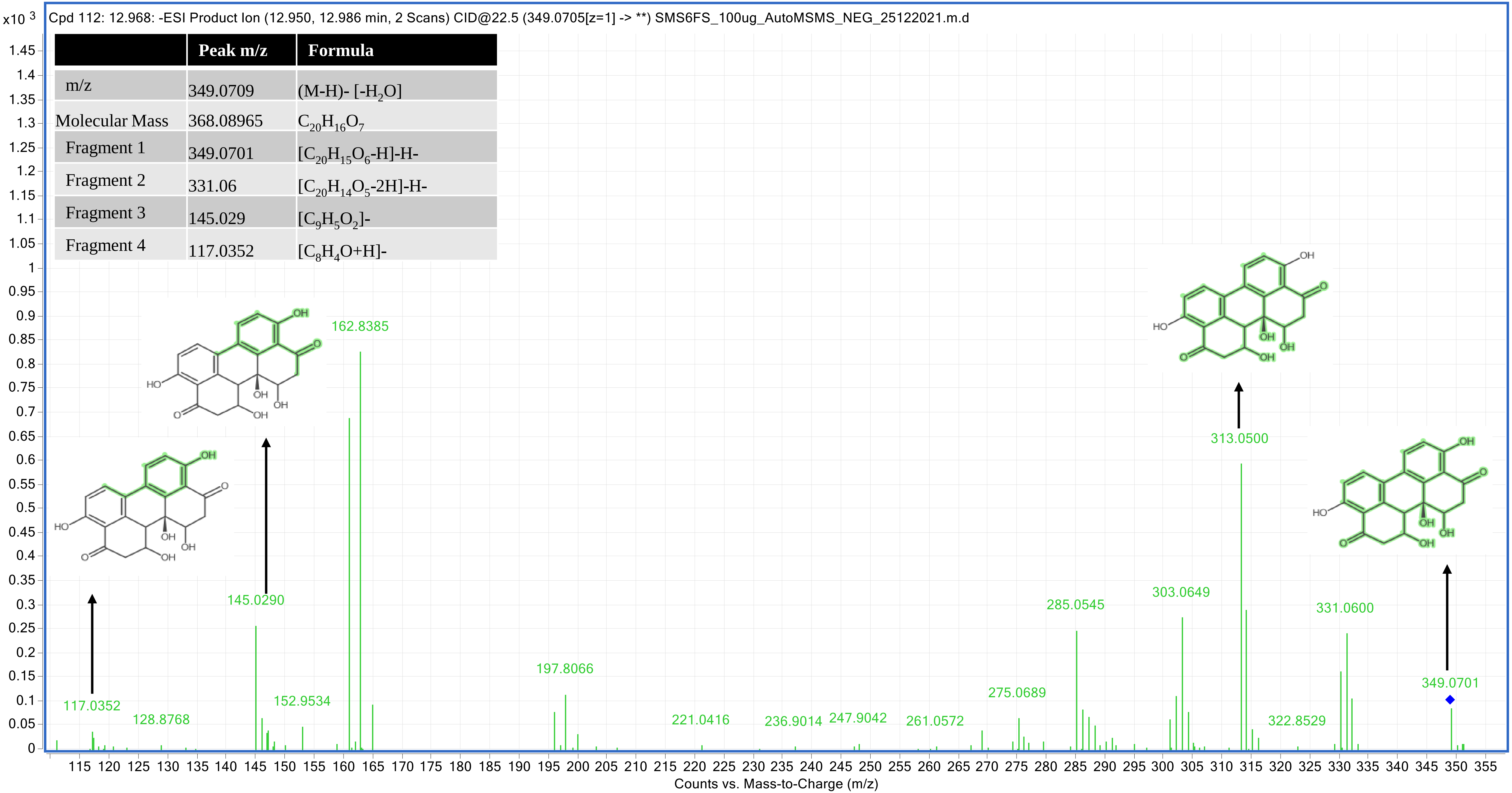


| | Peak m/z | Formula |
|---|---|---|
| m/z | 349.0709 | (M-H)- [-$H_2O$] |
| Molecular Mass | 368.08965 | $C_{20}H_{16}O_7$ |
| Fragment 1 | 349.0701 | [$C_{20}H_{15}O_6$-H]-H- |
| Fragment 2 | 331.06 | [$C_{20}H_{14}O_5$-2H]-H- |
| Fragment 3 | 145.029 | [$C_9H_5O_2$]- |
| Fragment 4 | 117.0352 | [$C_8H_4O$+H]- |

### 12-Methoxycitromycin (10CFS) _13.204

| | Peak m/z | Formula |
|---|---|---|
| m/z | 287.0568 | (M+HCOO)- |
| Molecular Mass | 260.06851 | $C_{14}H_{12}O_5$ |
| Fragment 1 | 229.05 | $[[C_{13}H_9O_4]$- |
| Fragment 2 | 228.0424 | $[C_{13}H_9O_4]$- |
| Fragment 3 | 200.0476 | $[C_{12}H_8O_4]$- |
| Fragment 4 | 211.0402 | $[C_{13}H_8O_3]$-H- |
| Fragment 5 | 243.06632 | $[C_{14}H_{11}O_4]$- |

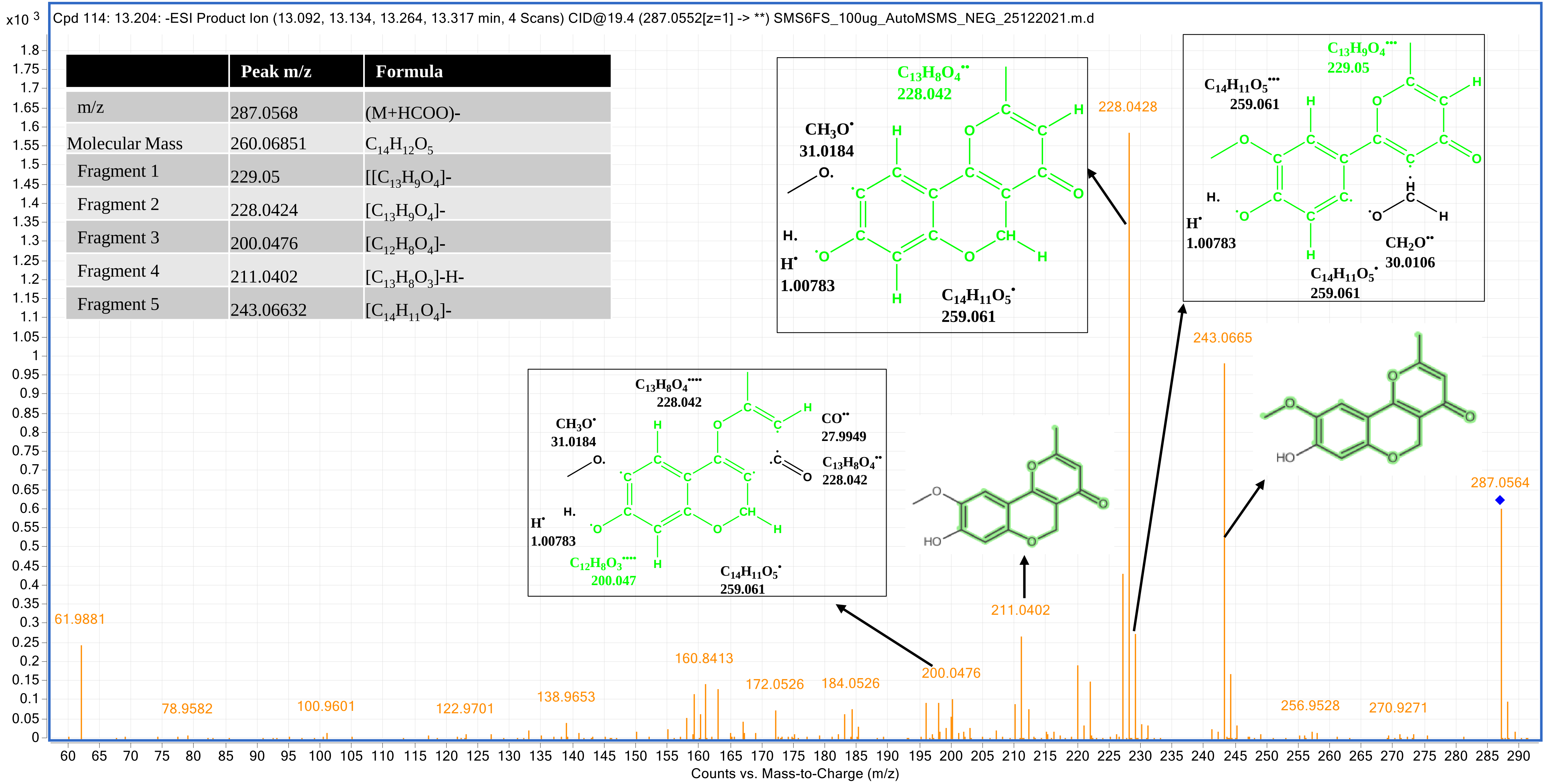

### 4-Hydroxyalternariol 9-methyl ether (11CFS) _13.532

| | Peak m/z | Formula |
|---|---|---|
| m/z | 287.0565 | (M-H)- |
| Molecular Mass | 288.06342 | $C_{15}H_{12}O_6$ |
| Fragment 1 | 271.0609 | $[C_{15}H_{11}O_5]$- |
| Fragment 2 | 245.0826 | $[C_{13}H_9O_5]$- |
| Fragment 3 | 230.0586 | $[C_{13}H_{10}O_4]$- |
| Fragment 4 | 160.0527 | $[C_{10}H_9O_2]$- |
| Fragment 5 | 202.0631 | $[C_{12}H_{11}O_3]$- |

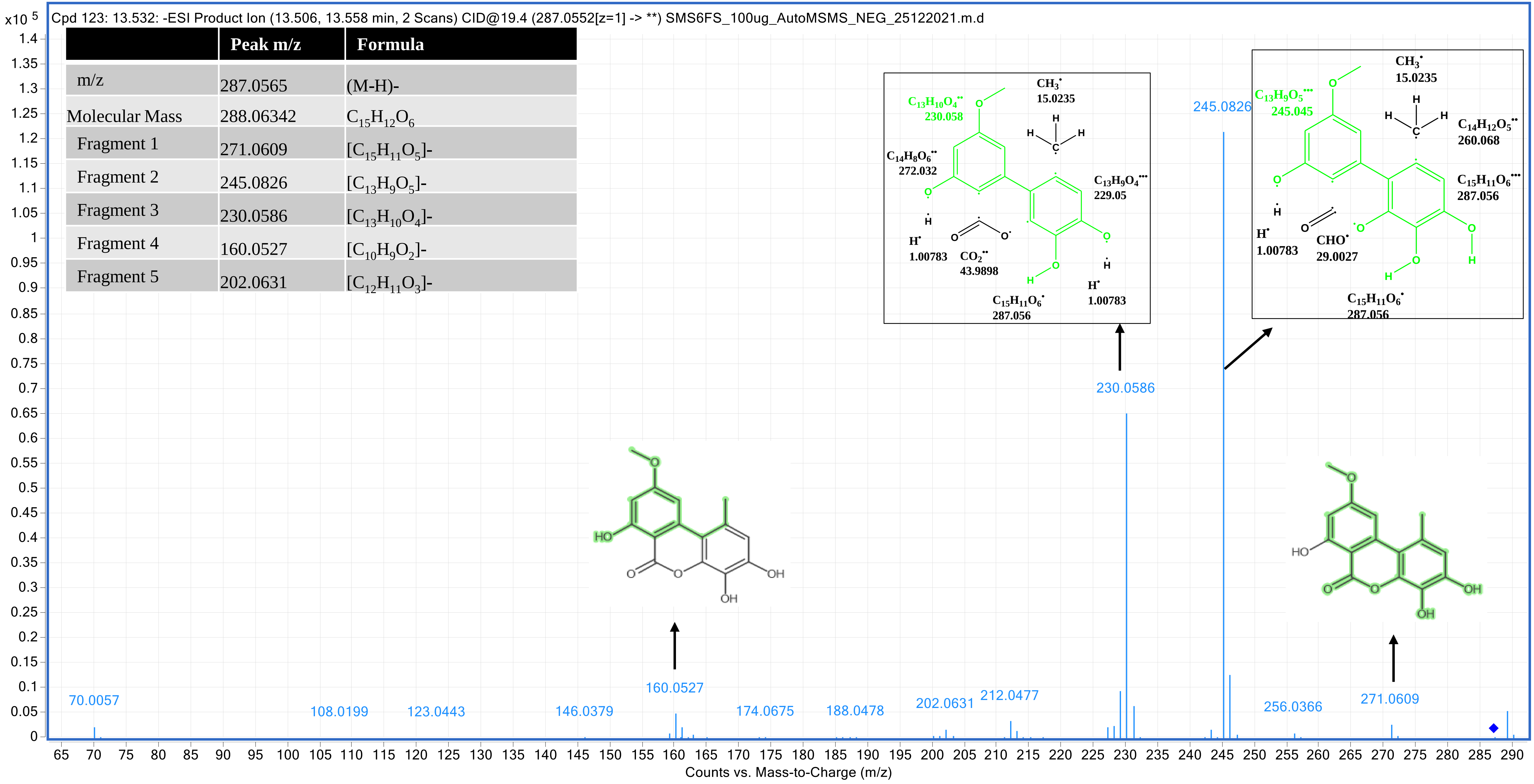

### Orthosporin (12CFS) _15.00

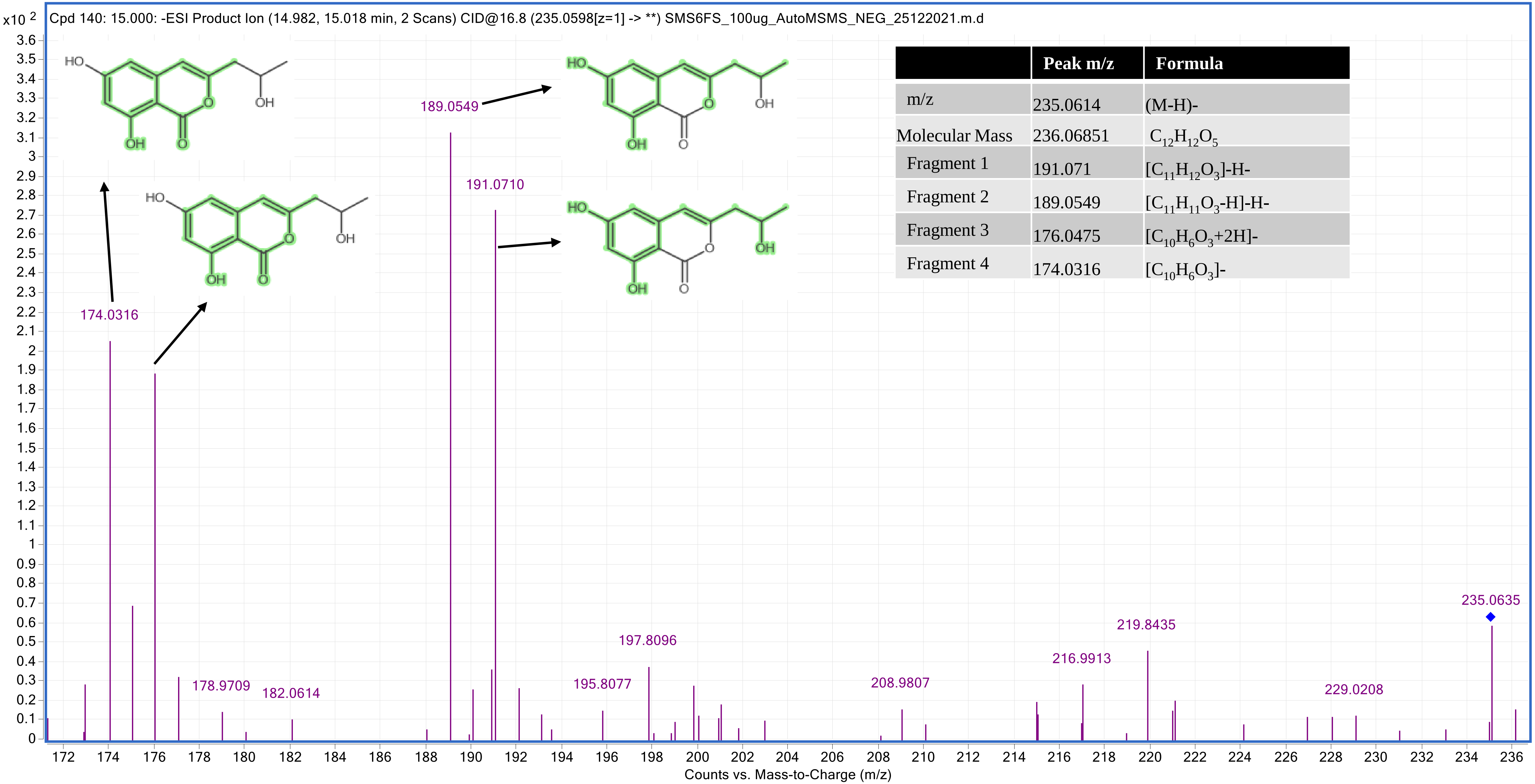


| | Peak m/z | Formula |
|---|---|---|
| m/z | 235.0614 | (M-H)- |
| Molecular Mass | 236.06851 | $C_{12}H_{12}O_5$ |
| Fragment 1 | 191.071 | $[C_{11}H_{12}O_3]$-H- |
| Fragment 2 | 189.0549 | $[C_{11}H_{11}O_3$-H]-H- |
| Fragment 3 | 176.0475 | $[C_{10}H_6O_3$+2H]- |
| Fragment 4 | 174.0316 | $[C_{10}H_6O_3]$- |

### Altenusin (13CFS) _19.829

| | Peak m/z | Formula |
|---|---|---|
| m/z | 317.0657 | (M+HCOO)- [-$H_2O$] |
| Molecular Mass | 290.07908 | $C_{15}H_{14}O_6$ |
| Fragment 1 | 272.068 | [$C_{15}H_{12}O_5$]- |
| Fragment 2 | 271.016 | [$C_{15}H_{11}O_5$]- |
| Fragment 3 | 270.0149 | [$C_{15}H_{10}O_5$]- |
| Fragment 4 | 242.0223 | [$C_{14}H_{10}O_4$]- |
| Fragment 5 | 198.0313 | [$C_{13}H_{10}O_2$]- |

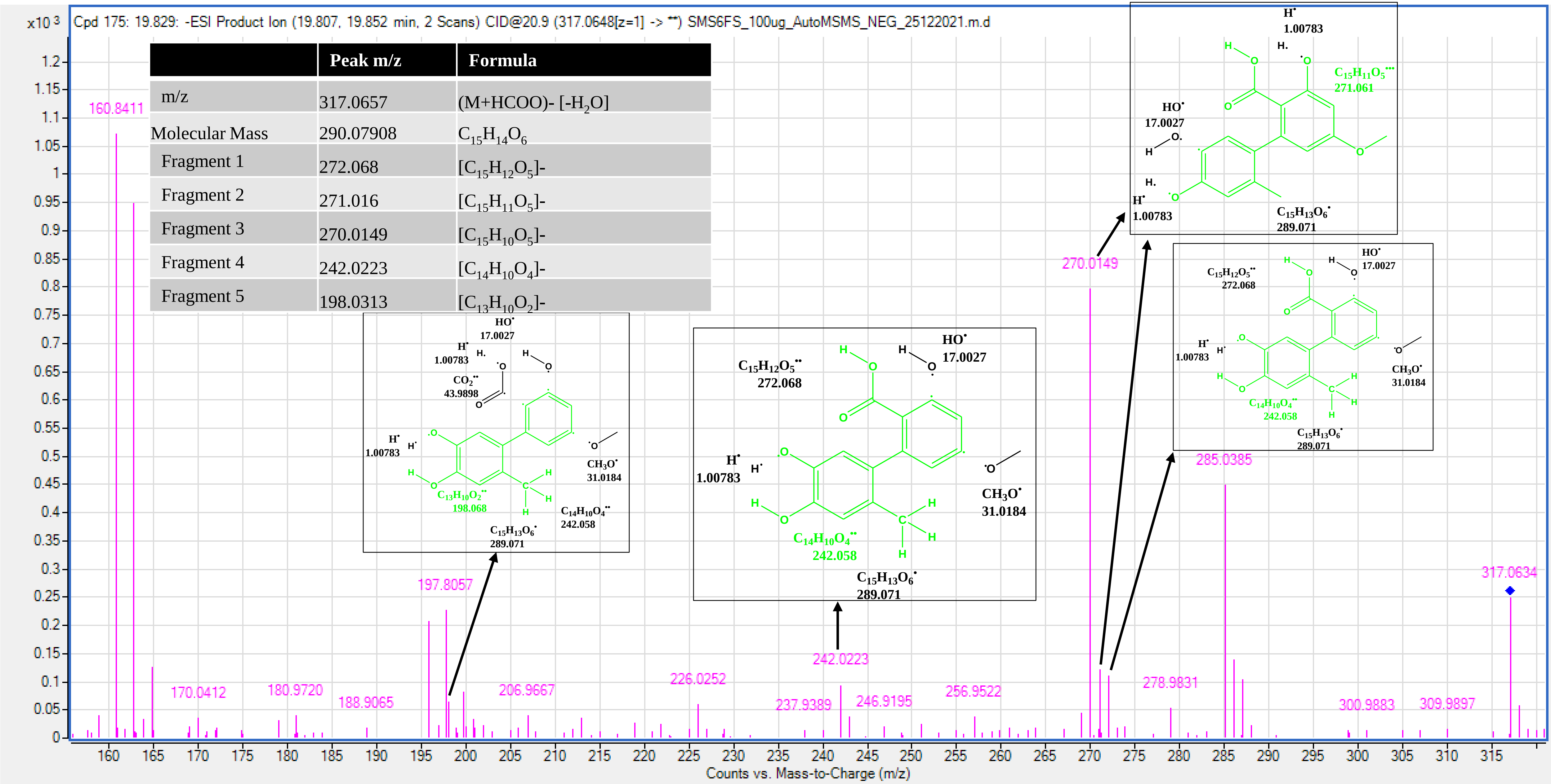

In the Unique Full-strength mode (UFS), MS/MS spectra of 6 molecules were generated

### 4-Ethylcatechol (1UFS) _5.595

| | Peak m/z | Formula |
|---|---|---|
| m/z | 119.0501 | (M-H)- [-$H_2O$] |
| Molecular Mass | 138.06812 | $C_8H_{10}O_2$ |
| Fragment 1 | 119.050 | [$C_8H_9O$-H]-H- |
| Fragment 2 | 93.0340 | [$C_6H_4$+H]- |
| Fragment 3 | 61.9881 | [$C_2H_4O_2$]- |

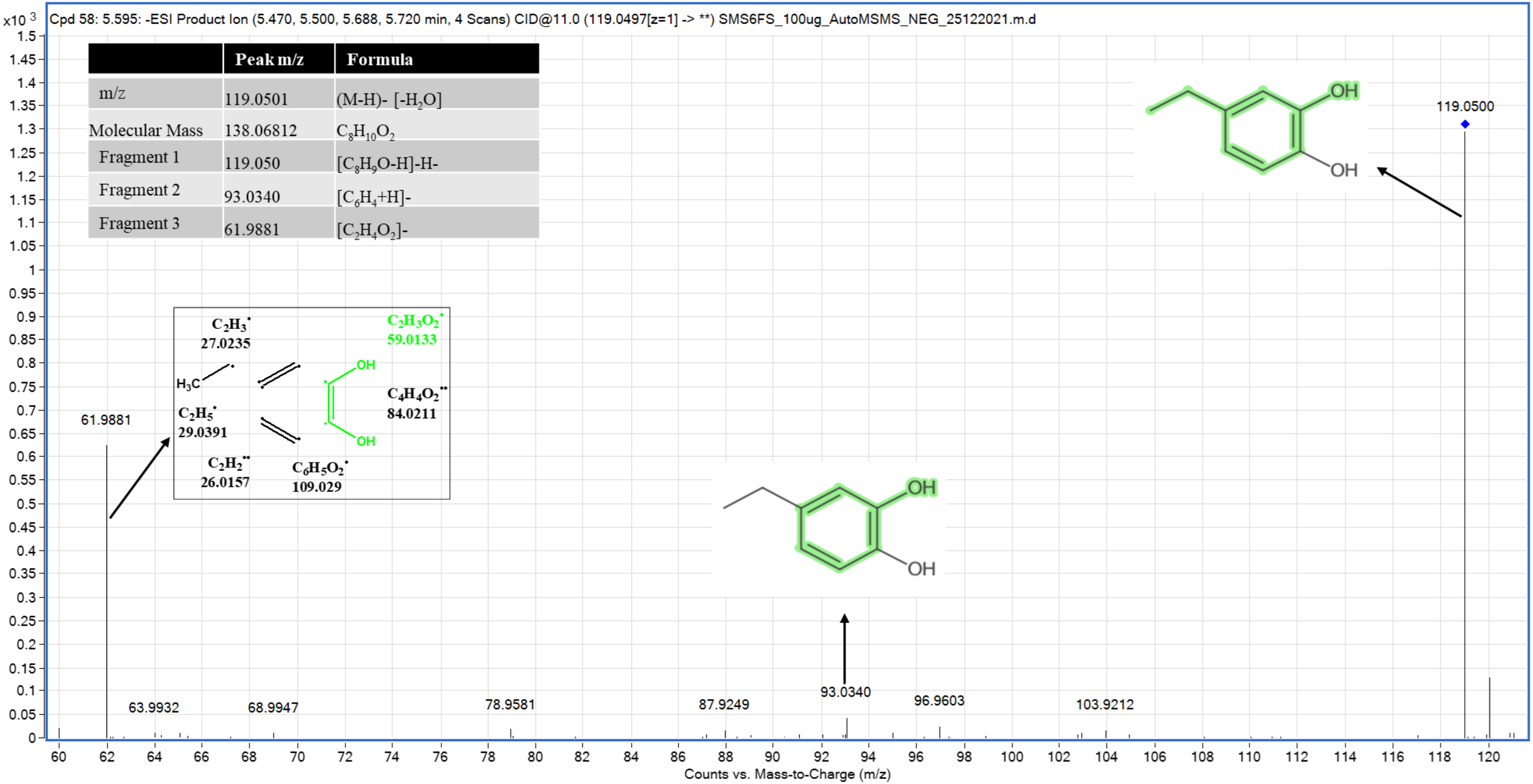

### p-Coumaric acid (2UFS) _7.023

| | Peak m/z | Formula |
|---|---|---|
| m/z | 209.0458 | (M+HCOO)- |
| Molecular Mass | 164.04737 | $C_9H_8O_3$ |
| Fragment 1 | 123.0453 | $[C_7H_7O_2]$- |
| Fragment 2 | 118.9625 | $[C_5H_6O]$- |

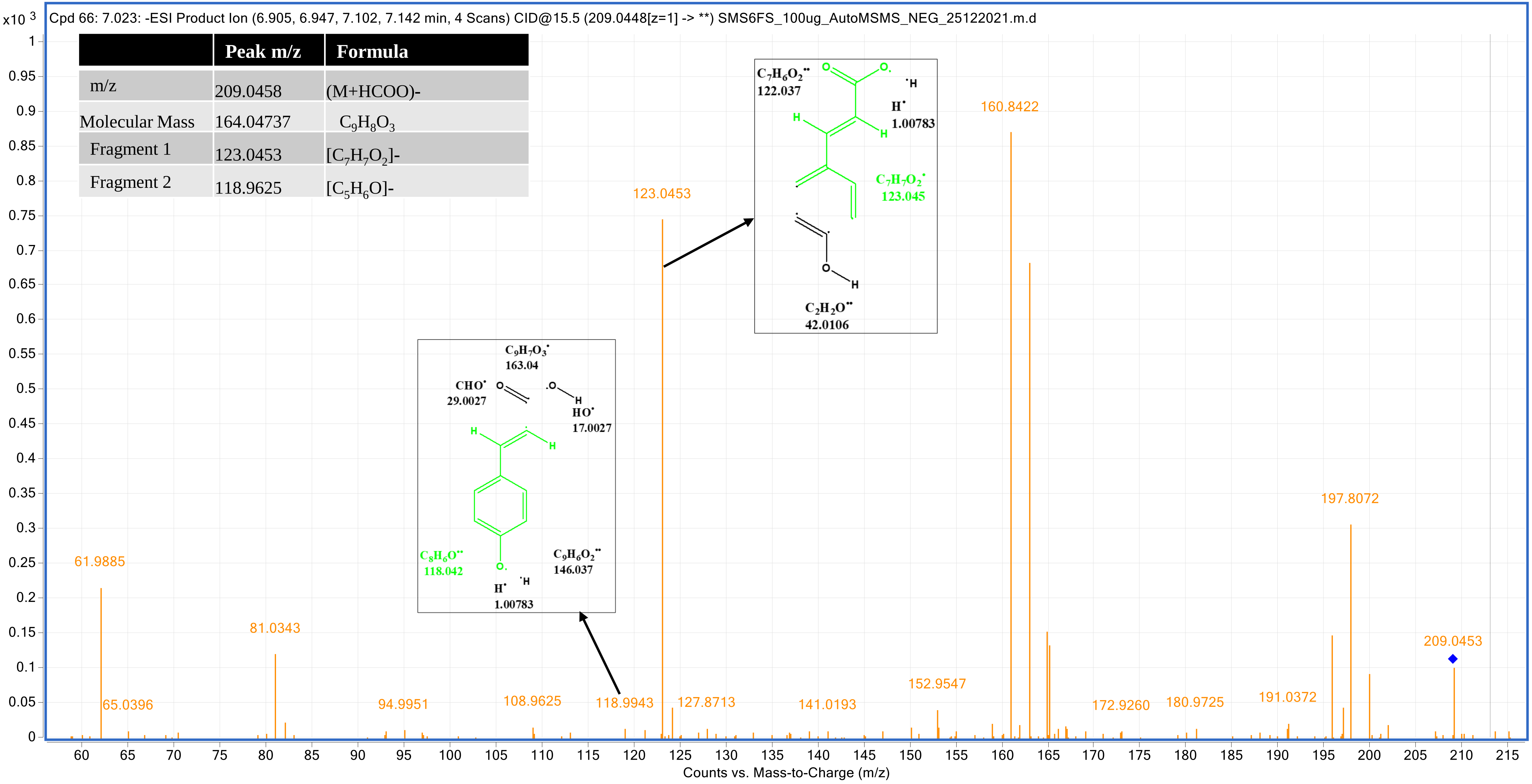

### Diaportinol (3UFS) _9.382

| | Peak m/z | Formula |
|---|---|---|
| m/z | 247.0613 | ((M-H)- [-$H_2O$] |
| Molecular Mass | 266.07908 | $C_{13}H_{14}O_6$ |
| Fragment 1 | 219.8431 | $[C_{12}H_{11}O_4]^-$ |
| Fragment 2 | 210.9479 | $[C_{10}H_{10}O_4]^-$ |
| Fragment 3 | 203.0703 | $[C_{12}H_{11}O_3]^-$ |
| Fragment 4 | 166.9587 | $[C_9H_{10}O_3]^-$ |

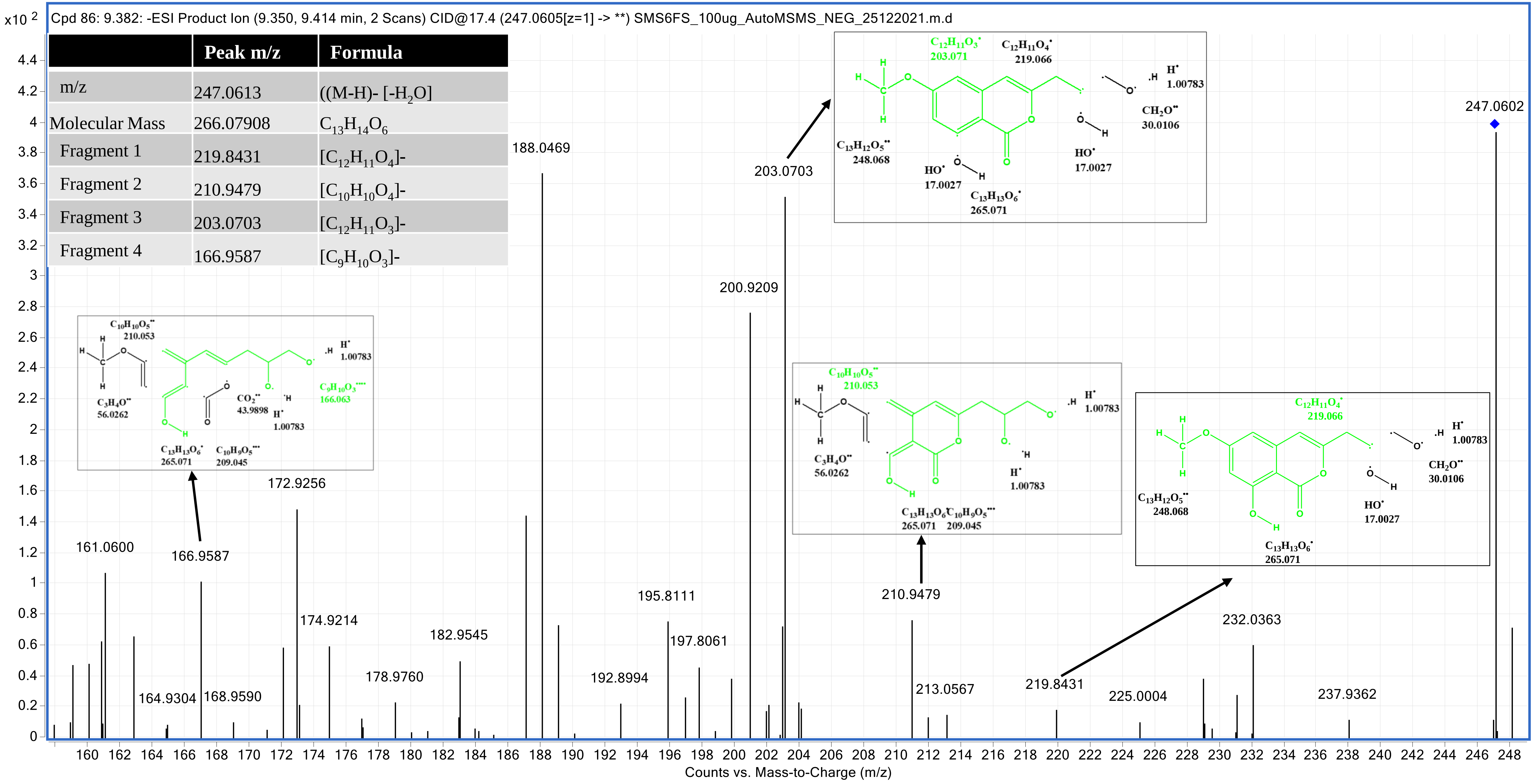

### Phenylacetic acid (4UFS) _5.707

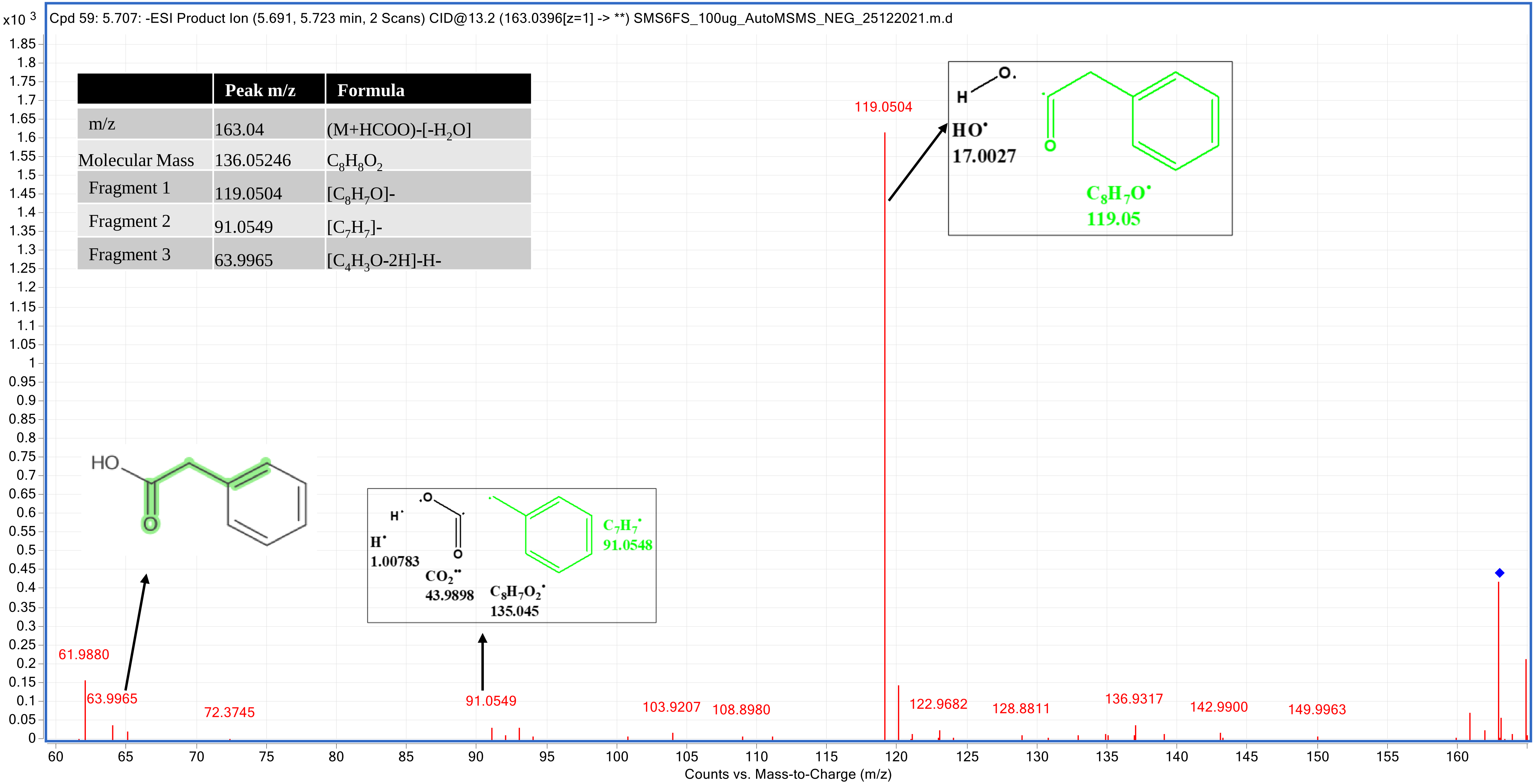


| | Peak m/z | Formula |
|---|---|---|
| m/z | 163.04 | (M+HCOO)-[-$H_2O$] |
| Molecular Mass | 136.05246 | $C_8H_8O_2$ |
| Fragment 1 | 119.0504 | [$C_8H_7O$]- |
| Fragment 2 | 91.0549 | [$C_7H_7$]- |
| Fragment 3 | 63.9965 | [$C_4H_3O$-2H]-H- |

### Procyanidin dimer B1 (5UFS) _16.753

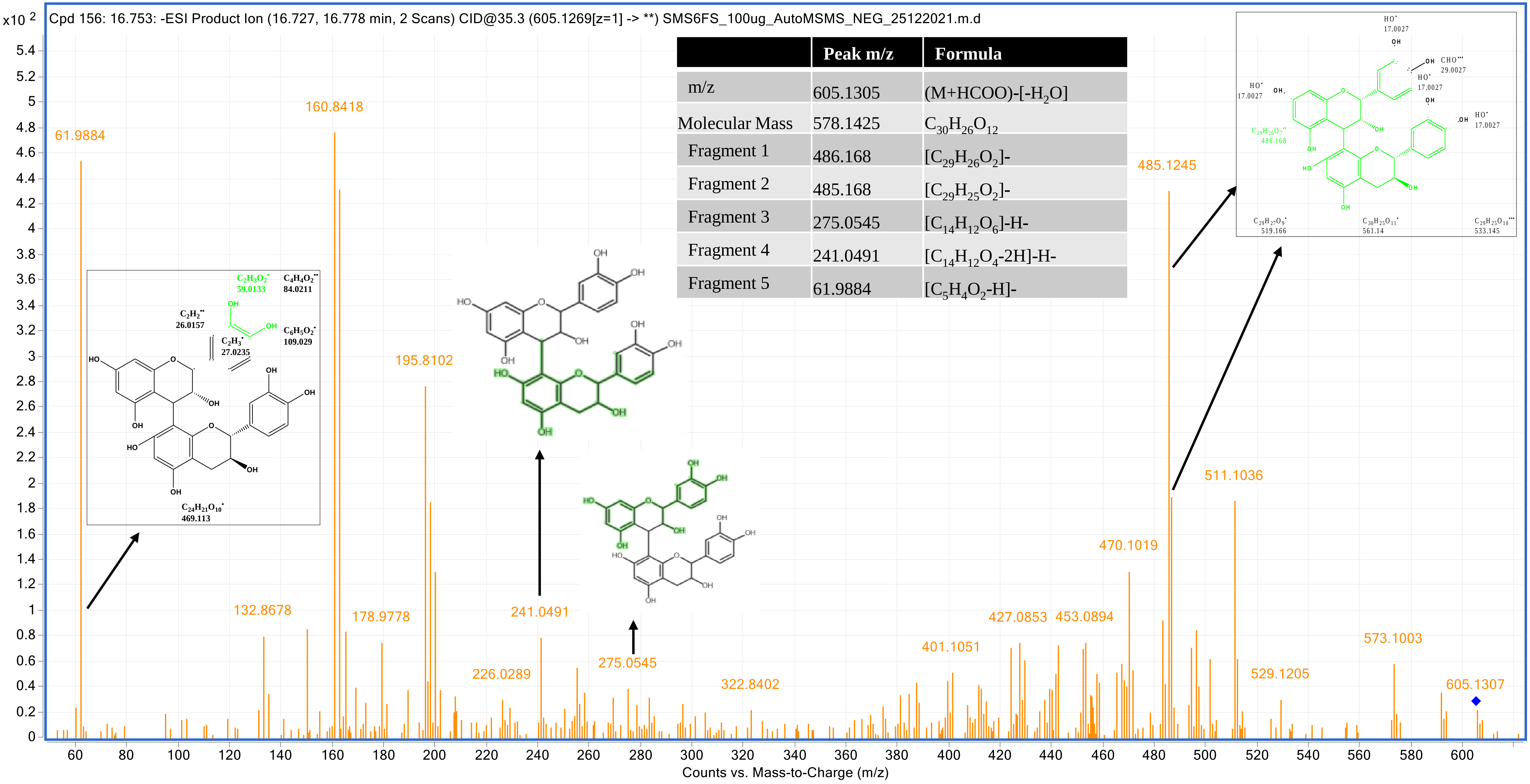


| | Peak m/z | Formula |
|---|---|---|
| m/z | 605.1305 | $(M+HCOO)-[-H_2O]$ |
| Molecular Mass | 578.1425 | $C_{30}H_{26}O_{12}$ |
| Fragment 1 | 486.168 | $[C_{29}H_{26}O_2]-$ |
| Fragment 2 | 485.168 | $[C_{29}H_{25}O_2]-$ |
| Fragment 3 | 275.0545 | $[C_{14}H_{12}O_6]-H-$ |
| Fragment 4 | 241.0491 | $[C_{14}H_{12}O_4-2H]-H-$ |
| Fragment 5 | 61.9884 | $[C_5H_4O_2-H]-$ |

### Theaflavin (6UFS) _22.497

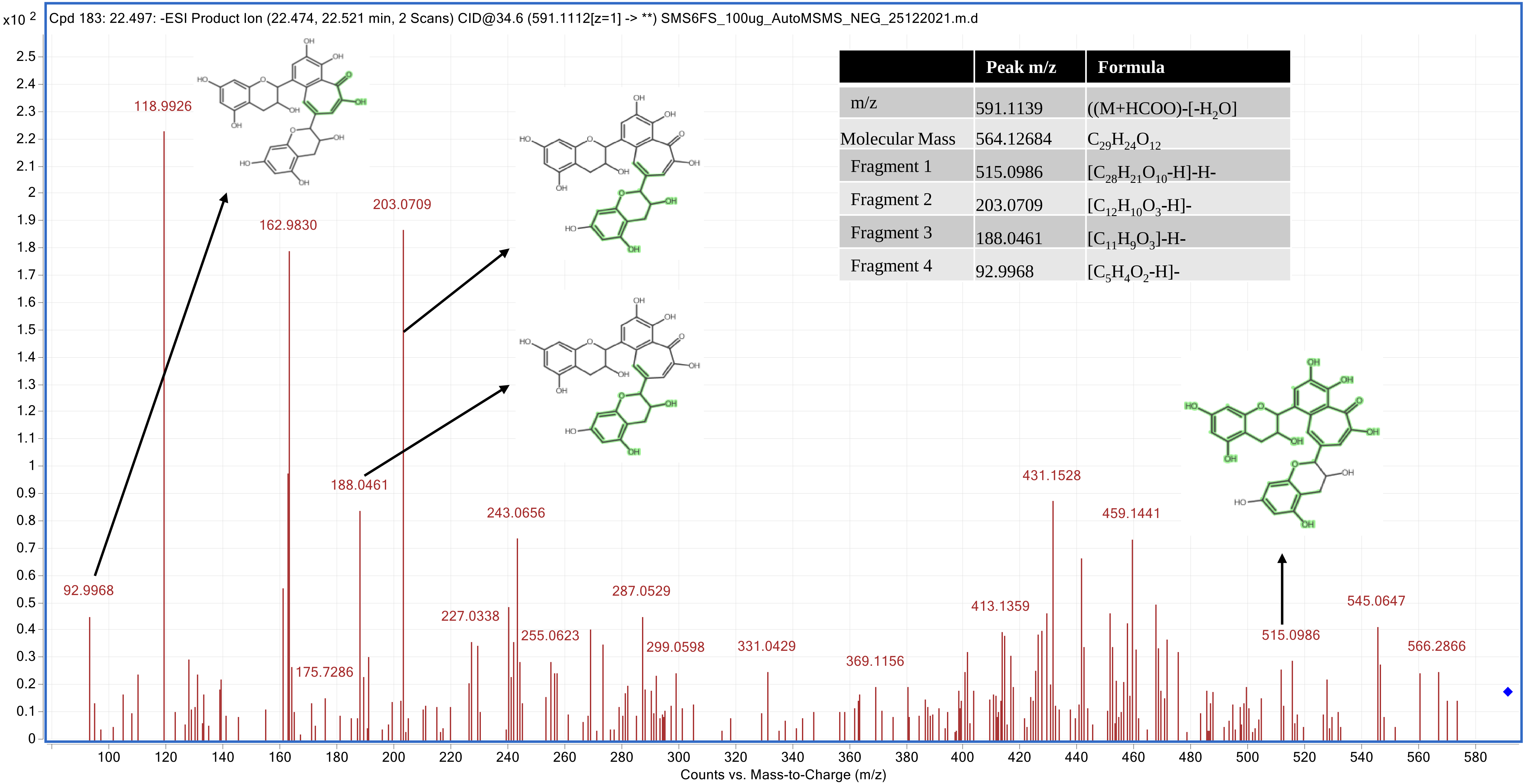


| | Peak m/z | Formula |
|---|---|---|
| m/z | 591.1139 | ((M+HCOO)-[-$H_2O$] |
| Molecular Mass | 564.12684 | $C_{29}H_{24}O_{12}$ |
| Fragment 1 | 515.0986 | [$C_{28}H_{21}O_{10}$-H]-H- |
| Fragment 2 | 203.0709 | [$C_{12}H_{10}O_3$-H]- |
| Fragment 3 | 188.0461 | [$C_{11}H_9O_3$]-H- |
| Fragment 4 | 92.9968 | [$C_5H_4O_2$-H]- |

In the Unique quarter-strength mode (UQS), MS/MS spectra of 7 molecules were generated

### 6-O-desmethylterphenyllin (1UQS) _11.712

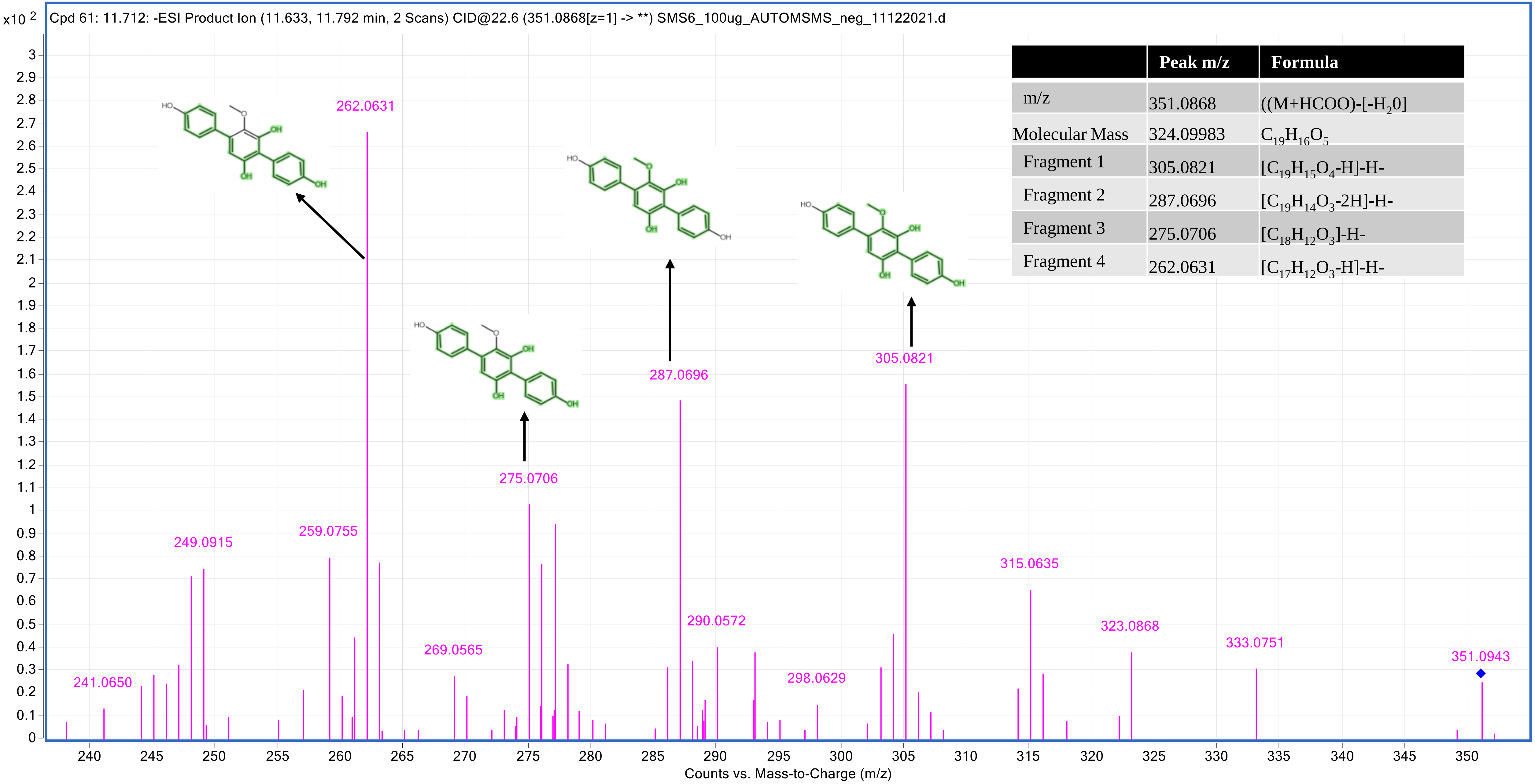


| | Peak m/z | Formula |
|---|---|---|
| m/z | 351.0868 | ((M+HCOO)-[-$H_2$0] |
| Molecular Mass | 324.09983 | $C_{19}H_{16}O_5$ |
| Fragment 1 | 305.0821 | [$C_{19}H_{15}O_4$-H]-H- |
| Fragment 2 | 287.0696 | [$C_{19}H_{14}O_3$-2H]-H- |
| Fragment 3 | 275.0706 | [$C_{18}H_{12}O_3$]-H- |
| Fragment 4 | 262.0631 | [$C_{17}H_{12}O_3$-H]-H- |

### Altertoxin I (2UQS) _11.712

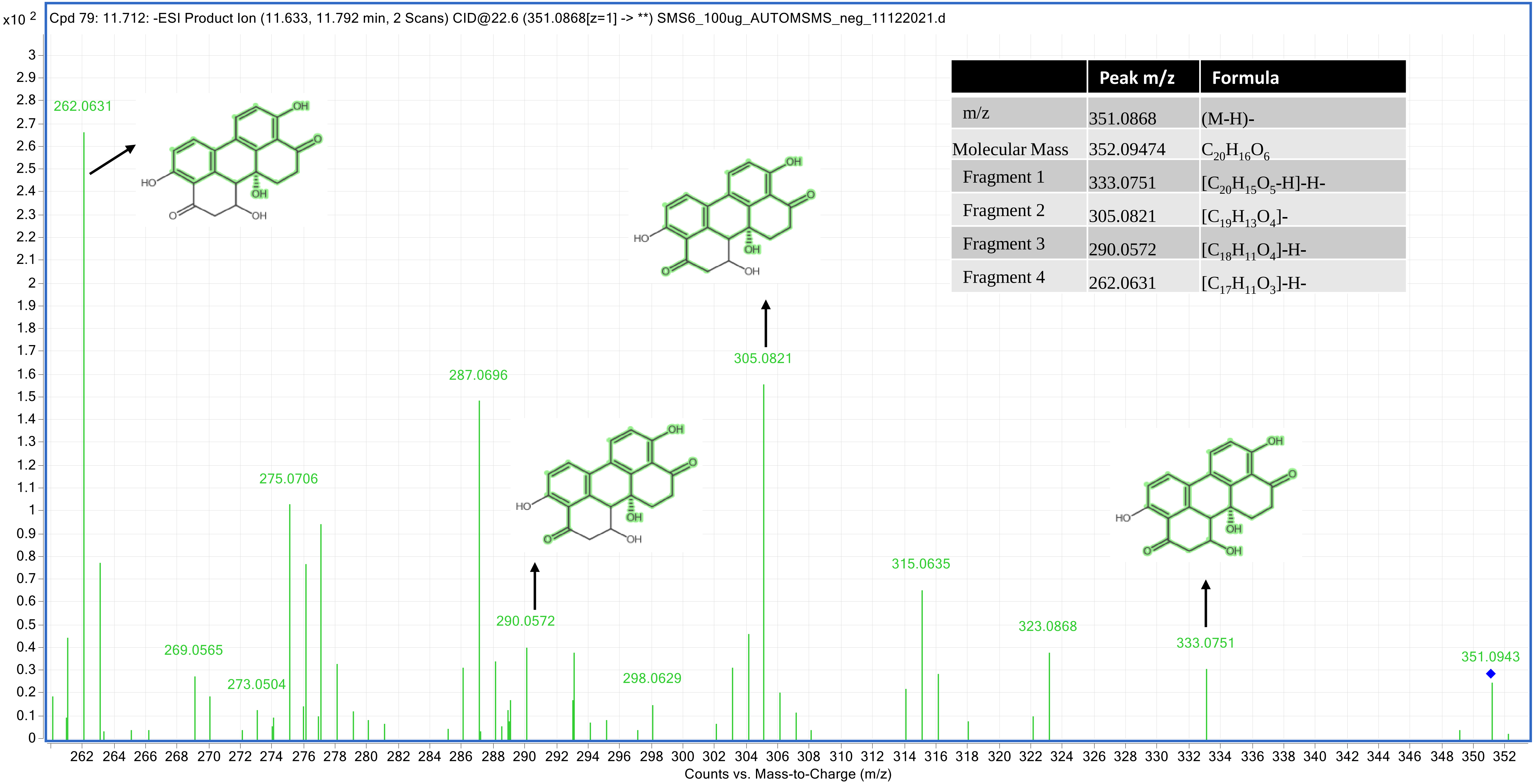


| | Peak m/z | Formula |
|---|---|---|
| m/z | 351.0868 | (M-H)- |
| Molecular Mass | 352.09474 | $C_{20}H_{16}O_6$ |
| Fragment 1 | 333.0751 | $[C_{20}H_{15}O_5$-H]-H- |
| Fragment 2 | 305.0821 | $[C_{19}H_{13}O_4]$- |
| Fragment 3 | 290.0572 | $[C_{18}H_{11}O_4]$-H- |
| Fragment 4 | 262.0631 | $[C_{17}H_{11}O_3]$-H- |

### Altechromone B (3UQS) _18.764

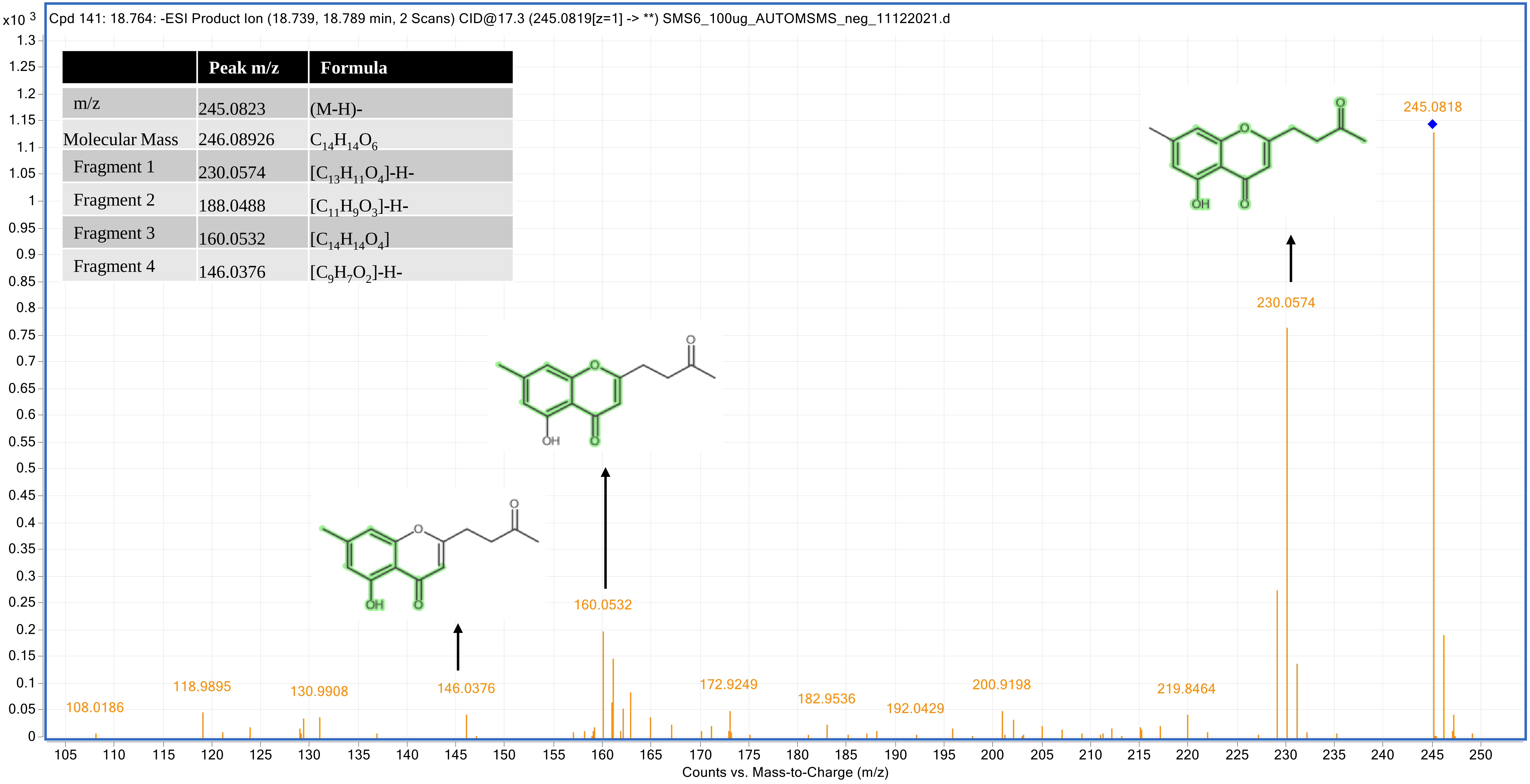


| | Peak m/z | Formula |
|---|---|---|
| m/z | 245.0823 | (M-H)- |
| Molecular Mass | 246.08926 | $C_{14}H_{14}O_6$ |
| Fragment 1 | 230.0574 | $[C_{13}H_{11}O_4]$-H- |
| Fragment 2 | 188.0488 | $[C_{11}H_9O_3]$-H- |
| Fragment 3 | 160.0532 | $[C_{14}H_{14}O_4]$ |
| Fragment 4 | 146.0376 | $[C_9H_7O_2]$-H- |

### Botryorhodine F (4UQS) _20.017

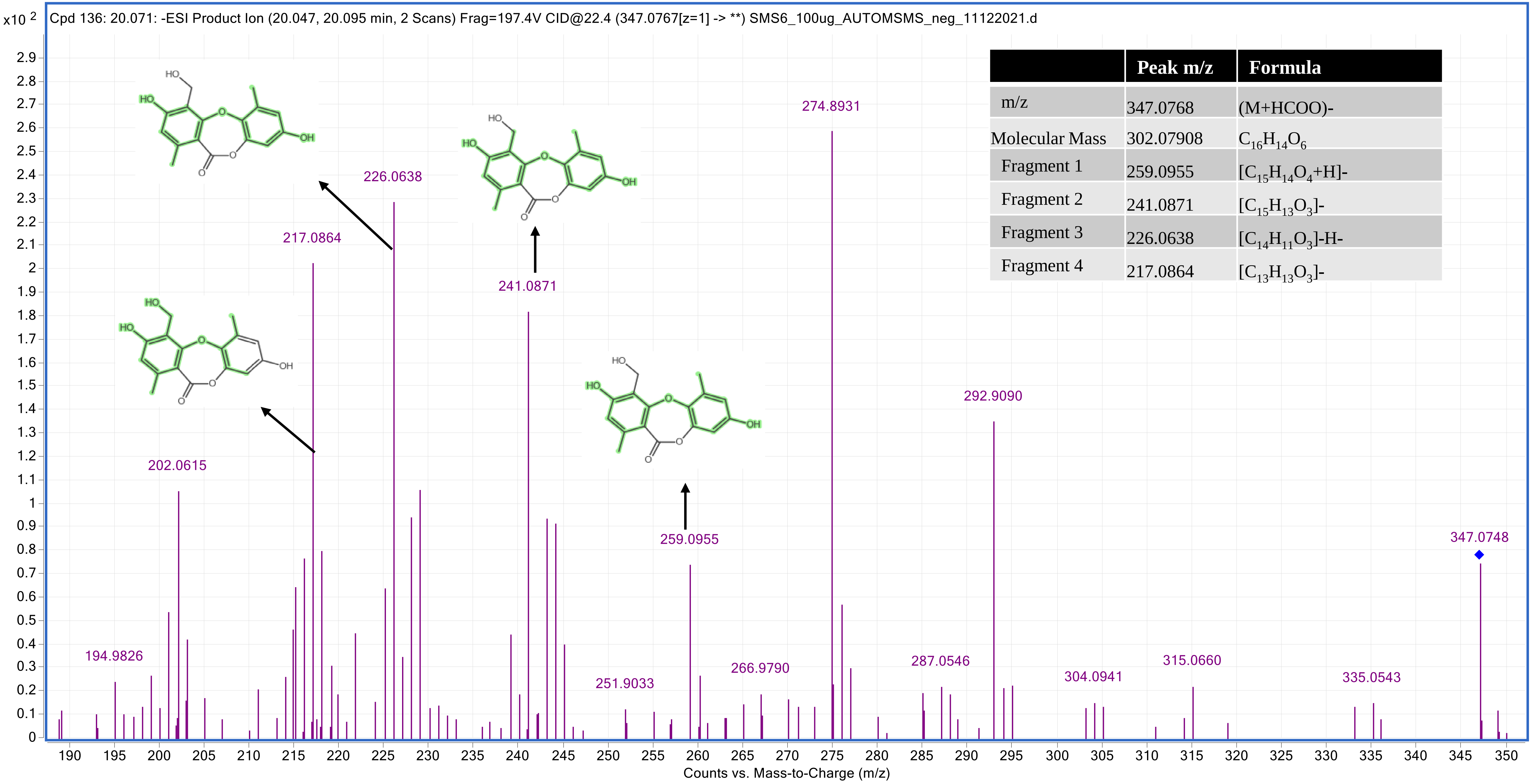


| | Peak m/z | Formula |
|---|---|---|
| m/z | 347.0768 | (M+HCOO)- |
| Molecular Mass | 302.07908 | $C_{16}H_{14}O_6$ |
| Fragment 1 | 259.0955 | $[C_{15}H_{14}O_4+H]^-$ |
| Fragment 2 | 241.0871 | $[C_{15}H_{13}O_3]^-$ |
| Fragment 3 | 226.0638 | $[C_{14}H_{11}O_3]$-H- |
| Fragment 4 | 217.0864 | $[C_{13}H_{13}O_3]^-$ |

### 3',4',7-Trihydroxyisoflavanone (5UQS) _22.848

| | Peak m/z | Formula |
|---|---|---|
| m/z | 317.0666 | (M+HCOO)- |
| Molecular Mass | 272.06851 | $C_{15}H_{12}O_5$ |
| Fragment 1 | 271.053 | $[C_{15}H_{11}O_5]^-$ |
| Fragment 2 | 270.0169 | $[C_{15}H_{10}O_5]^-$ |
| Fragment 3 | 242.0206 | $[C_{14}H_{10}O_4]^-$ |
| Fragment 4 | 226.0275 | $[C_{14}H_{10}O_3]^-$ |
| Fragment 5 | 198.0313 | $[C_{12}H_8O_3\text{-}H]^-$ |
| Fragment 6 | 214.0267 | $[C_{12}H_7O_4\text{-}H]^-$ |
| Fragment 7 | 256.0358 | $[C_{14}H_{10}O_5\text{-}H]\text{-}H^-$ |

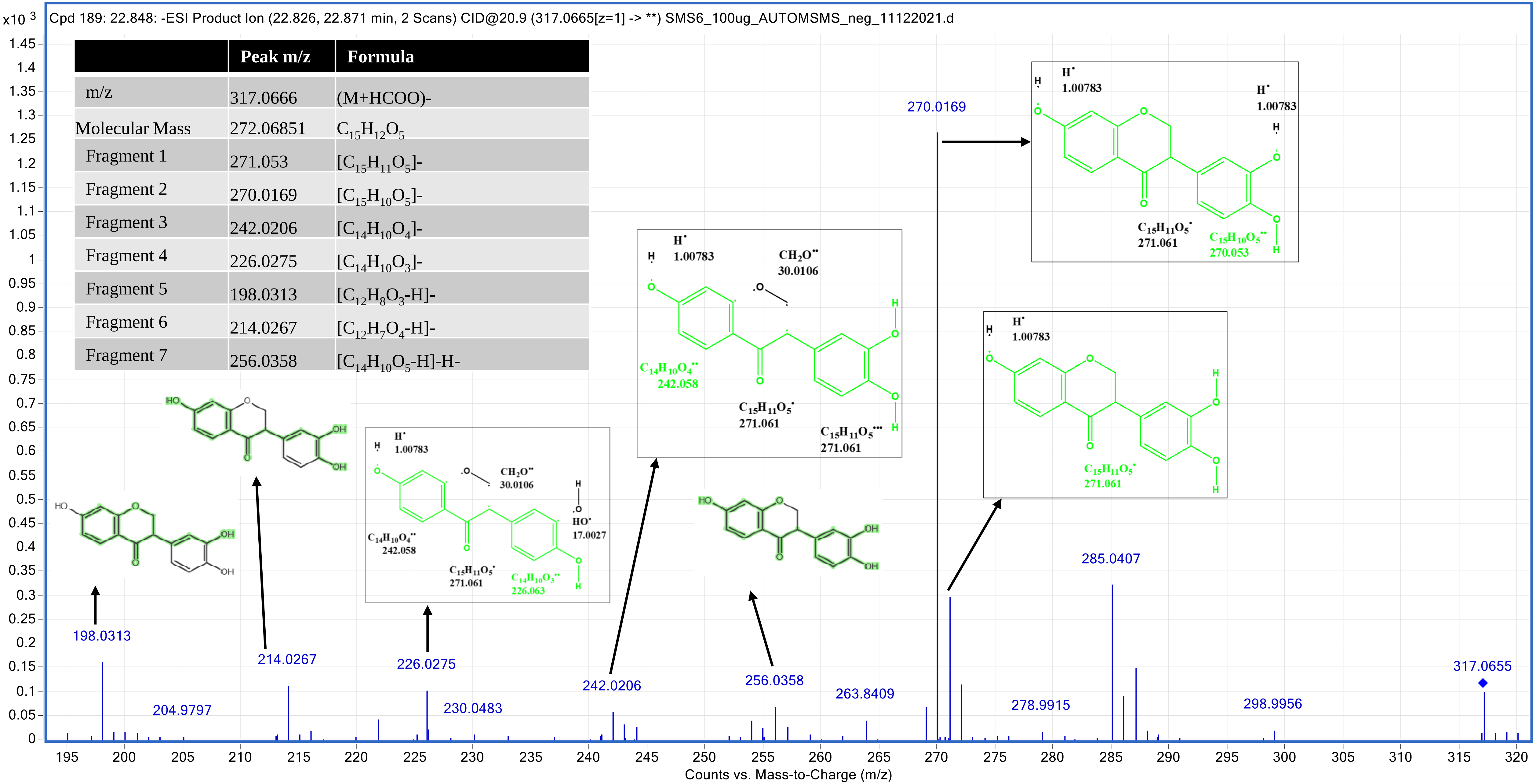

### Alternariol 9-methyl ether (6UQS) _25.377

| | Peak m/z | Formula |
|---|---|---|
| m/z | 317.0669 | (M+HCOO)- |
| Molecular Mass | 272.06851 | $C_{15}H_{12}O_5$ |
| Fragment 1 | 271.061 | $[C_{15}H_{11}O_5]^-$ |
| Fragment 2 | 270.061 | $[C_{15}H_{10}O_5]^-$ |
| Fragment 3 | 198.0338 | $[C_{13}H_{10}O_2]^-$ |
| Fragment 4 | 242.0206 | $[C_{13}H_6O_5]^-$ |

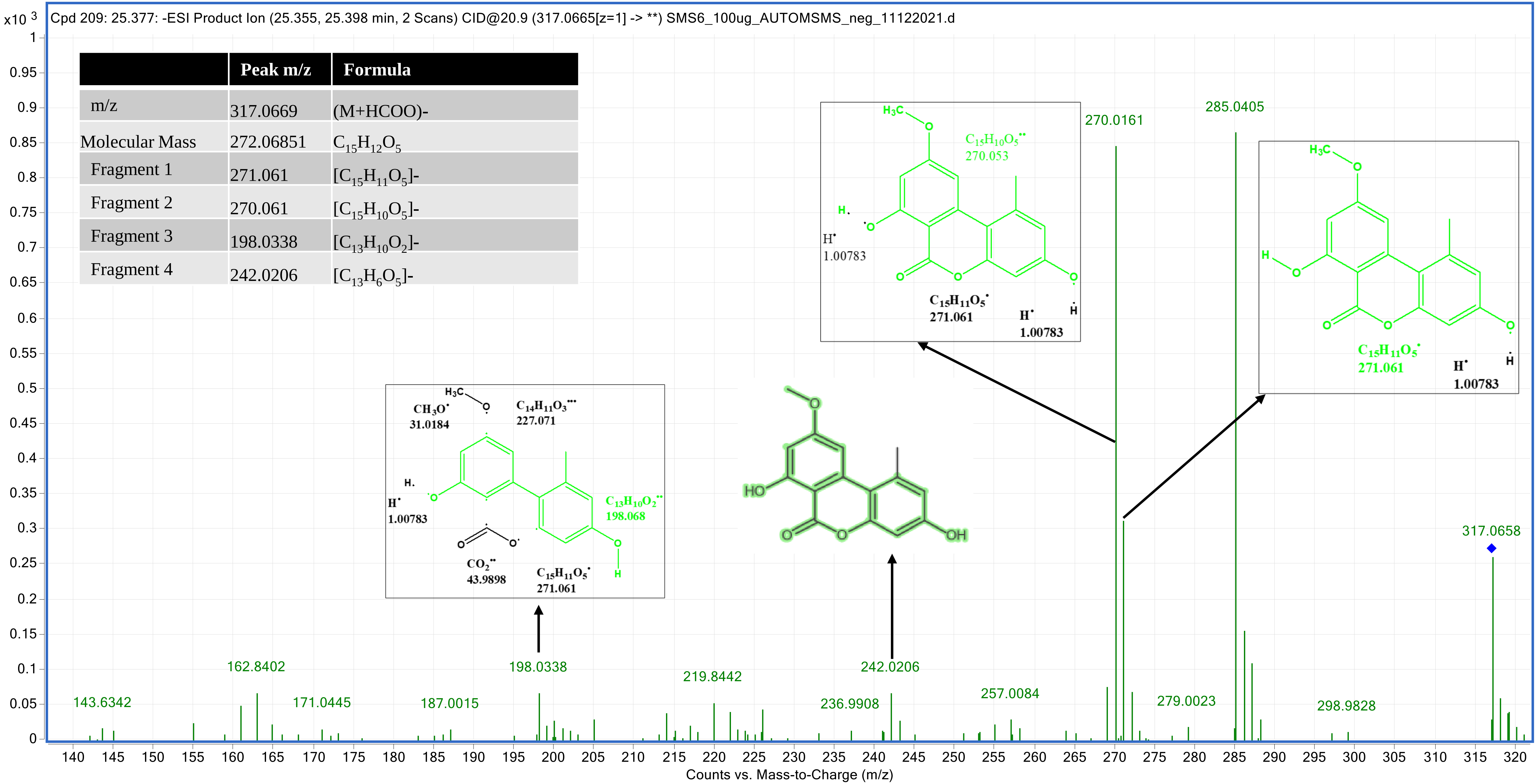

### Morin (7UQS) 26.284

|  | Peak m/z | Formula |
|---|---|---|
| m/z | 301.0349 | (M-H)- |
| Molecular Mass | 302.04267 | $C_{15}H_{10}O_7$ |
| Fragment 1 | 301.0349 | $[C_{14}H_9O_7]$- |
| Fragment 2 | 273.0399 | $[C_{14}H_9O_5]$- |
| Fragment 3 | 258.0159 | $[C_{13}H_8O_6$-H]-H- |
| Fragment 4 | 230.0240 | $[C_{12}H_7O_5]$-H- |

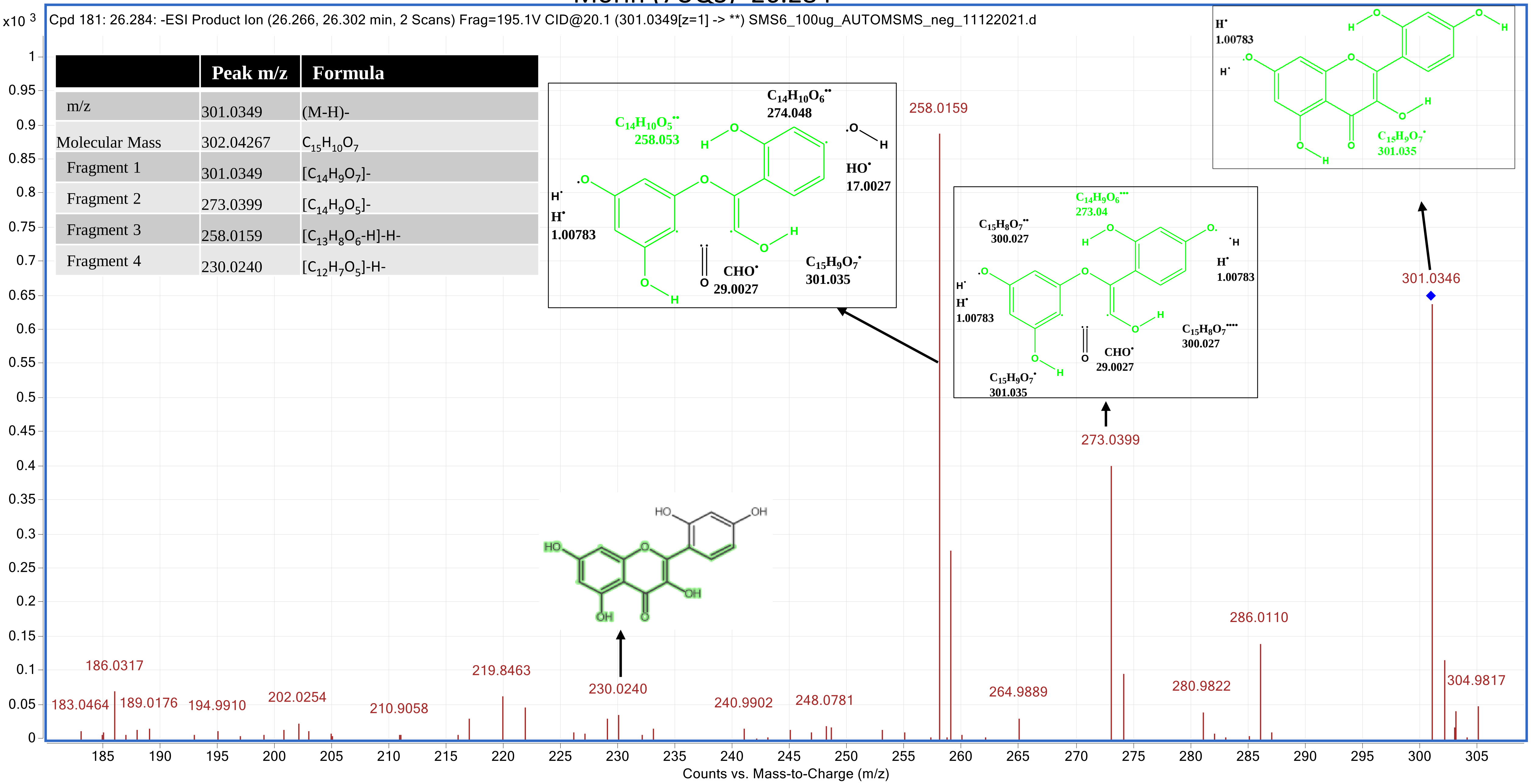

- In the Common quarter-strength mode (CQS), MS/MS spectra of 13 molecules were generated

### 4-Hydroxymellein(1CQS) _18.157

| | Peak m/z | Formula |
|---|---|---|
| m/z | 221.0458 | (M+HCOO)- [-$H_2O$] |
| Molecular Mass | 194.05794 | $C_{10}H_{10}O_4$ |
| Fragment 1 | 178.027 | $(C_9H_6O_4]^{-.}$ |
| Fragment 2 | 177.0553 | $[C_9H_5O_4]^-$ |
| Fragment 3 | 162.034 | $[C_9H_6O_3]^-$ |

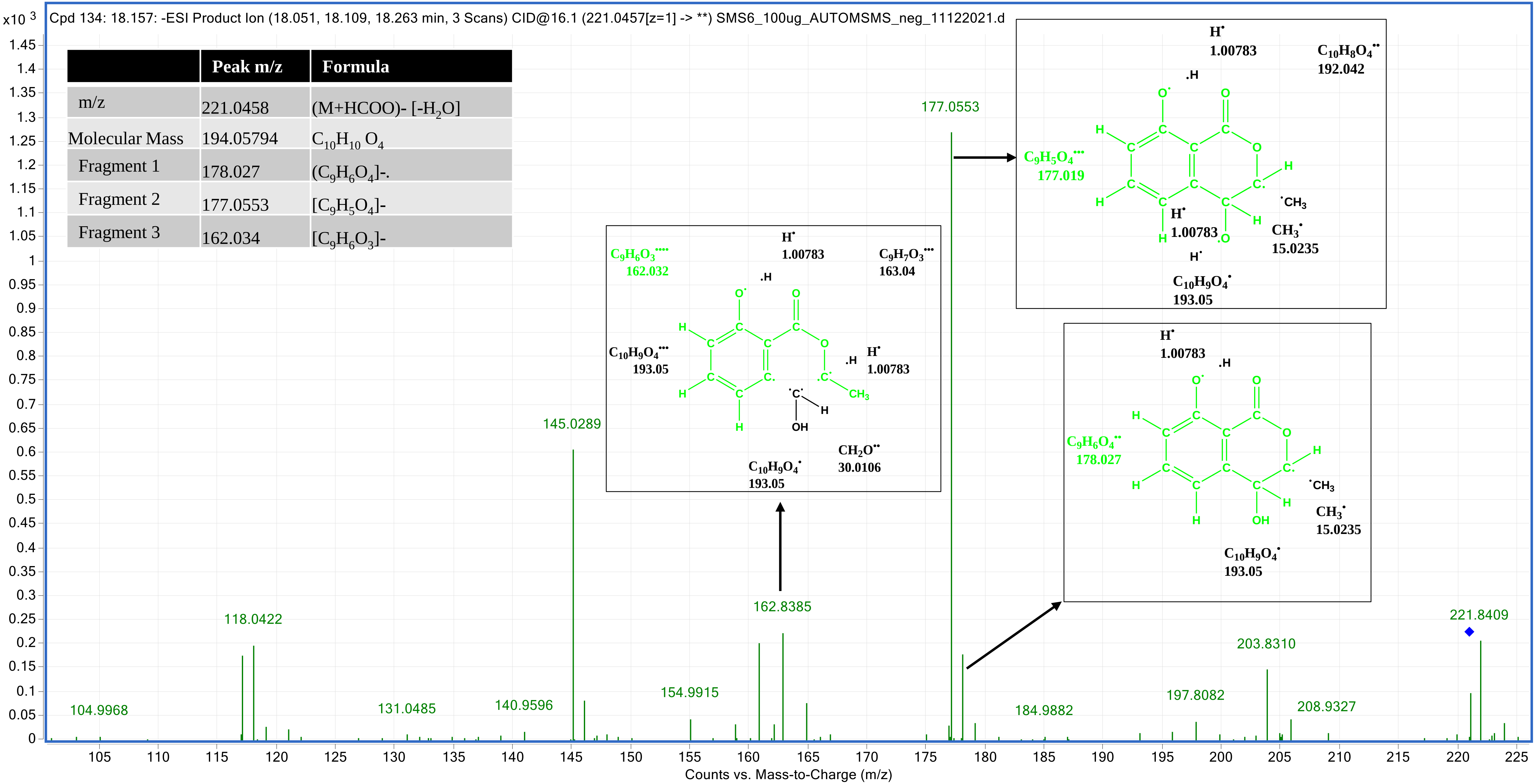

### 5'-Epialtenuene (2CQS) _16.419

| | Peak m/z | Formula |
|---|---|---|
| m/z | 291.0878 | (M-H)- |
| Molecular Mass | 292.09474 | $C_{15}H_{16}O_6$ |
| Fragment 1 | 291.0866 | $[C_{15}H_{15}O_6]$- |
| Fragment 2 | 274.8918 | $[C_{15}H_{14}O_5]$- |
| Fragment 3 | 230.0569 | $[C_{14}H_{14}O_3]$- |
| Fragment 4 | 203.0343 | $[C_{12}H_{11}O_3]$- |

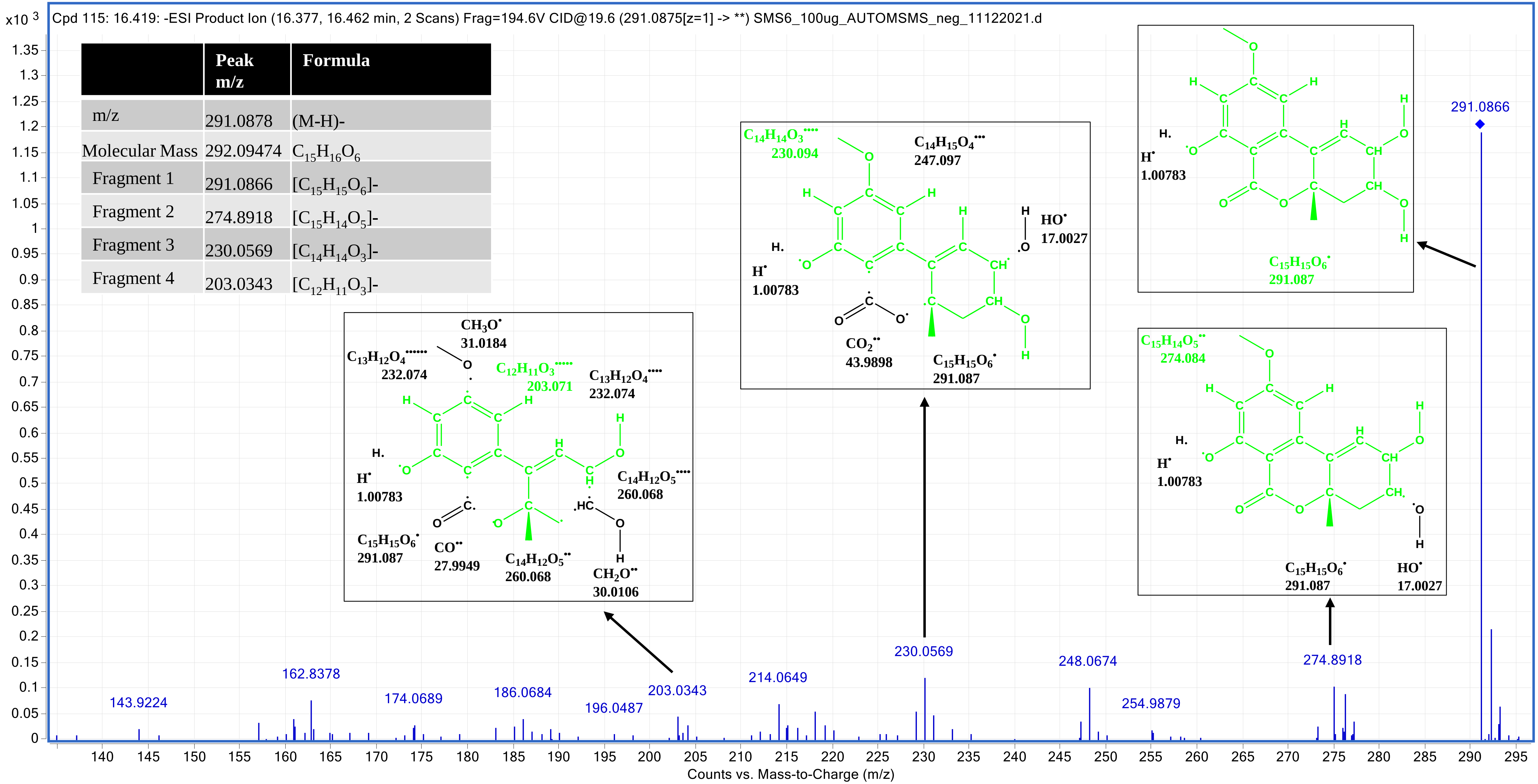

### Alternarienonic acid (3CQS) _14.99

| | Peak m/z | Formula |
|---|---|---|
| m/z | 277.07908 | (M-H)- |
| Molecular Mass | 278.0792 | $C_{14}H_{14}O_6$ |
| Fragment 1 | 277.0716 | $[C_{14}H_{12}O_6]$- |
| Fragment 2 | 233.0813 | $[C_{13}H_{13}O_4]$- |
| Fragment 3 | 218.0584 | $[C_{12}H_{10}O_4]$- |
| Fragment 4 | 205.0860 | $[C_{12}H_{13}O_3]$- |
| Fragment 5 | 190.0639 | $[C_{11}H_9O_3]$- |

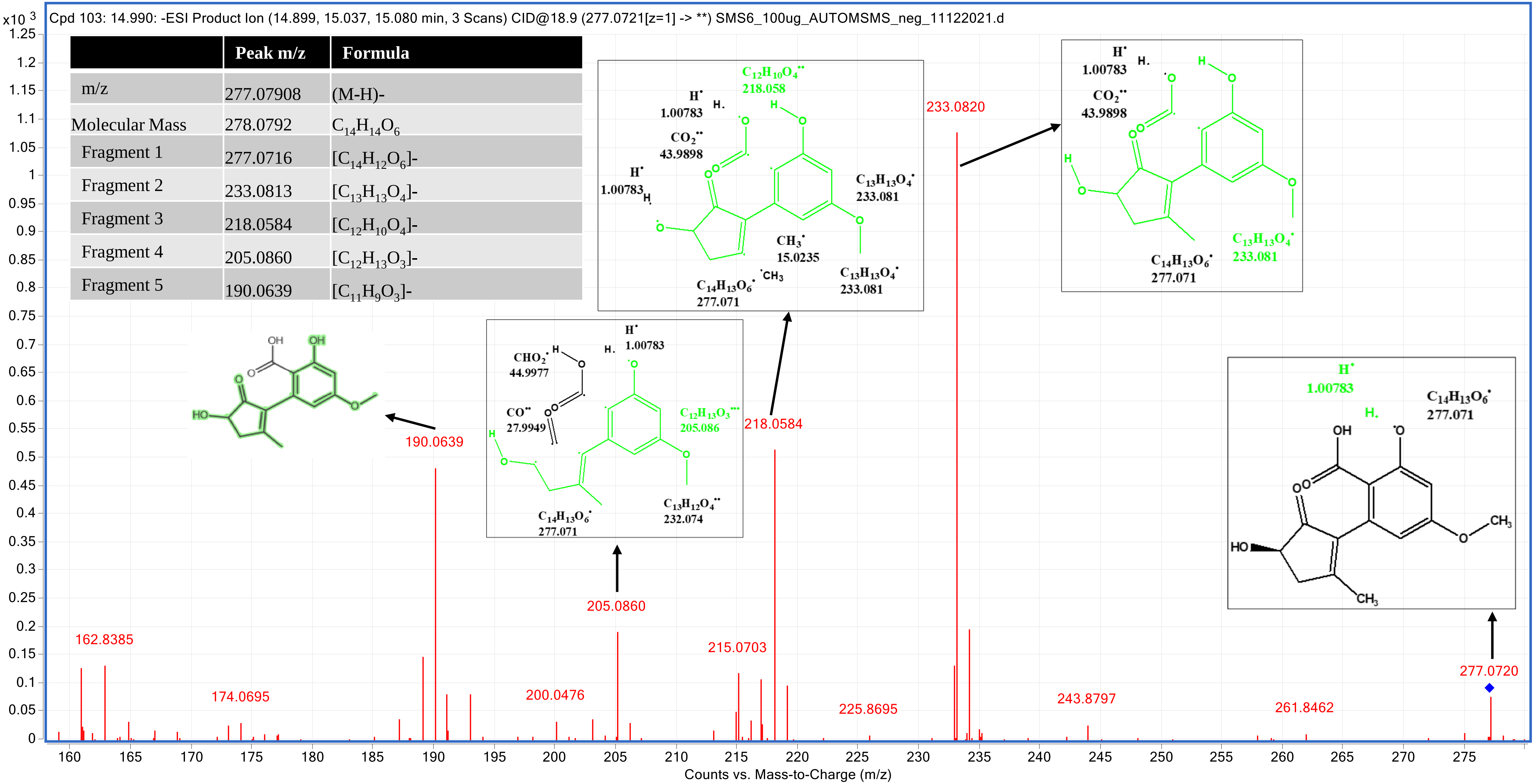

### (+)-talaroflavone (4CQS) _21.427

| | Peak m/z | Formula |
|---|---|---|
| m/z | 257.0465 | (M-H)- [-$H_2O$] |
| Molecular Mass | 276.06342 | $C_{14}H_{12}O_6$ |
| Fragment 1 | 257.0457 | [$C_{14}H_{10}O_5$]- |
| Fragment 2 | 228.9590 | [$C_{13}H_8O_4$]- |
| Fragment 3 | 215.0355 | [$C_{12}H_7O_4$]- |
| Fragment 4 | 202.063 | [$C_{12}H_{10}O_3$]- |

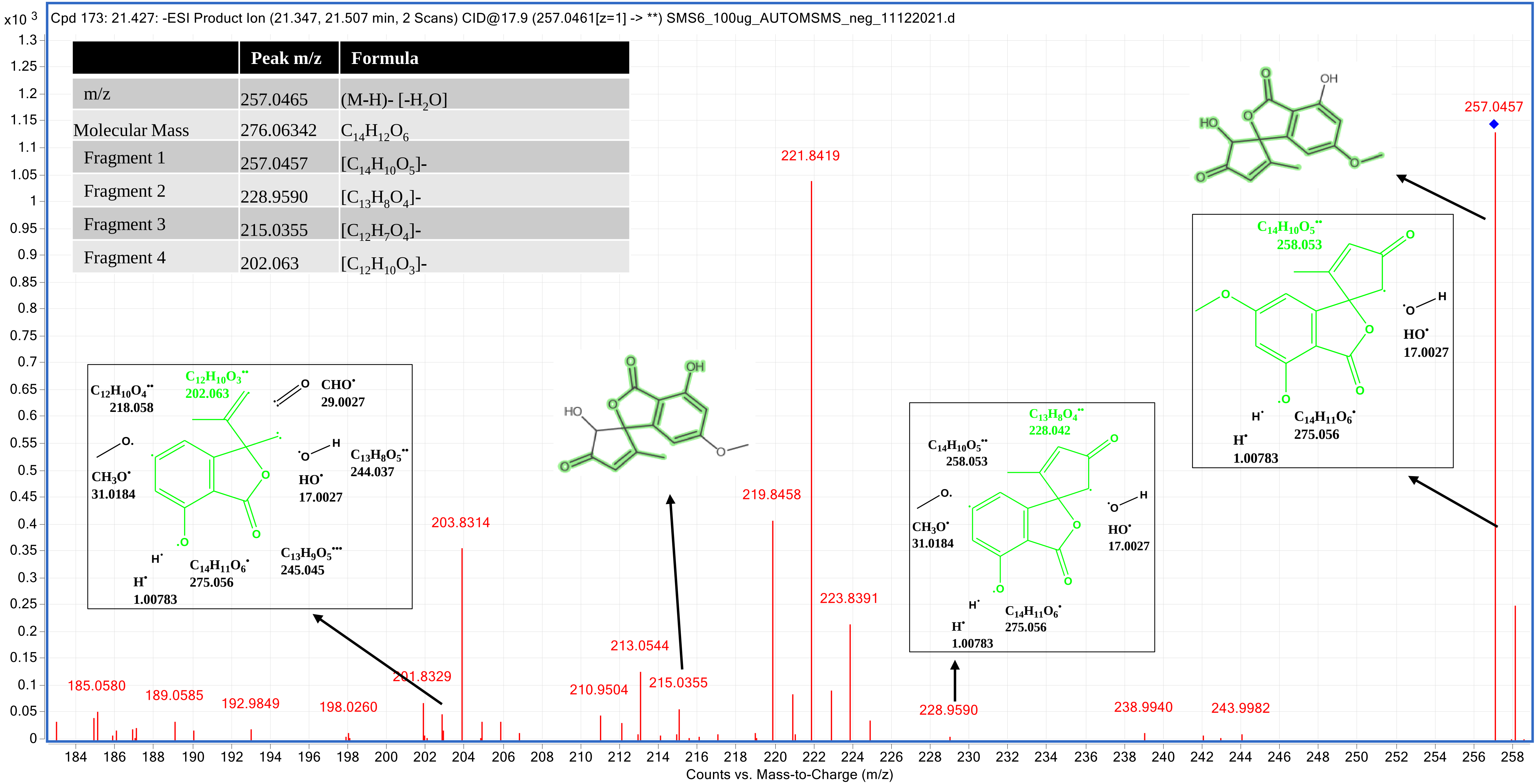

### 2,5-dimethyl-7-hydroxychromone (5CQS) _14.807

| | Peak m/z | Formula |
|---|---|---|
| m/z | 189.0563 | (M-H)- |
| Molecular Mass | 190.06303 | $C_{11}H_{10}O_3$ |
| Fragment 1 | 189.055 | $[C_{11}H_9O_3]$- |
| Fragment 2 | 174.0355 | $[C_{10}H_6O_3]$- |
| Fragment 3 | 162.8384 | $[C_{10}H_{10}O_2]$- |
| Fragment 4 | 105.0328 | $[C_{11}H_7O]$- |
| Fragment 5 | 146.0358 | $[C_9H_7O_2]$-H- |

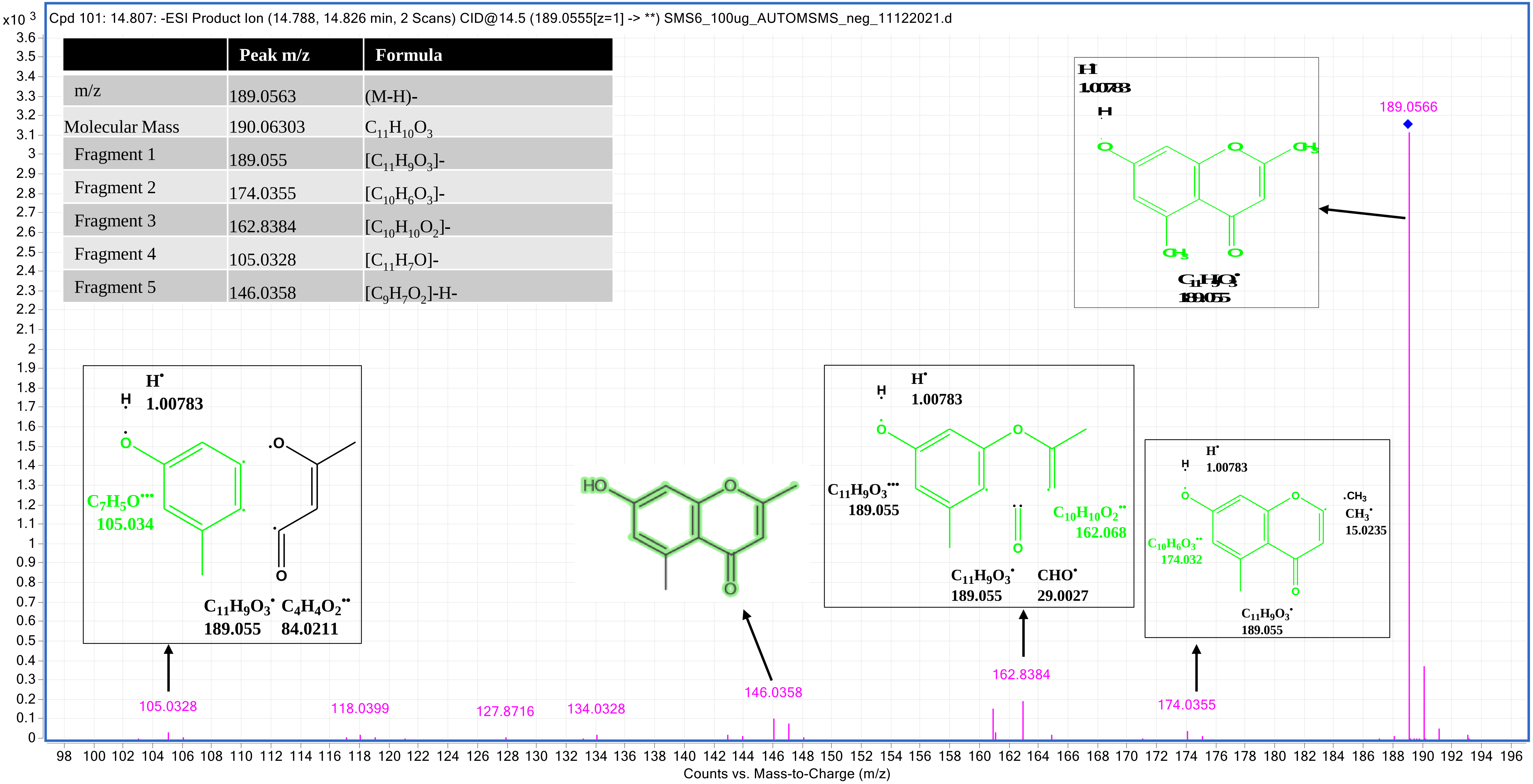

### Alternarian acid (6CQS) _15.882

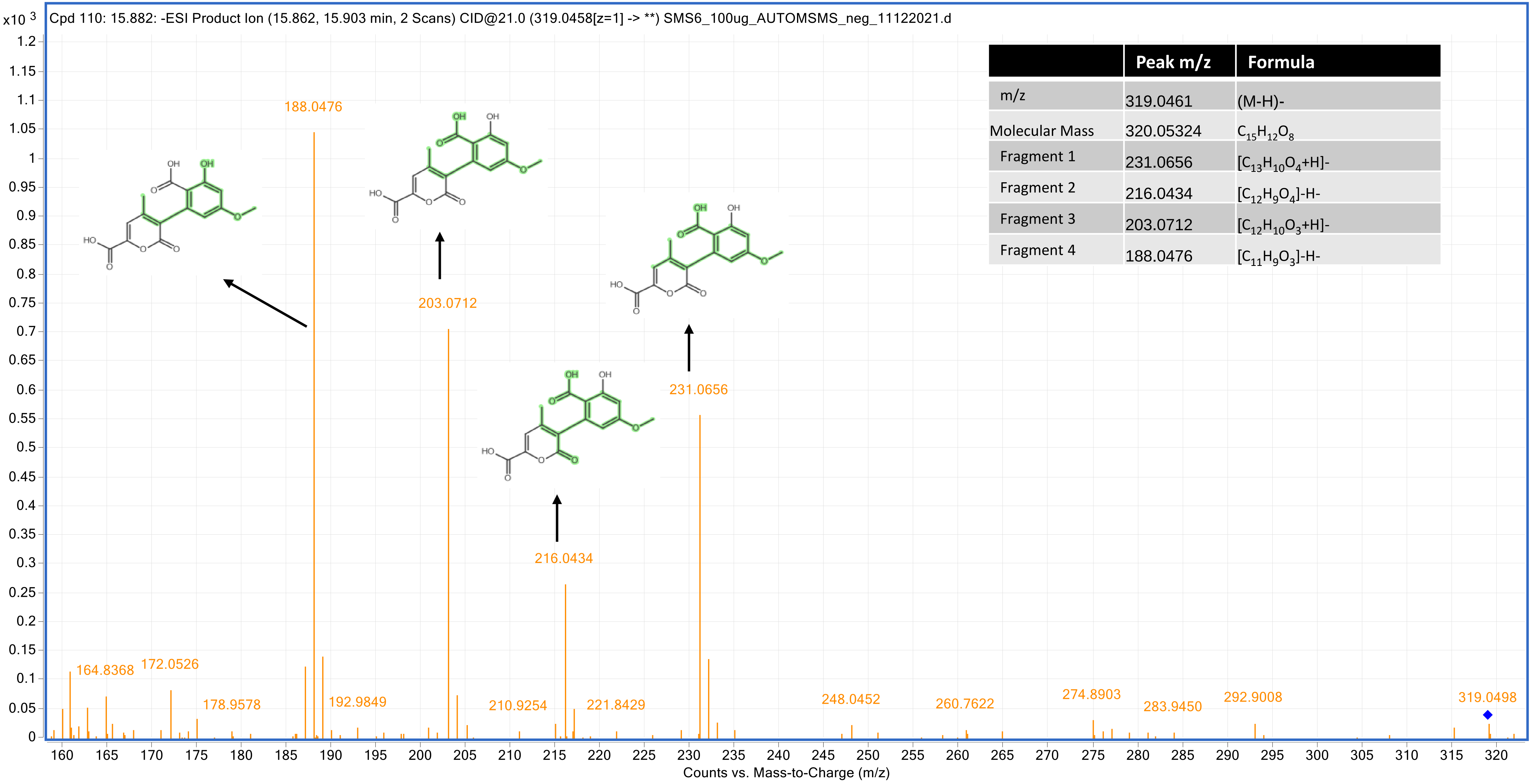


| | Peak m/z | Formula |
|---|---|---|
| m/z | 319.0461 | (M-H)- |
| Molecular Mass | 320.05324 | $C_{15}H_{12}O_8$ |
| Fragment 1 | 231.0656 | $[C_{13}H_{10}O_4+H]^-$ |
| Fragment 2 | 216.0434 | $[C_{12}H_9O_4]$-H- |
| Fragment 3 | 203.0712 | $[C_{12}H_{10}O_3+H]^-$ |
| Fragment 4 | 188.0476 | $[C_{11}H_9O_3]$-H- |

### 1,8-dihydroxynaphthalene (7CQS) _10.889

| | Peak m/z | Formula |
|---|---|---|
| m/z | 205.0509 | (M+HCOO)- |
| Molecular Mass | 160.05246 | $C_{10}H_8O_2$ |
| Fragment 1 | 143.0502 | $[C_{10}H_7O]^-$ |
| Fragment 2 | 118.9939 | $[C_8H_6O]^-$ |
| Fragment 3 | 105.0351 | $[C_7H_5O]^-$ |

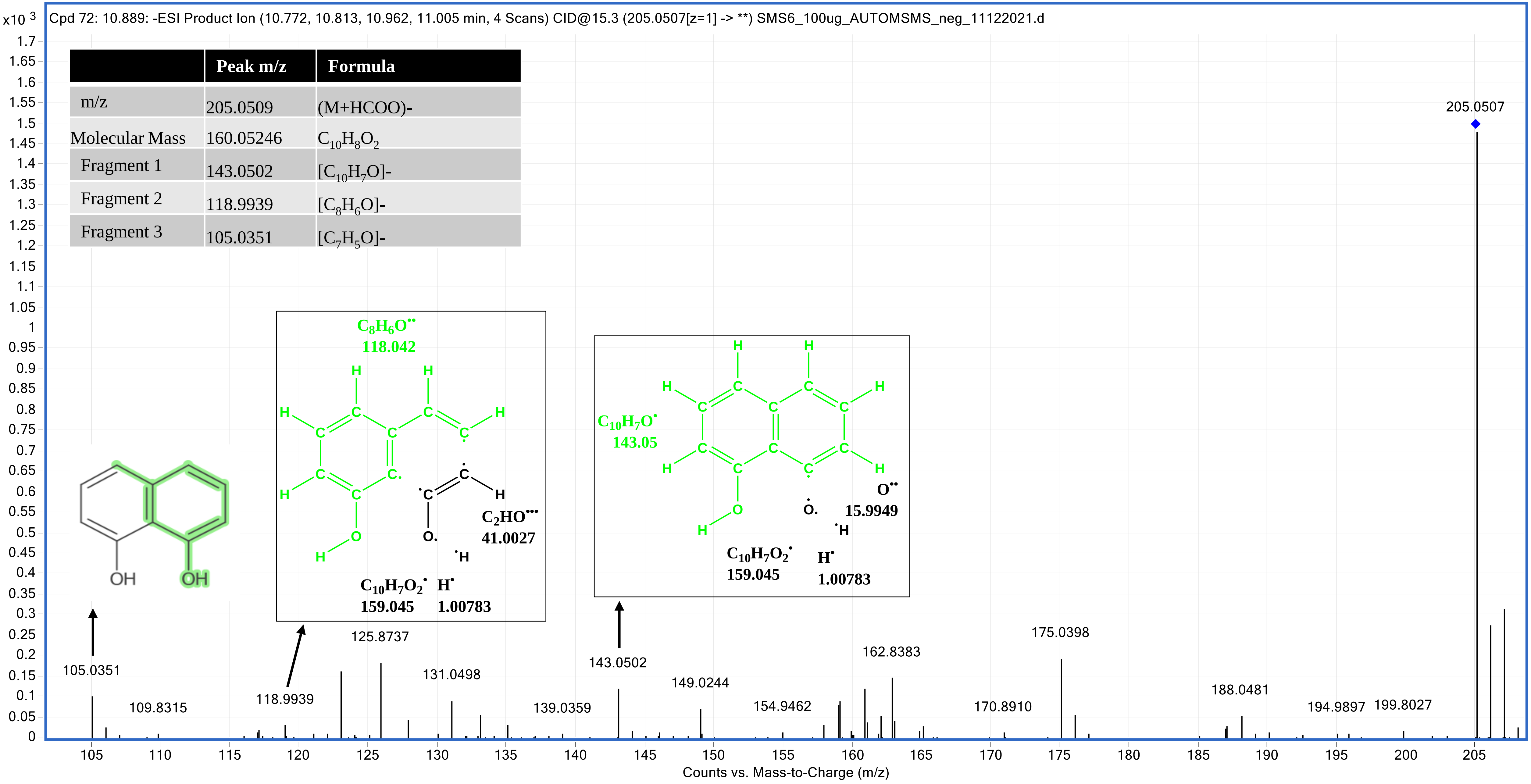

### Aspergone Q (8CQS) _17.024

| | Peak m/z | Formula |
|---|---|---|
| m/z | 223.0611 | (M-H)- [-$H_2O$] |
| Molecular Mass | 242.07908 | $C_{11}H_{14}O_6$ |
| Fragment 1 | 191.0352 | [$C_{10}H_{10}O_4$-2H]-H- |
| Fragment 2 | 149.0247 | [$C_8H_7O_3$-H]-H- |
| Fragment 3 | 123.0462 | [$C_7H_9O_2$-H]-H- |

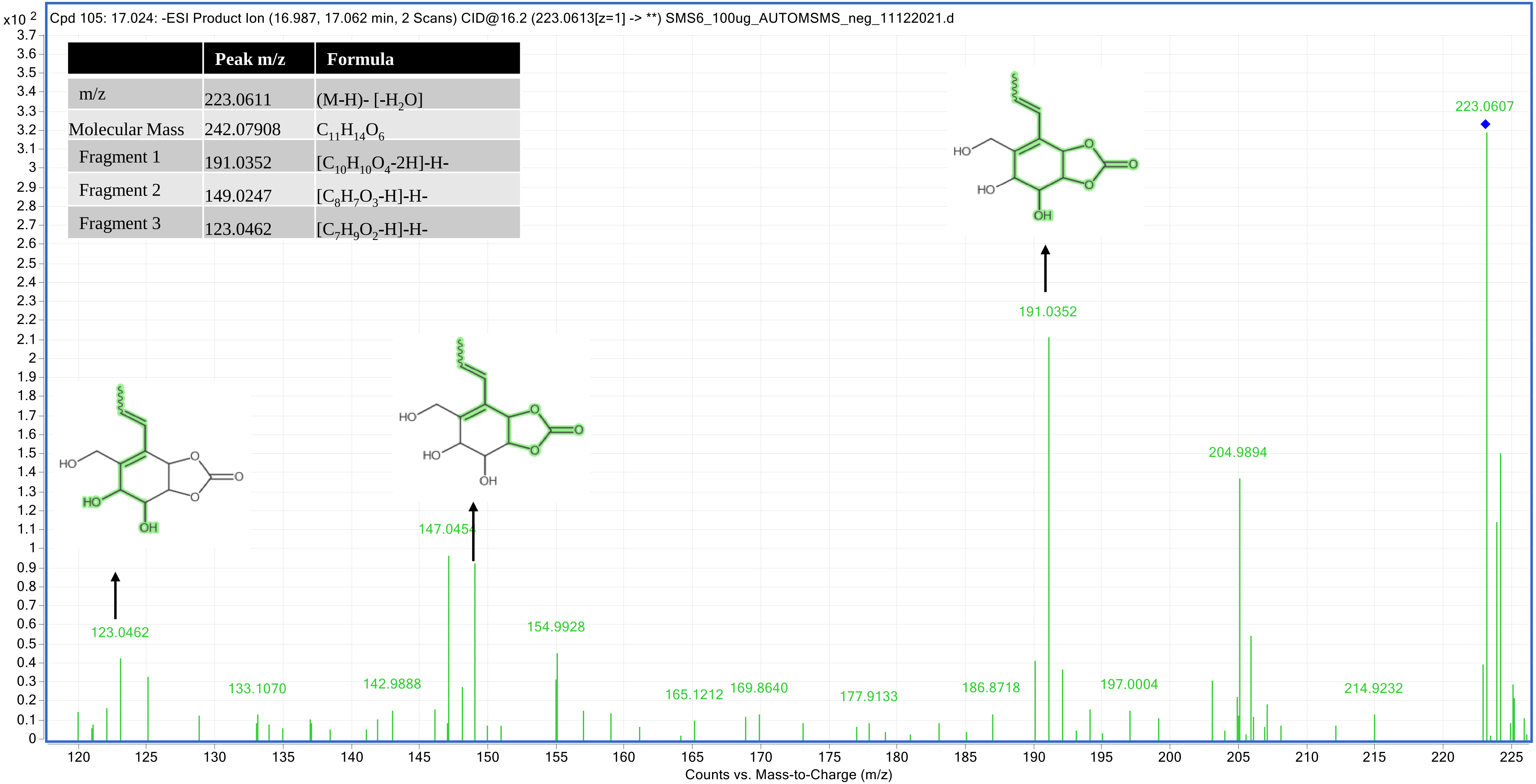

### 6-Epi-stemphytriol (9CQS) _18.361

| | Peak m/z | Formula |
|---|---|---|
| m/z | 349.0716 | (M-H)- [-$H_2$O] |
| Molecular Mass | 368.08965 | $C_{20}H_{16}O_7$ |
| Fragment 1 | 331.0592 | [$C_{20}H_{14}O_5$-2H]-H- |
| Fragment 2 | 292.061 | [$C_{14}H_{12}O_7$]- |
| Fragment 3 | 243.0665 | [$C_{14}H_{11}O_4$]- |

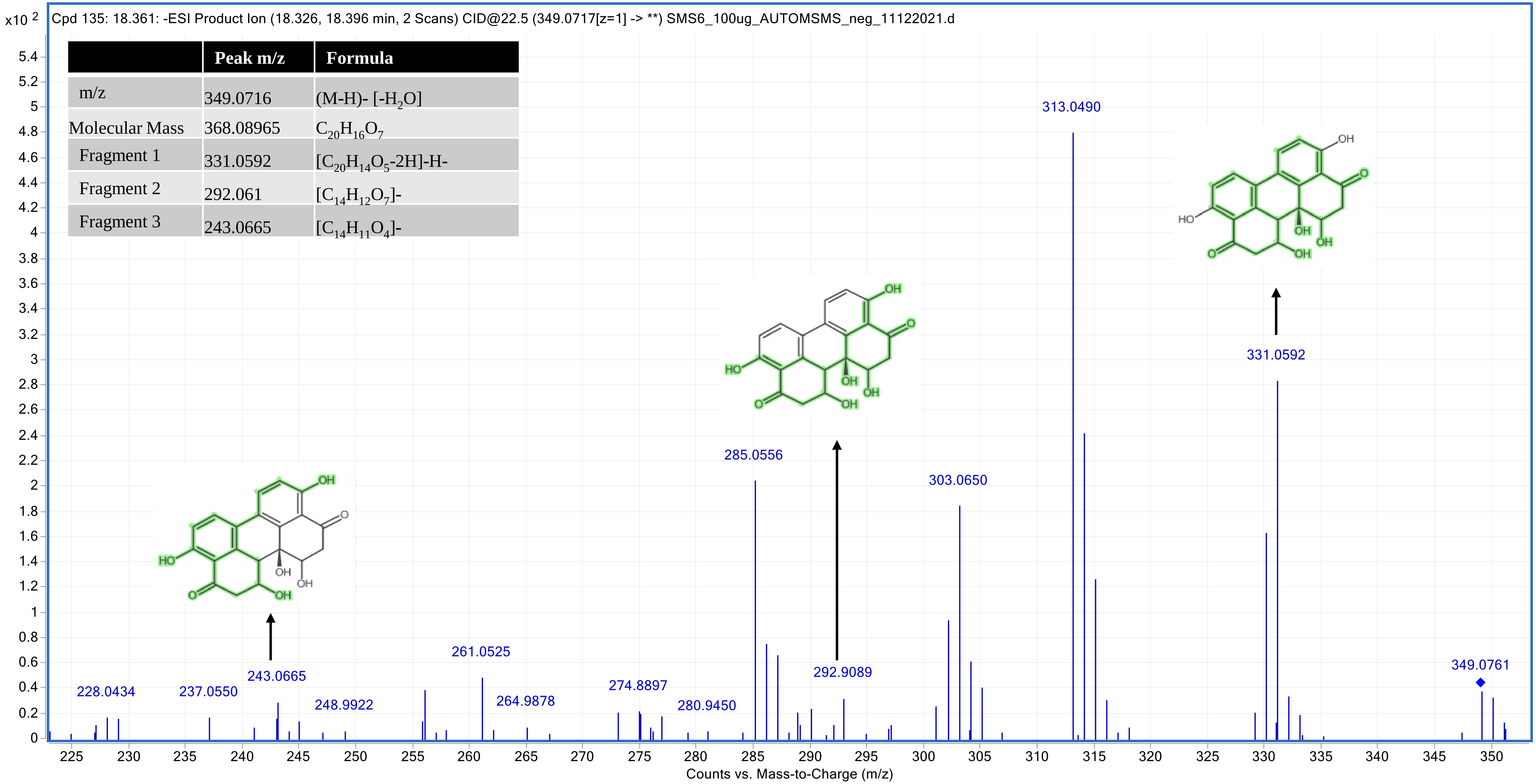

### 12-Methoxycitromycin (10CQS) _18.456

| | Peak m/z | Formula |
|---|---|---|
| m/z | 287.0568 | (M+HCOO)- |
| Molecular Mass | 260.06851 | $C_{14}H_{12}O_5$ |
| Fragment 1 | 229.05 | $[C_{13}H_9O_4]^-$ |
| Fragment 2 | 228.0424 | $[C_{13}H_8O_4]^-$ |
| Fragment 3 | 200.0467 | $[C_{12}H_8O_3]^-$ |
| Fragment 4 | 211.0386 | $[C_{13}H_8O_3]$-H- |

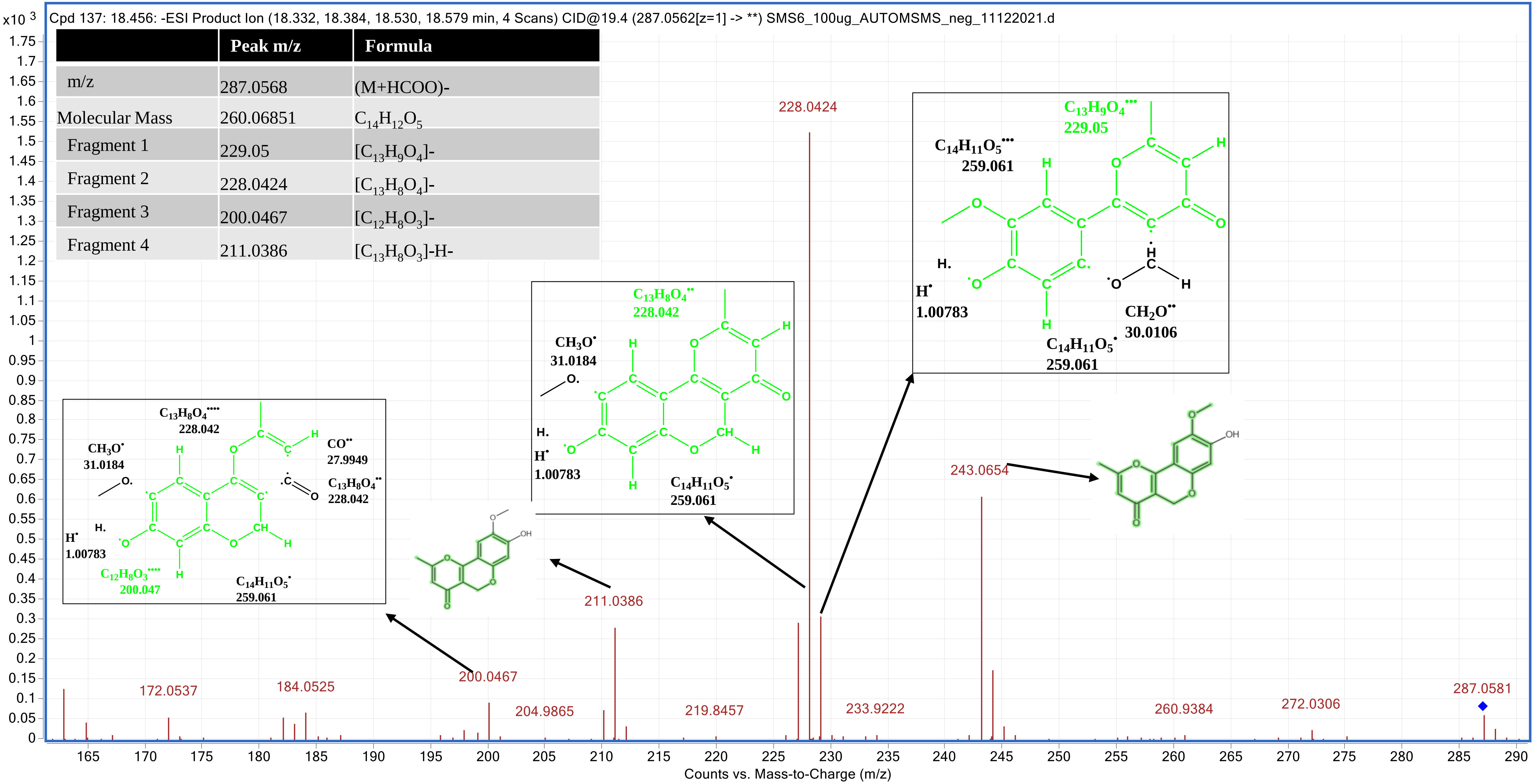

### 4-Hydroxyalternariol 9-methyl ether (11CQS) _23.997

| | Peak m/z | Formula |
|---|---|---|
| m/z | 287.0571 | (M-H)- |
| Molecular Mass | 288.06342 | $C_{15}H_{12}O_6$ |
| Fragment 1 | 287.0571 | $[C_{15}H_{11}O_6]$- |
| Fragment 2 | 272.0331 | $[C_{14}H_8O_6]$- |
| Fragment 3 | 257.0181 | $[C_{13}H_5O_6]$- |
| Fragment 4 | 228.0422 | $[C_{14}H_{12}O_3]$- |
| Fragment 5 | 188.047 | $[C_{11}H_9O_3]$-H- |

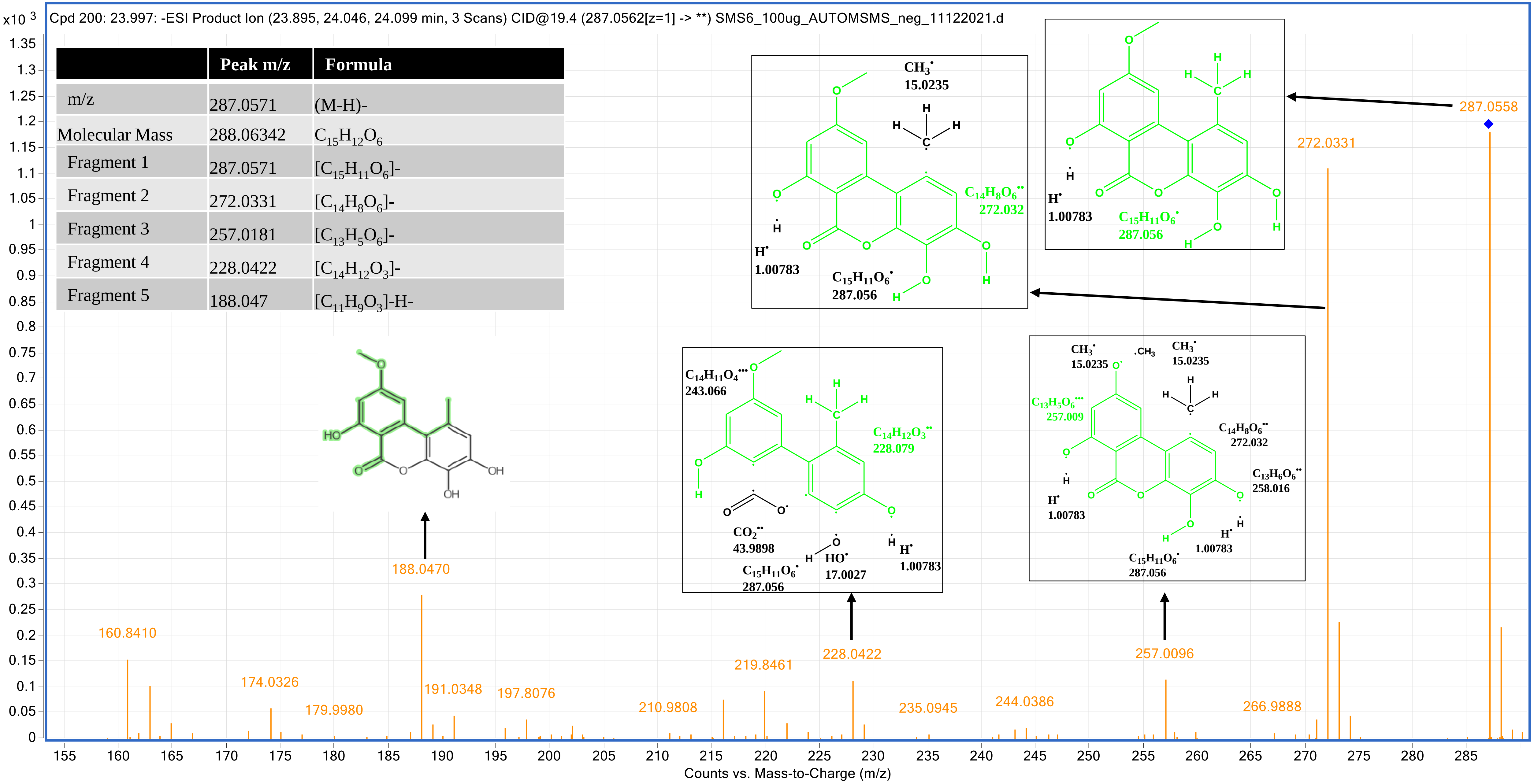

### Orthosporin (12CQS) _20.349

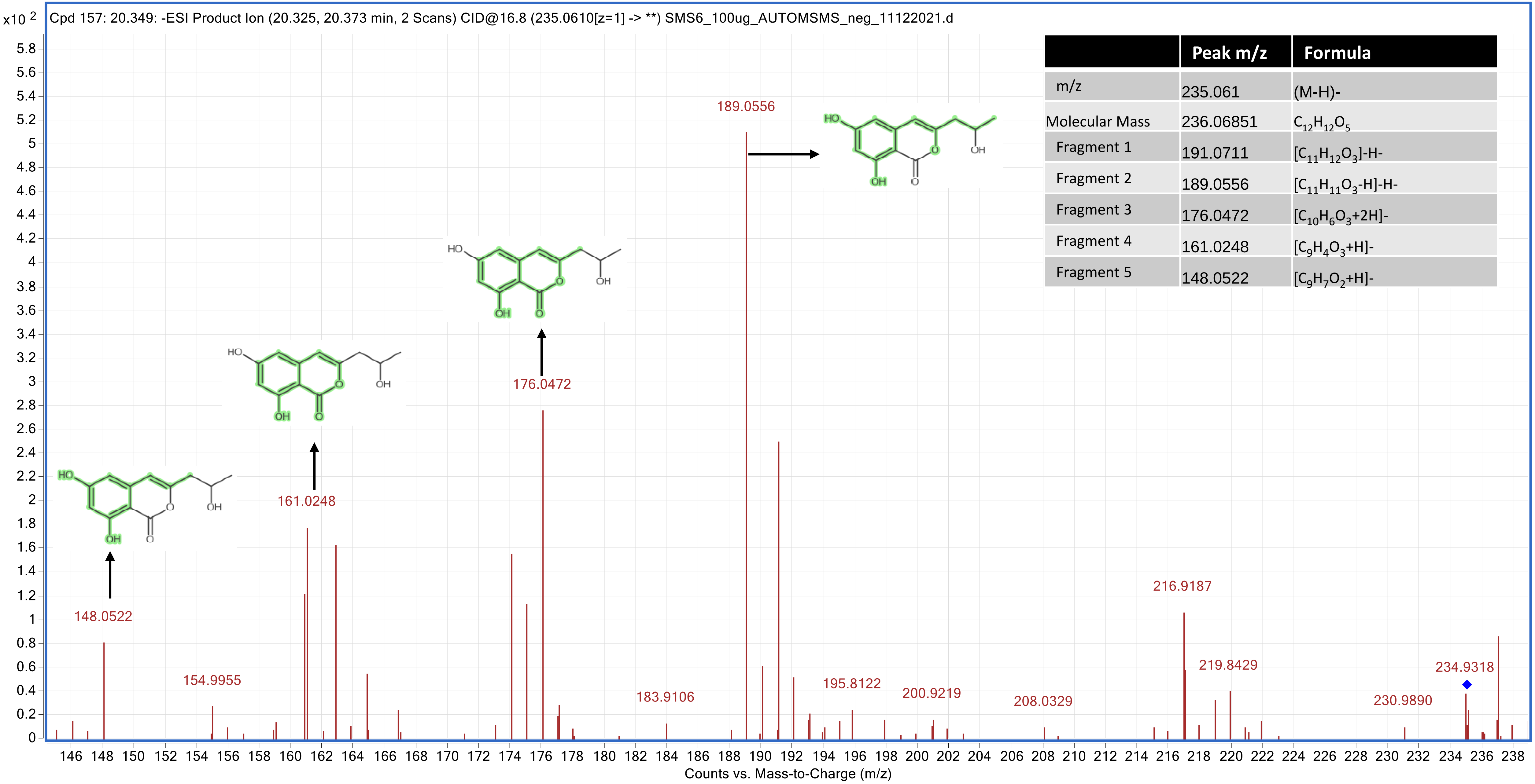


| | Peak m/z | Formula |
|---|---|---|
| m/z | 235.061 | (M-H)- |
| Molecular Mass | 236.06851 | $C_{12}H_{12}O_5$ |
| Fragment 1 | 191.0711 | $[C_{11}H_{12}O_3]$-H- |
| Fragment 2 | 189.0556 | $[C_{11}H_{11}O_3$-H]-H- |
| Fragment 3 | 176.0472 | $[C_{10}H_6O_3$+2H]- |
| Fragment 4 | 161.0248 | $[C_9H_4O_3$+H]- |
| Fragment 5 | 148.0522 | $[C_9H_7O_2$+H]- |

### Altenusin (13CQS) _22.848

| | Peak m/z | Formula |
|---|---|---|
| m/z | 317.0666 | (M+HCOO)- [-$H_2O$] |
| Molecular Mass | 290.07908 | $C_{15}H_{14}O_6$ |
| Fragment 1 | 271.019 | [$C_{15}H_{11}O_5$]- |
| Fragment 2 | 270.169 | [$C_{15}H_{10}O_5$]- |
| Fragment 3 | 242.0206 | [$C_{14}H_{10}O_4$]- |
| Fragment 4 | 214.0267 | [$C_{13}H_{10}O_3$]- |
| Fragment 5 | 198.0313 | [$C_{13}H_{10}O_2$]- |

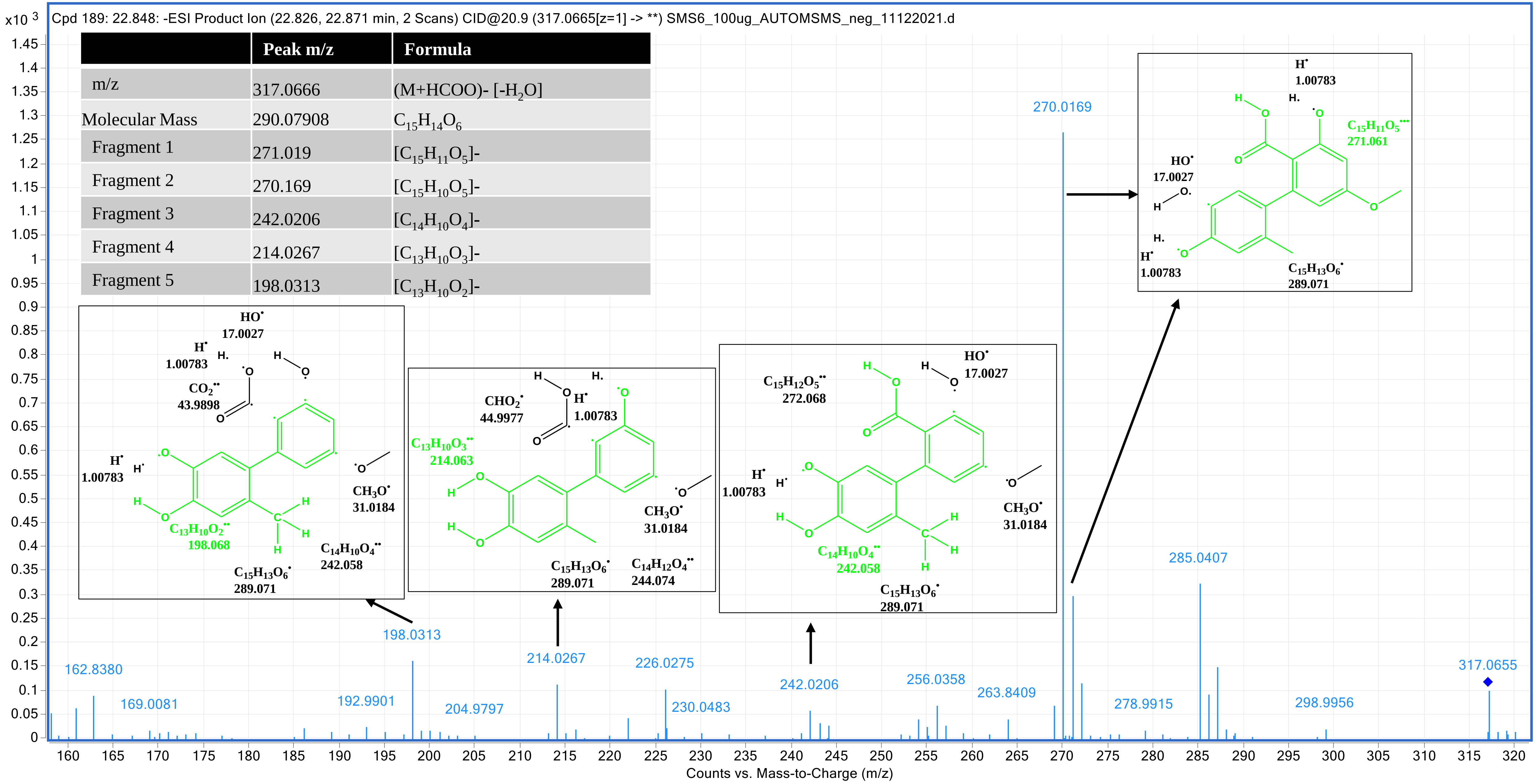
