## Supplementry Material for "Elucidation of α-glucosidase inhibitory activity and UHPLC-ESI-QTOF-MS based metabolic profiling of endophytic fungi *Alternaria* sp. BRN05. isolated from seeds of *Swietenia macrophylla* King": Supplementry Material 3.pdf

| S. No | Compound Name | Molecular Formula | Pose-1 Binding energy (kcal/mol) | RMSD |
|---|---|---|---|---|
| | **Molecules common to The EFS and EQS** | | | |
| 1CFS & 1CQS | 4-hydroxymellein | $C_{10}H_{10}O_4$ | -6 | 1.168 |
| 2CFS & 2CQS | 1,8-dihydroxynaphthalene | $C_{15}H_{16}O_6$ | -6.7 | 0.093 |
| 3CFS & 3CQS | Alternarienonic acid | $C_{14}H_{14}O_6$ | -6.1 | 1.953 |
| 4CFS & 4CQS | (+)-talaroflavone | $C_{14}H_{12}O_6$ | -6.5 | 2.833 |
| 5CFS & 5CQS | 2,5-dimethyl-7-hydroxychromone | $C_{11}H_{10}O_3$ | -5.5 | 1.581 |
| 6CFS & 6CQS | Alternarian acid | $C_{15}H_{12}O_8$ | -6.5 | 1.403 |
| 7CFS & 7CQS | 5'-Epialtenuen | $C_{10}H_8O_2$ | -6.7 | 1.757 |
| 8CFS & 8CQS | Aspergone Q | $C_{11}H_{14}O_6$ | -6.4 | 1.134 |
| 9CFS & 9CQS | 6-Epi-stemphytriol | $C_{20}H_{16}O_7$ | -7.2 | 1.544 |
| 10CFS & 10CQS | 12-Methoxycitromycin | $C_{14}H_{12}O_5$ | -6.5 | 2.267 |
| 11CFS & 11CQS | 4-Hydroxyalternariol 9-methyl ether | $C_{15}H_{12}O_6$ | -6.5 | 1.62 |
| 12CFS & 12CQS | Orthosporin | $C_{12}H_{12}O_5$ | -6.5 | 1.25 |
| 13CFS & 13CQS | Altenusin | $C_{15}H_{14}O_6$ | -6.4 | 2.356 |
| | **Molecules Unique to the EFS** | | | |
| 1UFS | 4-Ethylcatechol | $C_8H_{10}O_2$ | -6.4 | 13.571 |
| 2UFS | *p*-Coumaric acid | $C_9H_8O_3$ | -6.7 | 0.563 |
| 3UFS | Diaportinol | $C_{13}H_{14}O_6$ | -7 | 1.569 |
| 4UFS | Phenylacetic acid | $C_8H_8O_2$ | -6.4 | 2.721 |
| 5UFS | Procyanidin dimer B1 | $C_{30}H_{26}O_{12}$ | -4.5 | 2.661 |
| 6UFS | Theaflavin | $C_{29}H_{24}O_{12}$ | -4.23 | 4.48 |
| | **Molecules Unique to the EFS** | | | |

| | | | | |
|---|---|---|---|---|
| 1UQS | 6-O-desmethylterphenyllin | $C_{19}H_{16}O_5$ | -6.9 | 1.443 |
| 2UQS | Altertoxin I | $C_{20}H_{16}O_6$ | -7.8 | 4.518 |
| 3UQS | Altechromone B | $C_{14}H_{14}O_6$ | -6.8 | 2.083 |
| 4UQS | Botryorhodine F | $C_{16}H_{14}O_6$ | -6.7 | 1.521 |
| 5UQS | 3',4',7-Trihydroxyisoflavanone | $C_{15}H_{12}O_5$ | -7.5 | 1.811 |
| 6UQS | alternariol 9-methyl ether | $C_{15}H_{12}O_5$ | -7 | 3.741 |
| 7UQS | Morin | $C_{15}H_{10}O_7$ | -7.5 | 2.805 |

Supplementary file 3 (A): Docking studies were performed for twenty-six tentatively identified molecules from EFS and EQS using AutoDock Vina software with α-glucosidase (2QMJ).

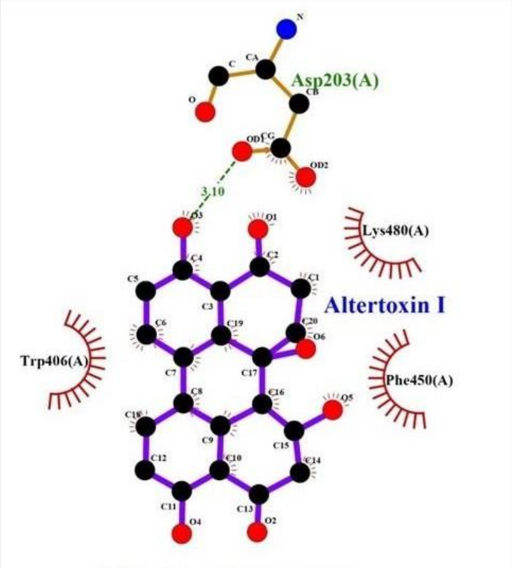


2QMJ_Altertoxin I

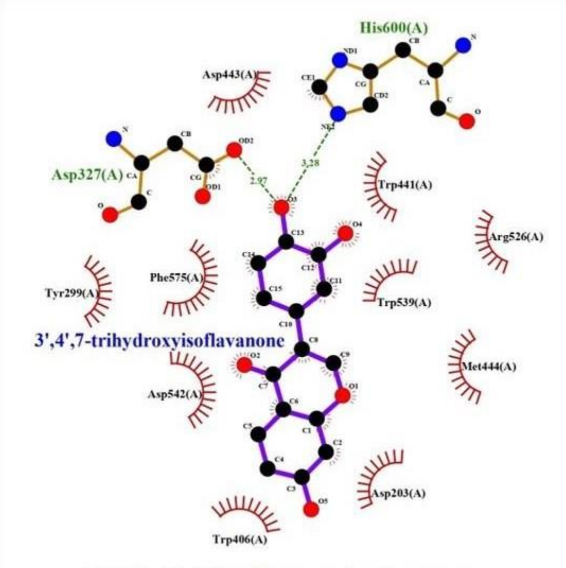


2QMJ_3',4',7-trihydroxyisoflavanone

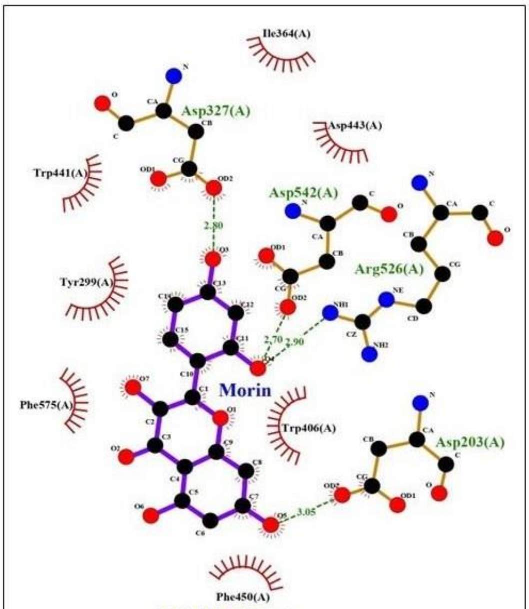


2QMJ_Morin

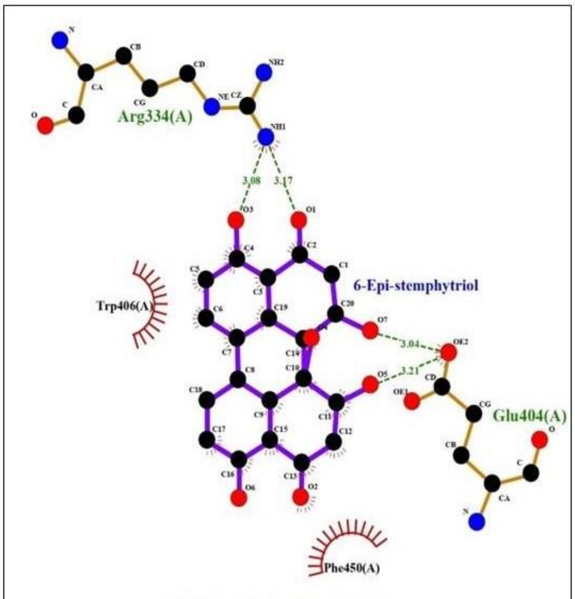


2QMJ_6-Epi-stemphytriol

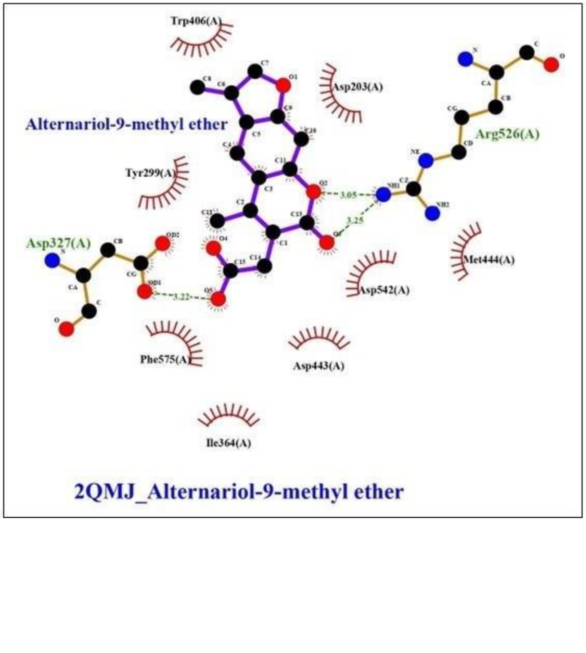


2QMJ_Alternariol-9-methyl ether

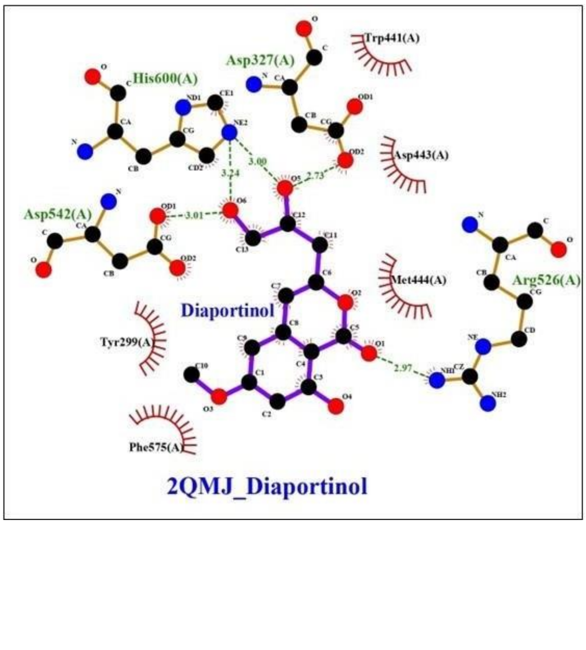


2QMJ_Diaportinol

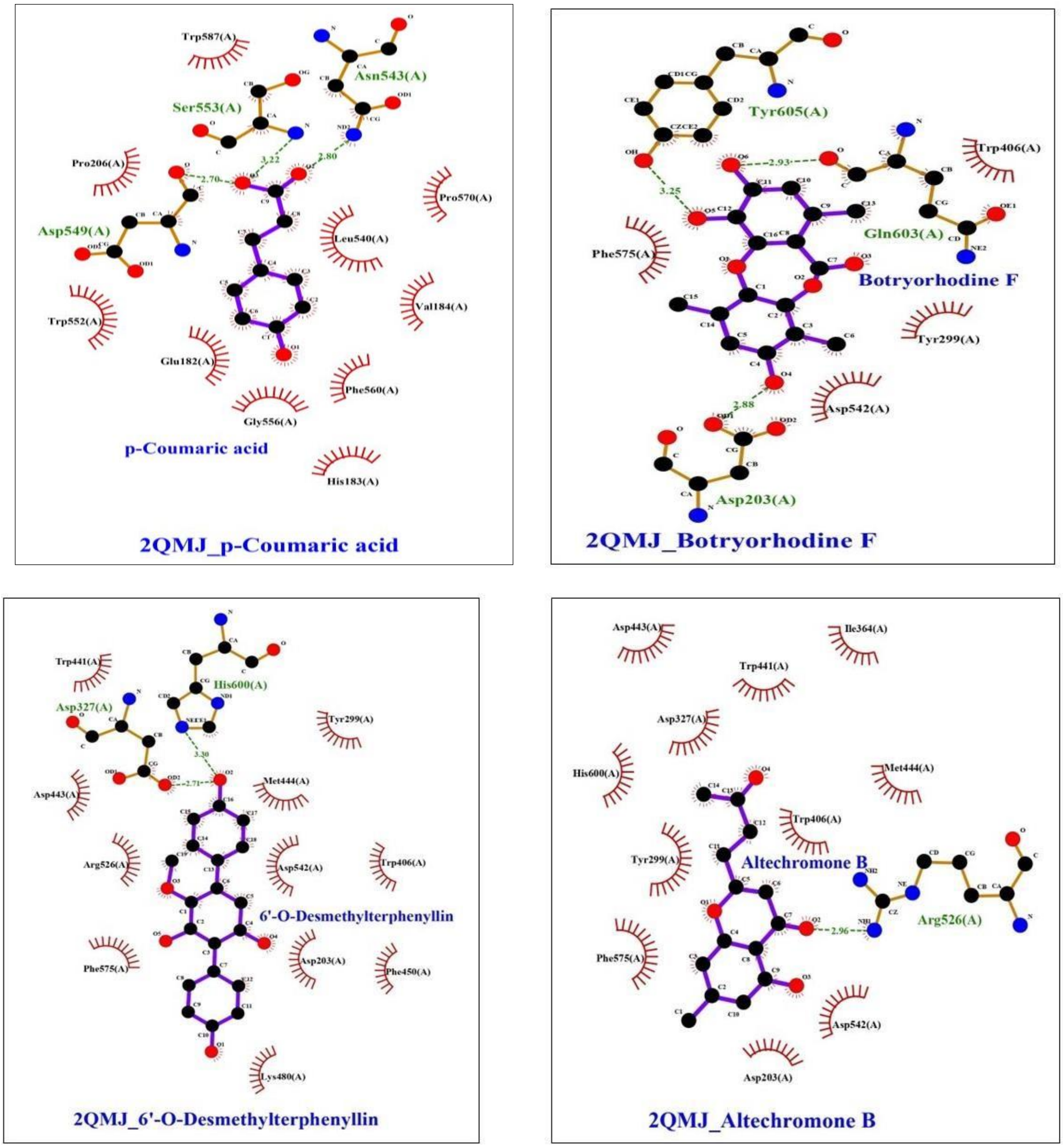

Trp587(A)
Asn543(A)
Ser553(A)
Pro206(A)
Pro570(A)
Asp549(A)
Leu540(A)
Val184(A)
Trp552(A)
Glu182(A)
Phe560(A)
Gly556(A)
p-Coumaric acid
His183(A)
2QMJ_p-Coumaric acid
Tyr605(A)
Trp406(A)
Gln603(A)
Phe575(A)
Botryorhodine F
Tyr299(A)
Asp542(A)
Asp203(A)
2QMJ_Botryorhodine F
Trp441(A)
His600(A)
Asp327(A)
Tyr299(A)
Asp443(A)
Met444(A)
Arg526(A)
Asp542(A)
Trp406(A)
6'-O-Desmethylterphenyllin
Phe575(A)
Asp203(A)
Phe450(A)
Lys480(A)
2QMJ_6'-O-Desmethylterphenyllin
Asp443(A)
Ile364(A)
Trp441(A)
Asp327(A)
His600(A)
Met444(A)
Trp406(A)
Tyr299(A)
Altechromone B
Arg526(A)
Phe575(A)
Asp542(A)
Asp203(A)
2QMJ_Altechromone B

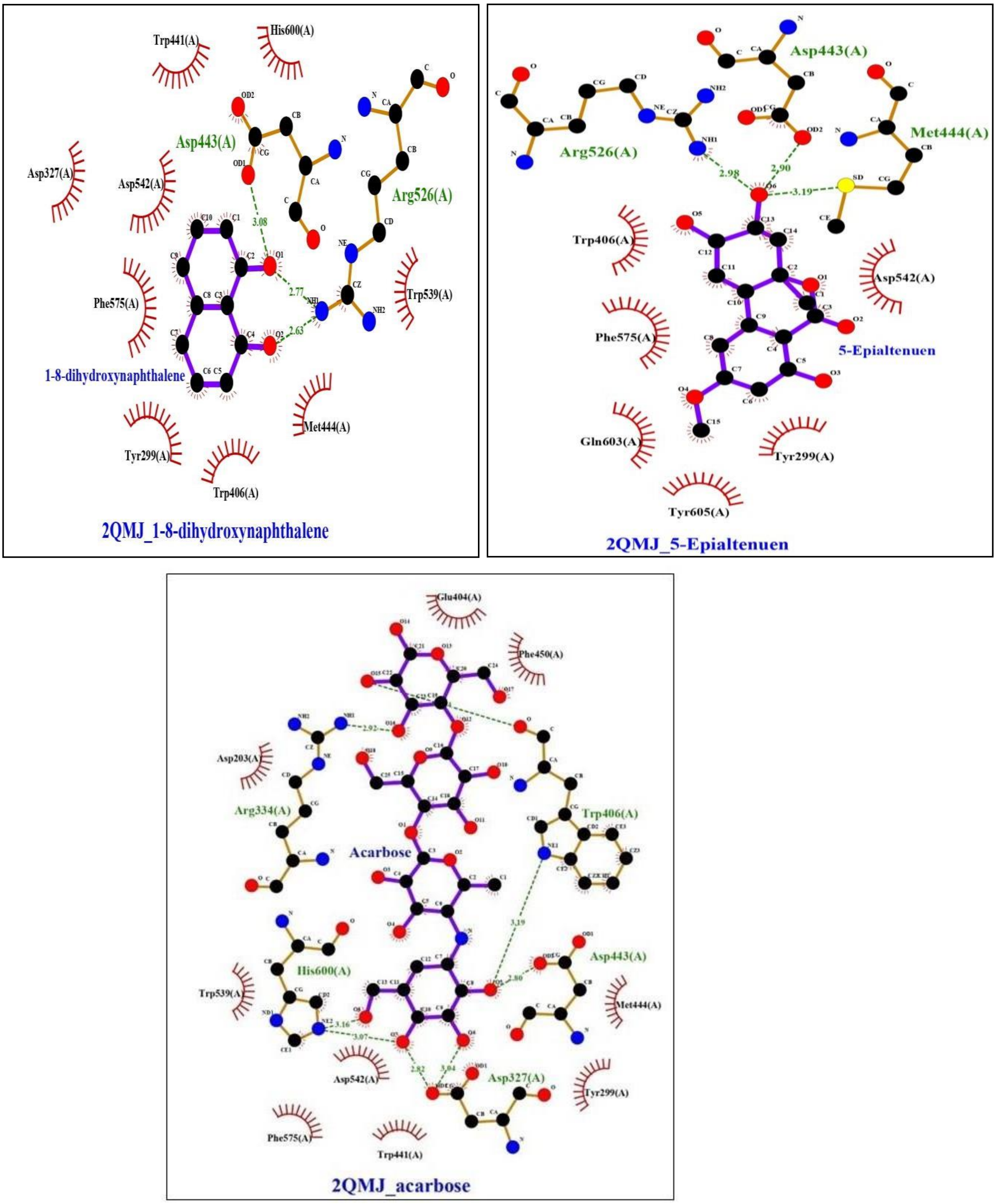


Supplementary file 3 (B). Twelve molecules identified from EFS and EQS exhibited greater binding affinities for the active site of α-glucosidase compared to acarbose.

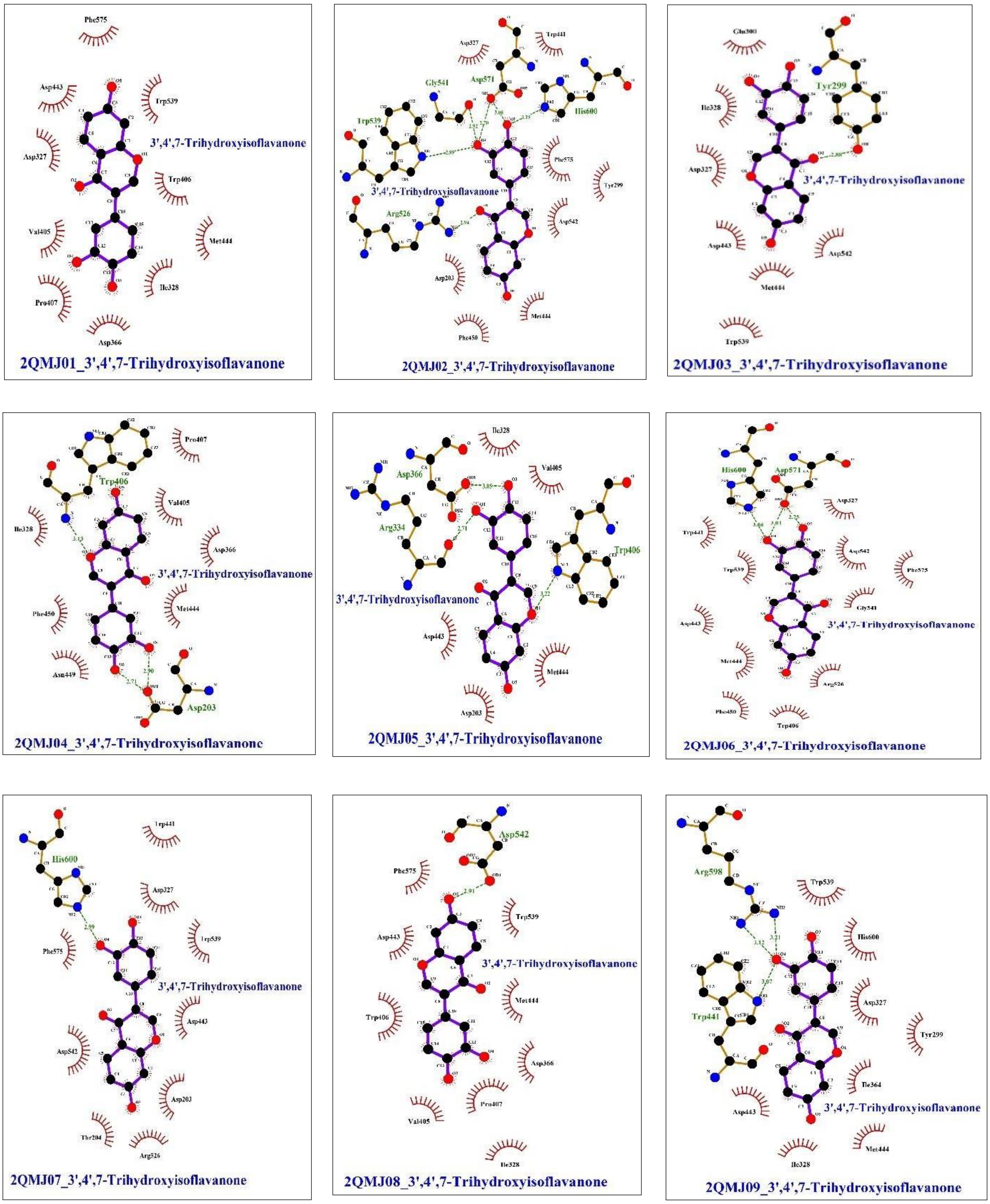


Supplementary file 3 (C). Ensemble docking of 3',4',7-Trihydroxyisoflavanone was plotted using the ligplot plus for the nine poses of a ligands bound to its 2QMJ protein.

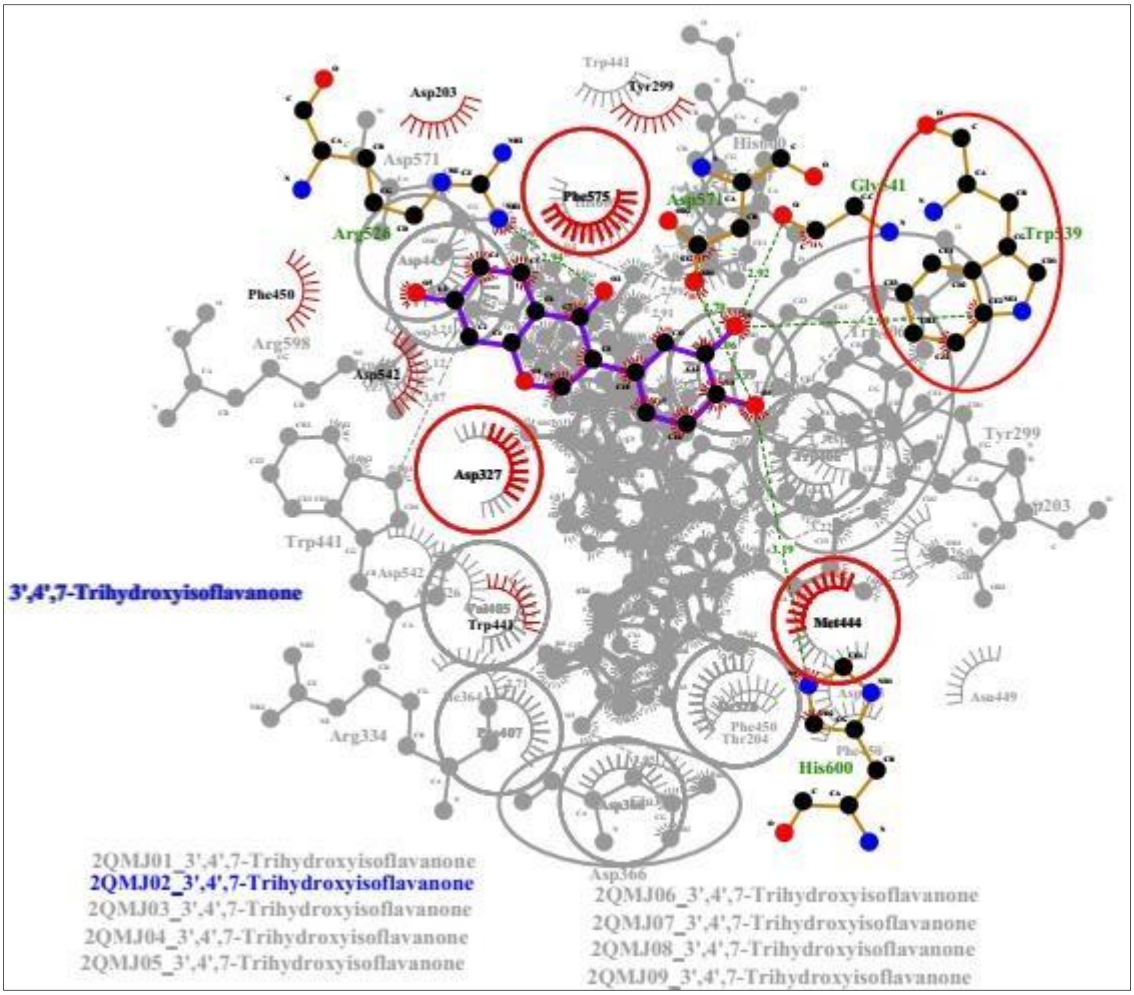


Supplementary file 3 (C). The overlapping of the docked complex of 3',4',7-Trihydroxyisoflavanone with nine poses of 2QMJ protein obtained from simulation.

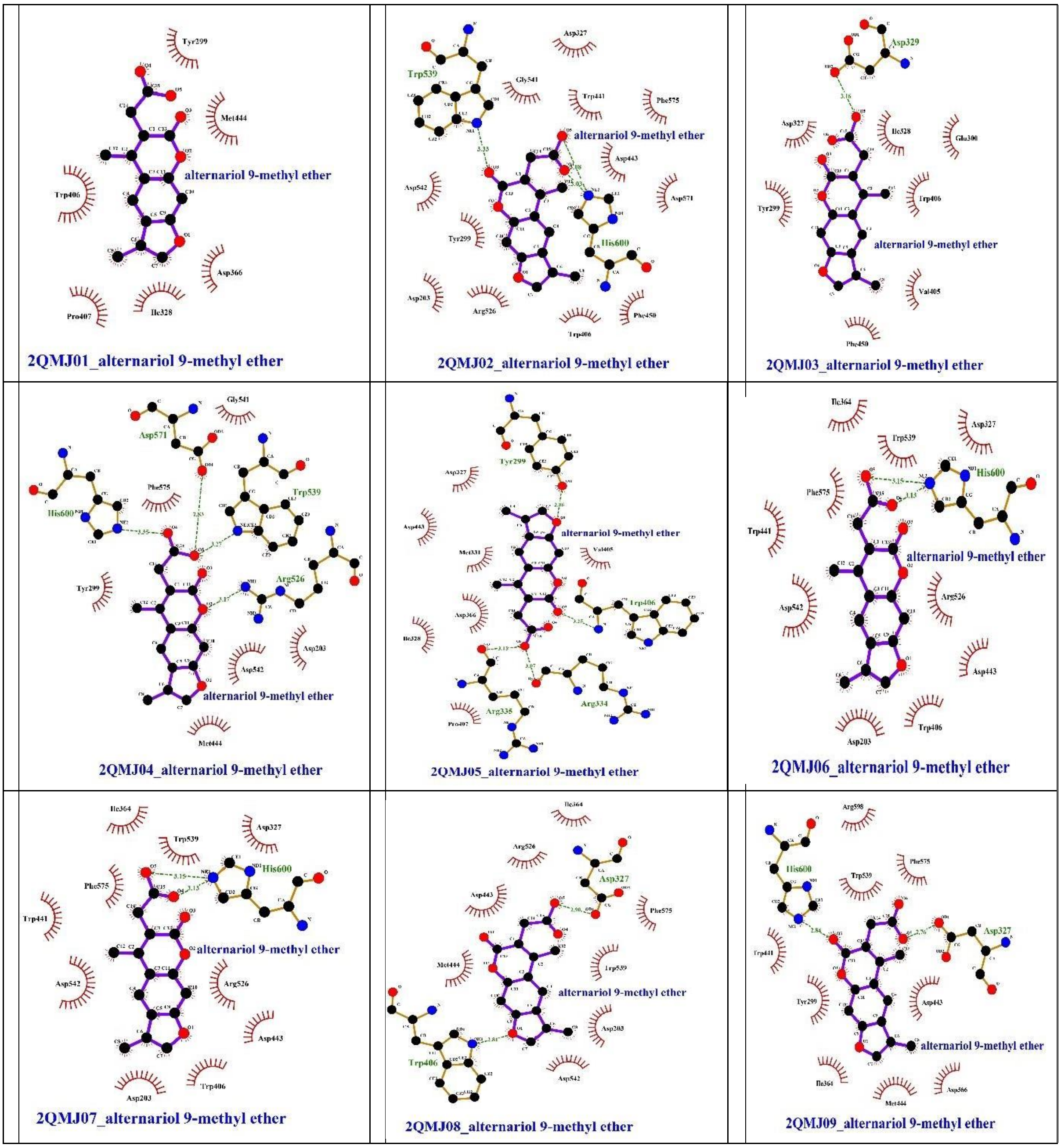


Supplementary file 3 (D). Ensemble docking of alternariol 9-methyl ether was plotted using ligplot plus for nine poses of a ligands bound to its 2QMJ protein.

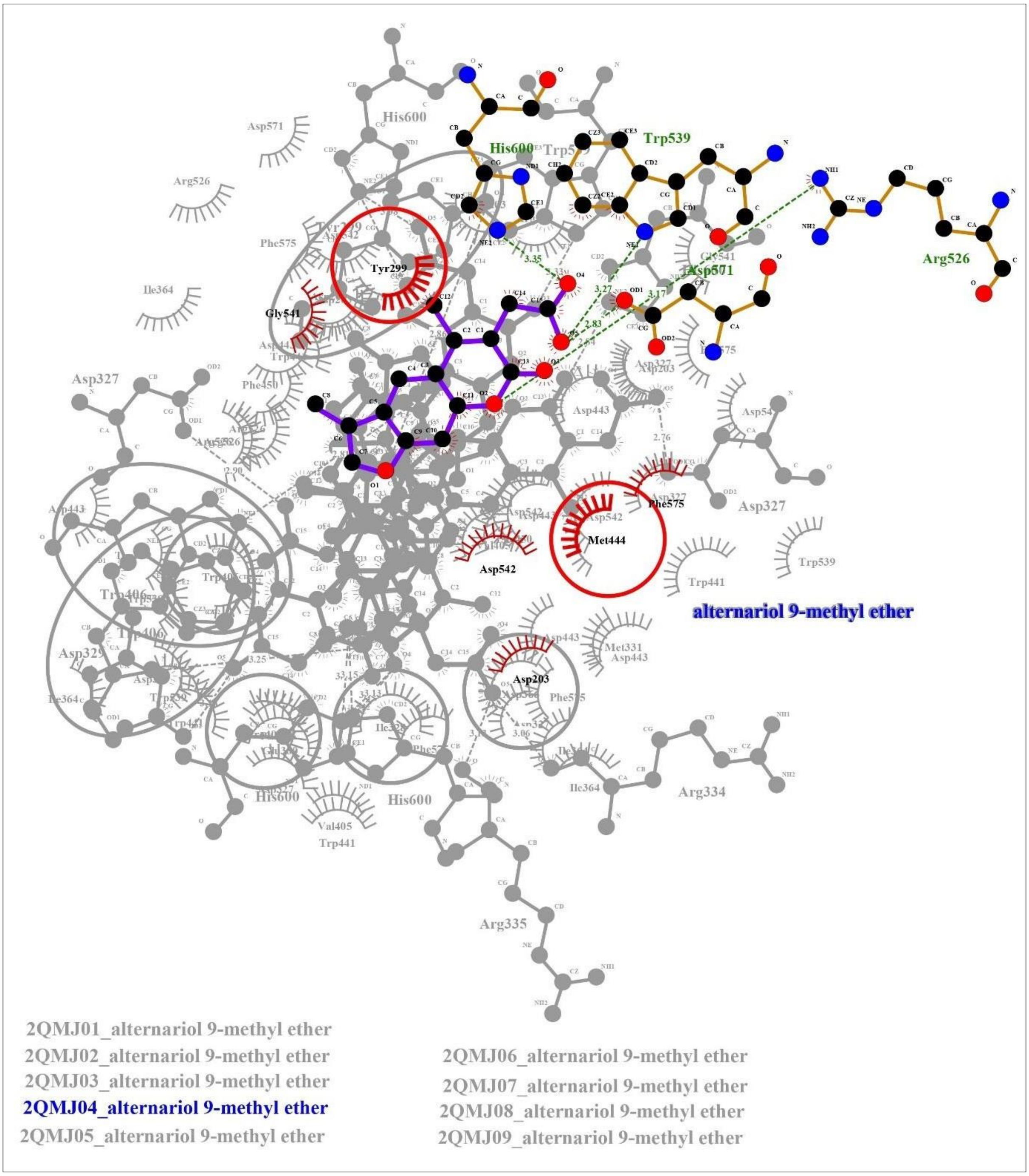


Supplementary file 3 (D). The overlapping of docked complex of alternariol 9-methyl ether with nine poses of protein obtained from simulation.

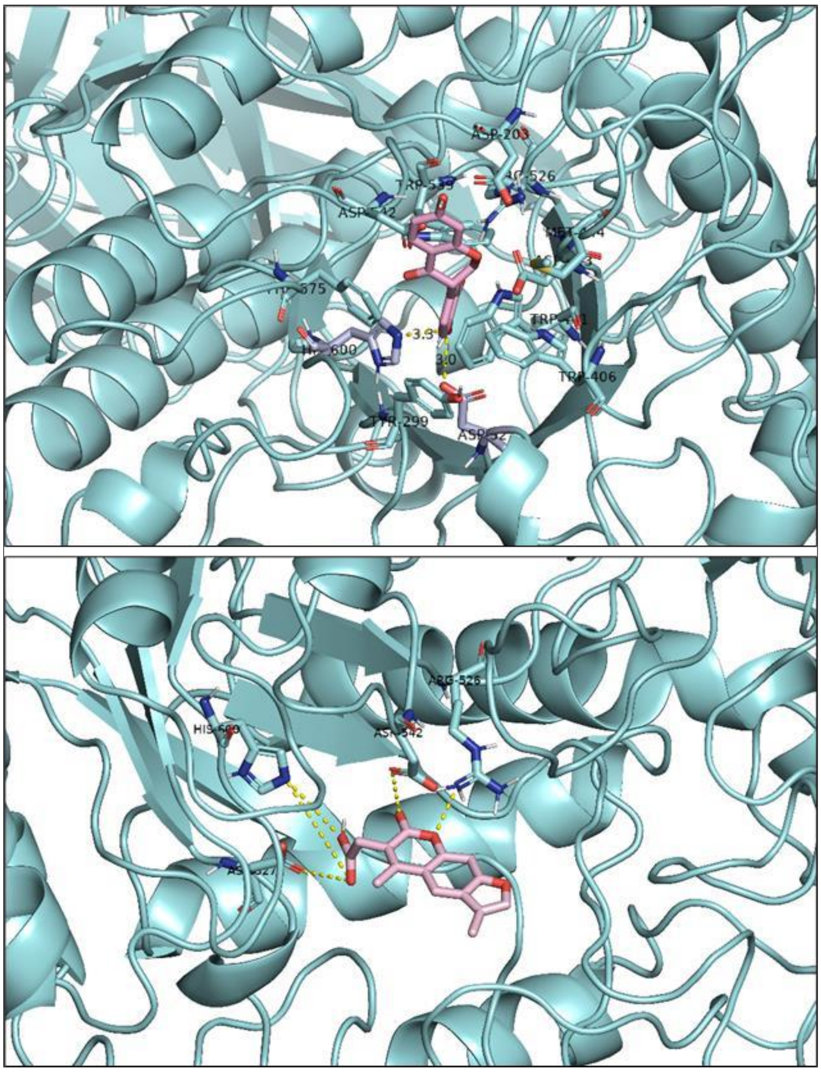


Supplementary file 3 (E). Docking of the 3',4',7-Trihydroxyisoflavanone (top) and alternariol 9-methyl ether (bottom). The figure was generated in the pymol software.
